# Plasmids modulate microindel mutations in *Acinetobacter baylyi* ADP1

**DOI:** 10.1101/2024.07.02.601687

**Authors:** Mikkel M. Liljegren, João A. Gama, Pål J. Johnsen, Klaus Harms

## Abstract

Plasmids can impact the evolution of their hosts, e.g. due to carriage of mutagenic genes, through cross-talk with host genes or as result of SOS induction during transfer. Here we demonstrate that plasmids can cause microindel mutations in the host genome. These mutations are driven by the production of single-stranded DNA molecules that invade replication forks at microhomologies and subsequently get integrated into the genome. Using the gammaproteobacterial model organism *Acinetobacter baylyi*, we show that carriage of broad host range plasmids from different incompatibility groups can cause microindel mutations directly or indirectly. The plasmid pQLICE belonging to the incompatibility group Q (IncQ) and replicating by a characteristic strand displacement mechanism can generate chromosomal microindel mutations directly with short stretches of DNA originating from pQLICE. In addition, the presence of plasmids can increase microindel mutation frequencies indirectly (i.e., with chromosomal ectopic DNA) as shown with the IncP plasmid vector pRK415 (theta replication mechanism), presumably through plasmid-chromosome interactions that lead to DNA damages. These results provide new mechanistic insights into the microindel mutation mechanism, suggesting that single-stranded DNA repair intermediates are the causing agents. By contrast, the IncN plasmid RN3 appears to suppress host microindel mutations. The suppression mechanism remains unknown. Other plasmids in this study confer ambiguous or no quantifiable mutagenic effects.

## Introduction

Plasmids are DNA molecules recognized primarily by their ability to transfer horizontally among different hosts, and in that process mobilize accessory genes that can play important roles in bacterial evolution. This is well illustrated by the plasmid-mediated spread of genes responsible for antimicrobial resistance, virulence factors and the ability to degrade xenobiotics (Heuer & Smalla, 2012). Other plasmid accessory genes may promote bacterial adaptation by enhancing mutagenesis instead, an effect that has been linked to plasmid-encoded error-prone polymerases (Woodgate & Sedgwick, 1992; Remigi et al., 2014; Sano et al., 2014; Nguyen et al., 2020). Although plasmids confer new phenotypes directly associated to the accessory genes they encode, they may also collaterally manipulate host behaviour via regulatory cross-talks, which result in the indirect expression of further phenotypic variation (Billane et al., 2021) [e.g., altered host metabolism (Hall et al., 2024)]. In addition, plasmids can also drive the movement of transposons onto the chromosome (Bergstrom et al., 2000; Hall et al., 2017) or the shuffling of integron cassettes (Baharoglu et al., 2010). The latter is a result of conjugative acquisition of plasmid DNA as single-stranded (ss) DNA molecules, which induce the bacterial SOS response. Alternatively, plasmids expressing mechanisms to prevent SOS induction (e.g., plasmidic SOS inhibition protein *psiB*) do not elicit such rearrangements (Baharoglu et al., 2010).

Plasmids comprise a wide diversity of autonomously replicating DNA molecules, and their level of autonomy depends on the replication strategy. In Gram-negative bacteria the large majority of known plasmids replicate via a uni- or bidirectional theta mechanism similar to that employed by circular chromosomes (Kim et al., 2020). Replication starts at the plasmid origin of replication, *oriV*, and proceeds from 5’ to 3’, and while the leading strand replicates continuously the lagging strand is replicated discontinuously (as Okazaki fragments). This process is regulated and initiated by plasmid-encoded factors, but involves the recruitment of parts of the host replication machinery, which are for example involved in unwinding the DNA duplex, controlling supercoiling, processing primers, or stabilizing single-stranded DNA. Some plasmids employ alternative replication mechanisms: rolling-circle and strand displacement replication. Rolling-circle replication is more commonly found among, but not restricted to, small plasmids from Gram-positive bacteria (Ruiz-Masó et al., 2015). This strategy is initiated by the plasmid replication protein Rep binding and nicking a plasmid double-strand origin of replication, that generates a free 3’-end to start leading-strand replication. Consequently, replication of the plus and minus strand is uncoupled and proceeds in two steps. As replication of the plus strand proceeds and DNA is unwound, the minus strand is displaced and covered with the single-stranded DNA binding protein. Upon termination of plus-strand synthesis, the Rep protein is inactivated to prevent sequential rounds of replication and the minus strand is released as a circular ssDNA intermediate molecule. This intermediate is then also replicated as leading strand. This step depends on the plasmid single-strand origin and on host-encoded enzymes. Lastly, the strand displacement replication mechanism is characteristic of the IncQ plasmid family (Meyer, 2009; Loftie-Eaton & Rawlings, 2011). This process is highly host-independent and responsible for the broad host range of IncQ plasmids, which besides the replication initiation protein further encode their own helicase and primase. Replication starts at *oriV*, which contains two single-strand initiation (*ssi*) sites, one in each strand. The plasmid primase recognizes each of these sites and separately initiates replication of both strands, which then will be sustained by the host DNA polymerase III. In this process both strands are replicated as leading strands simultaneously (unlike in rolling-circle replication), although synthesis from both *ssi* may not start exactly at the same time. As the replication forks progress, and the non-replicated strands are displaced, the displaced DNA surrounding the origin remains single-stranded.

Plasmid replication, despite being (semi-)autonomous, is not without consequences for the host cell, and the interaction between plasmid- and chromosome-encoded factors may lead to genetic conflicts. Sequestration of the host DNA primase DnaG by the replication-initiation protein RepA of theta-replicating plasmids can stall chromosomal replication forks, and the resulting unpaired ssDNA induces the SOS response and ultimately delays bacterial growth rates (Ingmer et al., 2001). Likewise, similar detrimental effects may result from interactions between plasmids and horizontally acquired accessory helicases (San Millan et al., 2015; Loftie-Eaton et al., 2017), which may be involved in preventing replication fork collisions that cause double-strand (ds) DNA breaks (Merrikh et al., 2012; Epshtein et al., 2014). These conflicts have however been shown to be resolved during plasmid-host co-evolution (San Millan et al., 2015; Loftie-Eaton et al., 2017; Sota et al, 2010), which may then generally improve plasmid carriage (i.e., of distinct plasmids with which the bacterial host did not evolve with). For non-theta replicating plasmids, not only the interaction with host factors, but also the accumulation of ssDNA replication intermediates, lead to genetic instability and induction of the bacterial stress response (Gigliani et al., 1993; del Solar et al., 1993; Bron et al., 1991; Zhang et al., 2019).

It was recently shown that ssDNA can be mutagenic through the Short-Patch Double Illegitimate Recombination (SPDIR) pathway (Harms et al., 2016; Liljegren et al., 2024). SPDIR mutations are caused by ssDNA from intragenomic or external sources (e.g., acquired in the course of horizontal gene transfer), and those ssDNA molecules can anneal with transiently single-stranded chromosomal DNA sections, such as lagging strands at replication forks. The annealing occurs at one or more microhomologies, i.e. short stretches of near-identical DNA sequence and may contain mismatches and gaps (extended microhomologies) in otherwise fully heterologous DNA. It is thought that these annealed ssDNA molecules are extended and eventually integrated into the nascent strand during genomic DNA replication (Fig. 1A), acting in effect as primers for Okazaki fragments (Harms et al., 2016; Liljegren et al., 2024). The resulting mutations are nucleotide polymorphism clusters or microindels of highly variable size and sequence, and the mutation patterns can be traced back to the DNA sequence of the templating single-strand (Fig. 1B, C; Supplemental file S1).

**Fig. 1.**
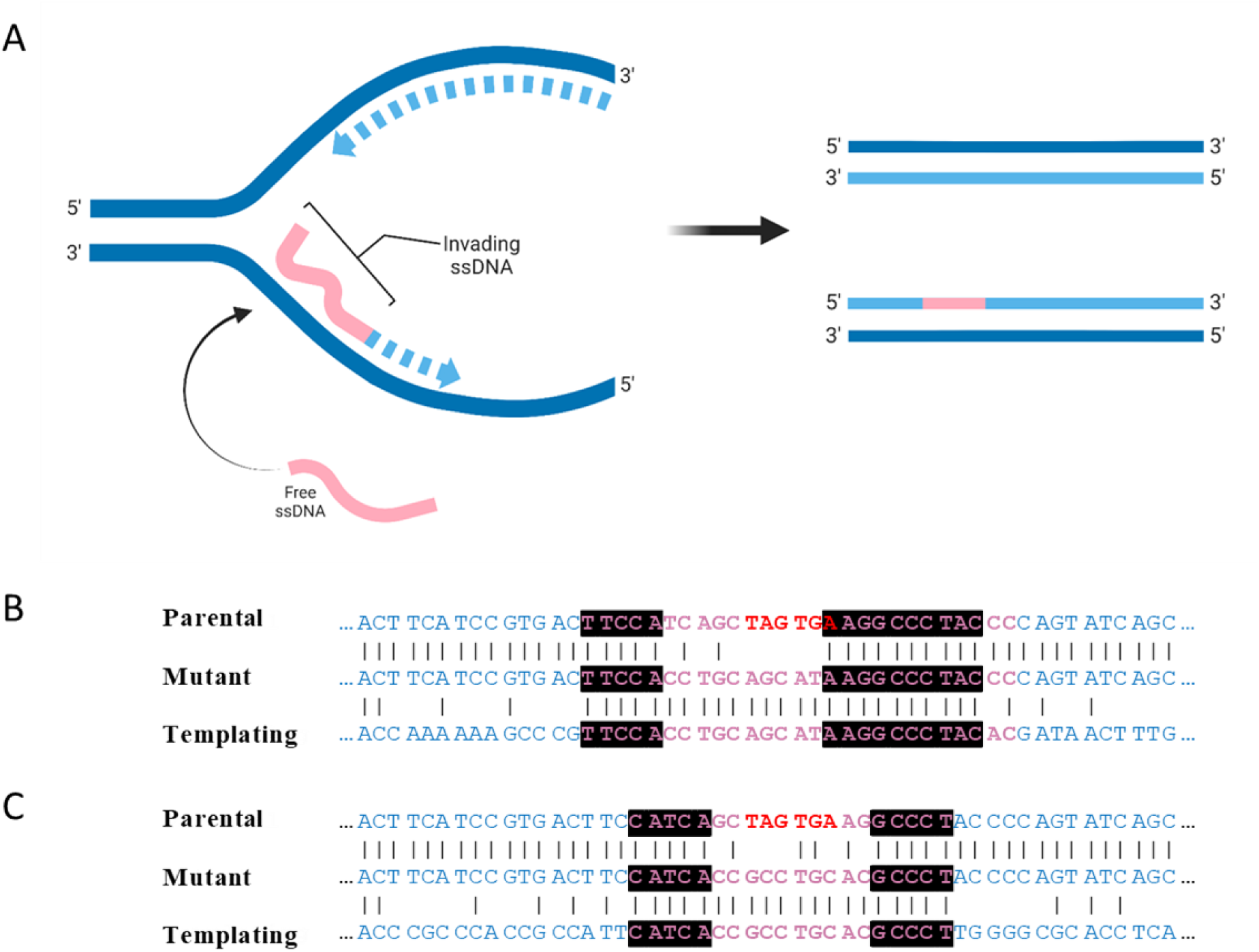
SPDIR mutation mechanism. A: a cytoplasmic DNA single-strand (pink) can anneal with a transiently single-stranded section of the chromosome, e.g. with the discontinuously replicated strand at a replication fork (left). The annealed strand is then extended in the course of DNA replication (light blue dashed arrows) and finally gets integrated into the nascent daughter strand (right). After an additional round of replication, the mutation is fixed. B, C: Examples of experimentally found SPDIR mutations, shown as triple DNA alignments of the parental sequence (top), the resulting mutant sequence (center) and the sequence of the templating DNA (bottom). The microhomologies (determined by ΔG^0^_min_ calculations) are depicted in pink, except for the parental stop codons (red). The two illegitimate crossover joints are highlighted in black. B: recurrent mutation A26 caused with templating DNA from the *A. baylyi* chromosome; C: novel mutation MK45 caused with DNA originating from plasmid pQLICE. Both mutations result in His^+^ reversion through replacement of the two stop codons by the templating DNA. All SPDIR mutations found in this study are reported in Supplemental file S1. Panel A was created using Biorender.com.

SPDIR mutations have been found to be very rare (between 10^-12^ and 10^-13^ per locus and cell in the gamma-proteobacterial model organism *Acinetobacter baylyi*; Liljegren et al., 2024). The SPDIR mutation mechanism is probably widespread and potentially universal, and SPDIR mutations have been identified retrospectively *in silico* in the Gram-positive *Streptococcus pneumoniae* and in the human genome as well (Harms et al., 2016). However, due to the very low frequency and to challenges in detecting microindel mutations (both experimental and bioinformatic), reports are lacking. Remarkably, SPDIR frequencies can increase by orders of magnitude under genotoxic stress, in the course of natural transformation (Harms et al., 2016), or through disruption of specific genome maintenance functions (Harms et al., 2016; Liljegren et al., 2024).

The detection and quantification of SPDIR mutations is not trivial. In *A. baylyi*, our lab mainly exploits a modified prototrophy marker gene (*hisC* encoding histidinol phosphate aminotransferase, essential for histidine biosynthesis) to experimentally enrich for naturally occurring SPDIR mutants. In the *hisC*::’ND5i’ allele, the *hisC* open reading frame is interrupted by a 228-bp insert containing two consecutive stop codons that prevent its expression (Overballe-Petersen et al., 2013; Harms et al., 2016; Fig. 2). Cells carrying *hisC*::’ND5i’ grow normally in rich medium (containing histidine) but cannot form colonies on minimal medium unless supplemented with histidine. Only cells that have the two stop codons removed in frame, or have acquired a new start codon in frame downstream of the stop codons, become His^+^, and subsequent DNA sequencing of the mutant *hisC* allows the classification of the mutation (typically, *in frame* deletions from three to 195 bp, and occasionally SPDIR mutations; Fig. 1B, C).

**Fig. 2.**
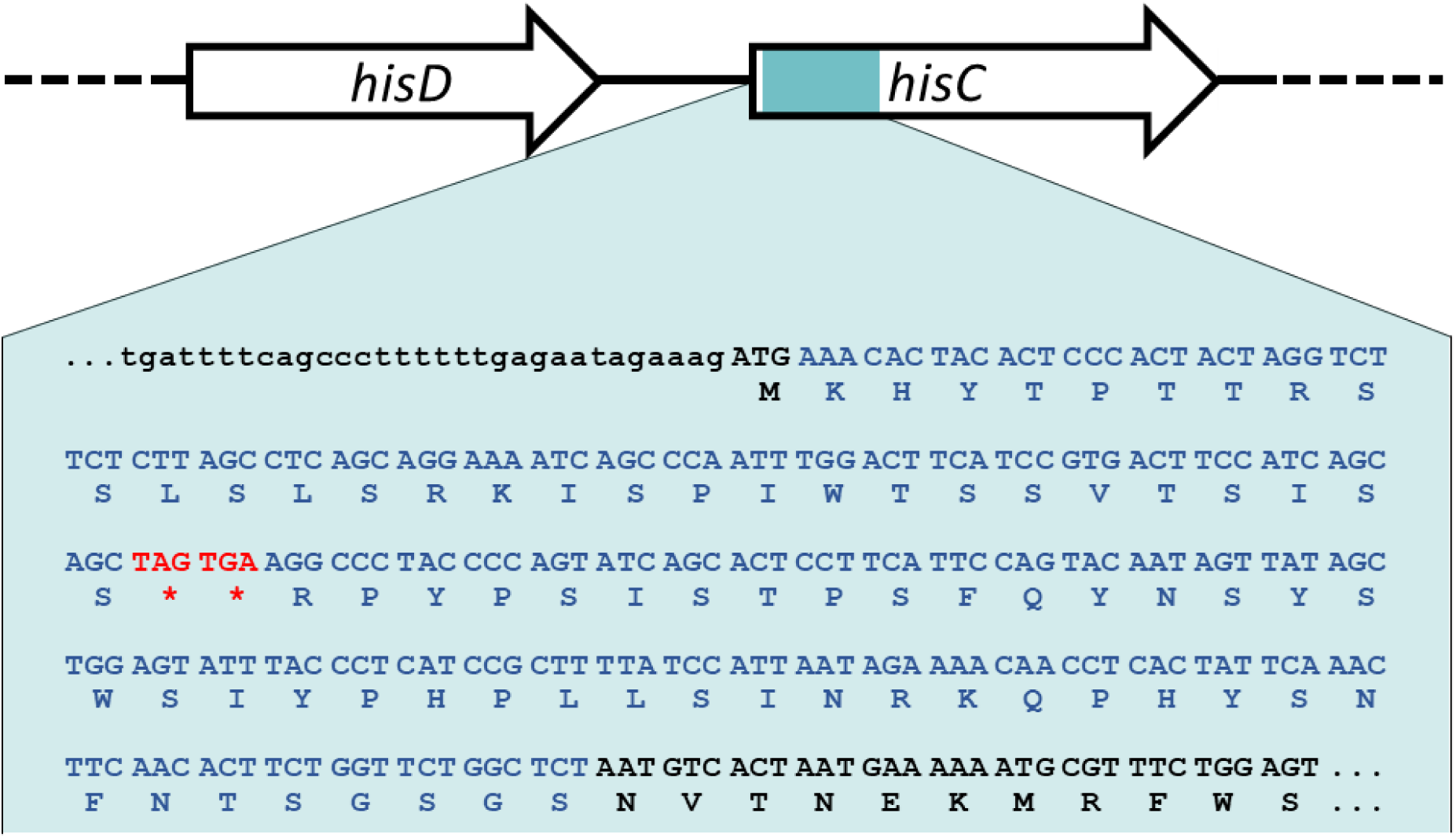
Genomic detail of the *A. baylyi hisC*::’ND5i’ detection allele for SPDIR mutations (Overballe-Petersen et al., 2013; Harms et al., 2016). The ‘ND5i’ insert is shown in blue with DNA sequence and codon details; the stop codons are indicated in red. Lowercase: non-coding DNA.

In our recent study, we investigated the roles of ssDNA-binding proteins (DprA and RecA) on SPDIR mutation frequencies in *A. baylyi* knock-out mutants (Liljegren et al., 2024), and one of our (unreported) experimental approaches was the investigation of overexpression of ssDNA-binding functions in *A. baylyi* using a broad host range overexpression plasmid vector. Unexpectedly, the carriage of the vector interfered with our experimental setup and boosted the generation of SPDIR mutations in *A. baylyi*. Here we investigated the impact of carriage of broad host-range plasmids on SPDIR mutations. Our results demonstrate that specific plasmids can be mutagenic through direct or indirect interactions with the host genome.

## Materials and Methods

### Bacterial strains and plasmids

The strains used in this study were derived from *A. baylyi* ADP1 and were published previously. Strain AL4 [ADP1 *hisC*::’ND5i’ *rpoB1 alkM*::(*nptII’ tg4*); Harms et al., 2016] was used as wildtype, and strain KOM218 (AL4 Δ*recJ* Δ*exoX*; Overballe-Petersen et al., 2013) was employed for quantification of SPDIR events above the limit of detection.

The plasmids used in this paper are listed in Table 1. In addition, overexpression vectors derived from pQLICE (Harms et al., 2007; Fig. 5A) were constructed as follows: the *dprA* (ACIAD0209), *recA* (ACIAD1385) and *ssb* (ACIAD3449) genes of *A. baylyi* ADP1 (GenBank NC_005966) were amplified with Phusion high-fidelity DNA polymerase using the primers dprA-M2-fw/dprA-M2-rv, recA-f4/recA-r4 and ssb-f/ssb-r, respectively (Table 2), and inserted into the *SmaI* site of pQLICE, resulting in the plasmids pQLICE-dprA, pQLICE-recA, and pQLICE-ssb, respectively. Cloning steps were performed using *Escherichia coli* DH5⍺ (Hanahan, 1983). The plasmid constructions were verified by PCR and by restriction analysis.

**Table 1:** List of plasmids.

| Plasmid | Antibiotic selection [mg/L] | Incompatibility group | Replication mechanism | Size [kbp] | Accession number |
| --- | --- | --- | --- | --- | --- |
| pQLICE | streptomycin 40 | IncQ | strand displacement (Sakai & Komano, 1996)) | 10.546 | EF189157 |
| pBBR1MCS-3 | tetracycline 10 | pBBR | theta? (Antoine & Locht, 1992) | 5.228 | XXU25059 |
| pRK415 | tetracycline 10 | IncP-1α | theta (Johnson, 2018) | 10.690 | EF437940 |
| R16a | kanamycin 25 | IncA/C<sub>2</sub> | theta (Johnson & Lang, 2012; Johnson, 2018) | 173.094 | KX156773 |
| pK71-77-1-NDM | ampicillin 100 | IncA/C<sub>2</sub> | theta (Johnson & Lang, 2012; Johnson, 2018) | 145.272 | CP040884 |
| RN3 | tetracycline 10 | IncN | theta (Johnson, 2018) | 54.205 | FR850039 |
| R388 | trimethoprim 250 | IncW | theta (Johnson, 2018) | 33.913 | NC_028464 |

**Table 2:**
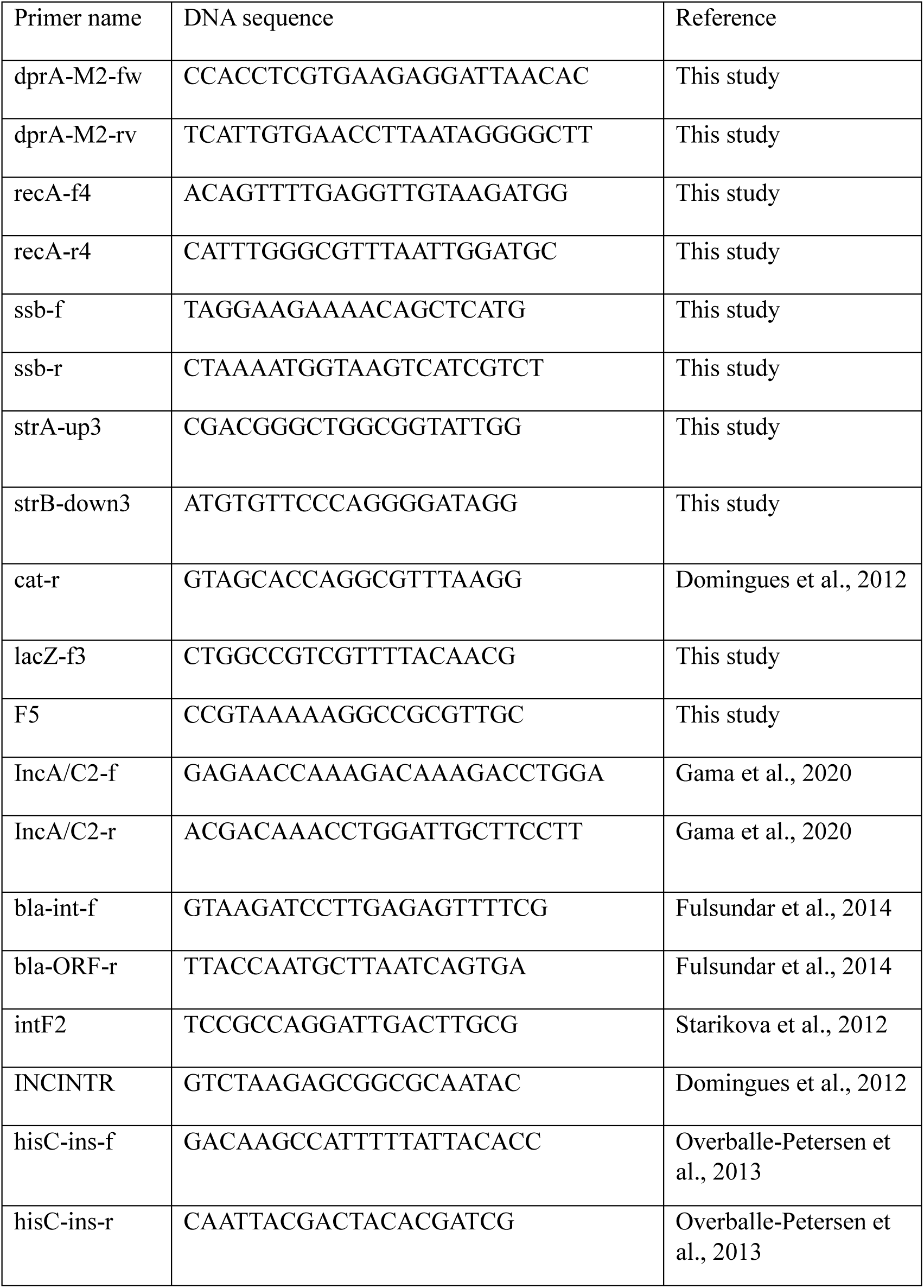
List of primers.

The pQLICE plasmid and its derivatives, as well as pRK415 and pBBR1MCS-3, were introduced into the *A. baylyi* strains by natural transformation as published (Harms et al., 2007) and confirmed by PCR (primers: strA-up3/strB-down3 for pQLICE; cat-r/lacZ-f3 for pBBR1MCS-3; F5/lacZ-f3 for pRK415; Table 2). The remaining plasmids were transferred from *E. coli* MG1655 derivatives (Δ*ara* or Δ*malF*) to *A. baylyi* by conjugation (Fig. 3A). Briefly, the donor strains were grown individually in shaking culture tubes [1 mL Lysogeny Broth (LB) broth amended with an appropriate antibiotic; Table 1] at 37°C to stationary phase. The recipient strains were grown accordingly with 50 mg/L rifampicin at 30°C. The strains were harvested, washed and resuspended in fresh LB. 5 µL of donor and 5 µL of recipient were added to 490 µL of fresh LB, and the mating cultures were incubated at room temperature overnight without shaking. 200 µL of the mating cultures were plated on double-selective LB media (50 mg/L rifampicin plus plasmid-specific selective antibiotics; Table 1). Colonies were restreaked on appropriate medium and verified by host-specific (recA-f4/recA-r4 targeting the *A. baylyi recA* gene) and by plasmid-specific PCR (IncA/C2-f and IncA/C2-r for pK71-77-1-NDM and R16a; bla-int-f/bla-ORF-r for R16a; and intF2/INCINTR targeting the different integrons of pK71-77-1-NDM, RN3 and R388, respectively). All primers are listed in Table 2.

**Fig. 3.**
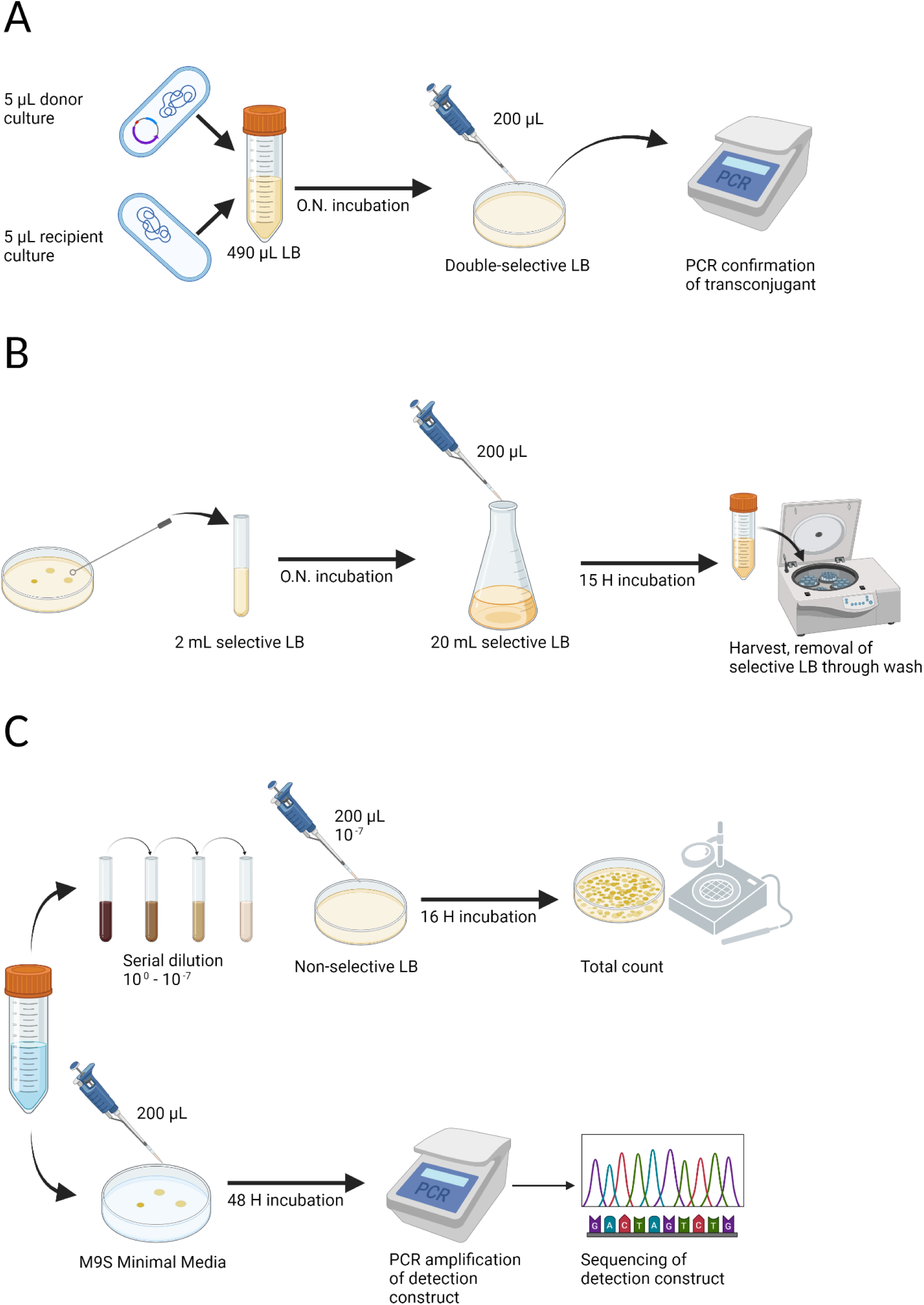
Overview of experimental methodology. **A:** Conjugation assay for strain constructions. Donor strains (plasmid-carrying *E. coli*) were mated with recipient *A. baylyi* strains overnight and then plated on double-selective media. Isolates were purified and confirmed by PCR for plasmid DNA and for *A. baylyi* genomic DNA. **B:** Mutation assay. *A. baylyi* strains with or without plasmids were pre-grown for 16-18 hours, diluted 1:200, and grown in 20 ml cultures under aeration for 15 hours. Plasmid-carrying strains were grown under appropriate antibiotic selection. The cells were then washed twice in PBS and resuspended in 2 mL PBS. **C:** His^+^ frequency determination: 20 µL of the washed cells of step B were diluted 10^-7^, plated on LB and incubated for 15 hours (total cfu determination). The rest of the cells (∼2 ml) were evenly distributed on ten M9S minimal medium plates (selective for His^+^ mutants) and incubated 48 hours. Colonies were restreaked on M9S, and the *hisC* allele of each isolate was amplified and Sanger-sequenced. Mutations were called using pairwise BLAST. Created with Biorender.com.

### Mutation assays

The mutation assays were conducted as reported previously (Liljegren et al., 2024) and illustrated in Fig. 3B. In brief, *A. baylyi* starter cultures were grown in LB overnight in a shaker (all incubation steps were conducted at 30°C). From each starter culture, a single 20 mL-culture was inoculated in LB 1:100 and incubated with aeration for 15 hours. The cells were harvested, washed twice with ice-cold phosphate-buffered saline (PBS) and resuspended in 2 ml PBS. The cells were then distributed on M9 minimal medium supplemented with 10 mM succinate (M9S; 200 µl cells per plate; for His^+^ mutant titer determination) and in appropriate dilution on LB (cfu titer determination) (Fig. 3C). The plates were incubated for 16 (LB) or 48 hours (M9S). Next, the total cfu and His^+^ titers were determined and the His^+^ frequencies were calculated as mutant titer per cfu titer, and for each group of parallel experiments, we determined the median His^+^ frequency. When strains carried plasmids, both cultures (starter culture and 20 mL 15-hour growth cultures) were amended with selective antibiotics at concentrations given in Table 1 unless indicated otherwise. For some control experiments, 1 mg/L streptomycin (below minimal inhibitory concentration) was added to plasmid-free cultures.

### SPDIR frequency determination

The His^+^ colonies were restreaked on M9S medium and grown for 48 hours at 30°C. From each isolate, the recombinant section of the *hisC*::’ND5i’ allele was amplified by PCR using DreamTaq (Thermo Scientific) and primers hisC-ins-f/hisC-ins-r (Table 2), purified according to the Exo-SAP procedure (NEB), and Sanger-sequenced by Azenta Life Services GenWiz (sequencing primer: hisC-ins-f). SPDIR mutations were clearly identified and separated from other His^+^ mutations (typically, small deletions ranging from 3 to 195 bp in size) with pairwise BLAST against the *A. baylyi* genome (GenBank: NC_005966) and, when appropriate, the plasmid genomes (GenBank column in Table 1) as subject. When identical mutations were identified from the same experiment, the mutants were conservatively regarded as siblings and were scored as a single mutation event. For SPDIR mutations, we identified the templating DNA sections (from the *A. baylyi* chromosome or from the plasmids) and confirmed the SPDIR mechanism by calculating the minimal Free Energy of Hybridization (using the nearest neighbour and mismatch/gap penalty parameters from Wetmur, 2006) at each illegitimate crossover joint. All SPDIR mutations identified in this study are listed in Supplemental file 1. SPDIR frequencies were determined as number of SPDIR events divided by the number of all His^+^ events, multiplied with the overall median His^+^ frequency (“calculated SPDIR frequencies”). For statistical evaluations of experiments with the *A. baylyi* Δ*recJ* Δ*exoX* strain, we additionally calculated the “median SPDIR frequencies” (regardless of His^+^ events).

### Plasmid stability determination

Each *A. baylyi* KOM218 strain containing plasmids was inoculated from a single culture in 2 mL LB and grown for 15 hours at 30°C without antibiotics. The cells were harvested, diluted 10^-7^ in PBS, and 200 µL were plated out on LB with and without selective antibiotic (Supplemental Table S1). Colonies were counted after incubating the plates for 16 hours at 30°C. The plasmid stabilities were determined as colony counts on selective per non-selective medium (Supplemental Table S1).

### Statistical Analysis

Statistical analyses were performed in R (R Core Team. 2018) version 4.3.2. Package PMCMRplus (Pohlert. 2023) was required for the Dunn many-to-one post hoc test (kwManyOneDunnTest). Graphs were produced with packages ggplot2 (Wickham, 2016), patchwork (Pedersen, 2020), ggExtra (Attali & Baker, 2023), and rcartocolor (Nowosad, 2023).

## Results

### The overexpression vector pQLICE stimulates SPDIR

We cloned each of the *dprA, recA* and *ssb* genes of *A. baylyi* separately into the IncQ plasmid broad host range expression vector pQLICE and inserted the resulting plasmids individually into an *A. baylyi* Δ*recJ* Δ*exoX hisC*::’ND5i’ strain to investigate how overexpression of each of these genome maintenance functions affected suppression of microindel mutations caused by SPDIR. Unexpectedly, SPDIR frequencies were increased up to tenfold compared with the plasmid-free *A. baylyi* counterpart (Table 3). We repeated the experiments with the empty pQLICE vector and found that its presence increased the calculated SPDIR frequencies 3.5-fold (Table 3). Comparing the median SPDIR frequencies (Table 3) revealed that the increase was significant (Wilcoxon rank sum test, p = 0.004). Moreover, with pQLICE, we found that more than half of the individual His^+^ mutations were caused by SPDIR. In contrast, in absence of the plasmid, the fraction of SPDIR mutations was less than one third (Table 3).

**Table 3.** SPDIR frequencies of *A. baylyi* Δ*recJ* Δ*exoX hisC*::’ND5i’ with and without pQLICE plasmid derivatives.

| Plasmid vector | Median His<sup>+</sup> frequency | SPDIR event per His<sup>+</sup> event | Calculated SPDIF frequency | Fold change | Median SPDIF frequency (± interquartile range)* | n |
| --- | --- | --- | --- | --- | --- | --- |
| - | 2.5×10<sup>-10</sup> | 32% (43/135) | 8.0×10<sup>-11</sup> | =1 | 7.6×10<sup>-11</sup> ± 1.5×10<sup>-10</sup> | 39 |
| pQLICE | 5.4×10<sup>-10</sup> | 52% (90/172) | 2.8×10<sup>-10</sup> | 3.5× | 2.5×10<sup>-10</sup> ± 6.3×10<sup>-10</sup> | 26 |
| pQLICE-dprA | 1.5×10<sup>-9</sup> | 59% (42/71) | 8.0×10<sup>-10</sup> | 10× | 8.5×10<sup>-10</sup> ± 7.7×10<sup>-10</sup> | 12 |
| pQLICE-recA | 1.1×10<sup>-9</sup> | 47% (40/86) | 5.2×10<sup>-10</sup> | 6.5× | 5.4×10<sup>-10</sup> ± 6.0×10<sup>-10</sup> | 12 |
| pQLICE-ssb | 1.1×10<sup>-9</sup> | 45% (25/56) | 4.8×10<sup>-10</sup> | 6.0× | 4.0×10<sup>-10</sup> ± 8.7×10<sup>-10</sup> | 13 |

\* values estimates from N experiments as indicated in Suppl. Fig. S2A.

We repeated the experiments in the *recJ*^+^ *exoX*^+^ (“wildtype”) *A. baylyi hisC*::’ND5i’ strain. In this strain, SPDIR frequencies are generally about 30-fold lower than in a Δ*recJ* Δ*exoX* background (Harms et al., 2016; Liljegren et al., 2024), precluding significance statements due to zero inflation, but the tendencies were very similar to the results observed in the Δ*recJ* Δ*exoX* strain. Without plasmid, the median His^+^ frequency was the lowest, and no SPDIR mutant was recovered. With pQLICE or its derivatives, the His^+^ frequencies were at least 2-fold higher, and altogether three SPDIR events were observed (Table 4).

**Table 4.** SPDIR frequencies of *A. baylyi hisC*::’ND5i’ with and without pQLICE plasmid derivatives.

| Plasmid vector | Median His<sup>+</sup> frequency | SPDIF event per His<sup>+</sup> event | Calculated SPDIF frequency | n |
| --- | --- | --- | --- | --- |
| - | 5.7×10<sup>-11</sup> | <2.5% (0/39) | <1.8×10<sup>-12</sup> | 34 |
| pQLICE | 2.2×10<sup>-10</sup> | 3.6% (2/56) | 7.7×10<sup>-12</sup> | 26 |
| pQLICE-dprA | 2.7×10<sup>-10</sup> | <4.0% (0/25) | <1.1×10<sup>-11</sup> | 12 |
| pQLICE-recA | 2.0×10<sup>-10</sup> | <4.2% (0/24) | <8.1×10<sup>-11</sup> | 13 |
| pQLICE-ssb | 1.2×10<sup>-10</sup> | 3.4% (1/29) | 4.2×10<sup>-12</sup> | 13 |

pQLICE-carrying strains were grown with 40 mg/L streptomycin to prevent plasmid loss, while plasmid-free strains were not exposed to antibiotics. We performed control experiments to examine whether the increased SPDIR frequencies were due to plasmid carriage or to antibiotic treatment. We cultured the *A. baylyi* Δ*recJ* Δ*exoX* strain at subinhibitory concentrations of streptomycin (1 mg/L) and obtained a median SPDIR frequency of 6.1×10^-11^ (n = 7) that was not significantly different from the experiment without antibiotic (Wilcoxon rank sum test, p = 0.75). Furthermore, experiments with pQLICE-carrying *A. baylyi* Δ*recJ* Δ*exoX* performed in the absence of selection for the plasmid revealed a median SPDIR frequency of 1.4×10^-10^ (n = 9). Despite high plasmid loss (approximately 40%, Supplemental Table S1) the absence of selection did not significantly affect the median SPDIR frequency compared to streptomycin selection (Wilcoxon rank sum test, p = 0.30). In addition, we recovered one SPDIR event in eight experiments with wildtype *A. baylyi hisC*::’ND5i’ carrying pQLICE without selection. In contrast, in seven experiments no SPDIR events were detected with plasmid-free *A. baylyi* wildtype when grown with subinhibitory concentrations of streptomycin. Altogether, our results show that pQLICE carriage increases SPDIR frequencies in *A. baylyi*, independently of host genotype and antibiotic treatment.

### Distinct plasmids modulate SPDIR differently

The results with pQLICE led us to test six additional broad host-range plasmids (Table 1) to evaluate their effect on SPDIR mutation frequencies, and the results are summarized in Table 5. We observed SPDIR mutants in 15 out of 17 experiments (88%) conducted with *A. baylyi* Δ*recJ* Δ*exoX* strain carrying the IncP-1 RK2-derived cloning vector pRK415, demonstrating a higher ratio compared to its plasmid-free counterpart (23 out of 39; 59%). Concomitantly, carriage of pRK415 resulted in a calculated SPDIR frequency of 1.2×10^-9^, indicating a 17-fold increase relative to the plasmid-free strain. pRK415 also appears to stimulate SPDIR in the wildtype strain because we detected a SPDIR mutant in the presence, but not in the absence of pRK415. Therefore, we conclude that the effect of increasing the frequency of SPDIR mutations is not restricted to pQLICE.

**Table 5.** SPDIR frequencies of *A. baylyi hisC*::’ND5i’ strains carrying plasmids.

| Plasmid | Relevant *A. baylyi* genotype | Median His<sup>+</sup> frequency | SPDIR event per His<sup>+</sup> event | Calculated SPDIF frequency | fold change relative to $\Delta recJ \Delta exoX$ | Median SPDIF frequency (± interquartile range)* | n |
| --- | --- | --- | --- | --- | --- | --- | --- |
| pRK415 | $\Delta recJ \Delta exoX$ | $2.2 \times 10^{-9}$ | 53% (49/93) | $1.2 \times 10^{-9}$ | 17× | $1.3 \times 10^{-9} \pm 2.8 \times 10^{-9}$ | 17 |
| pRK415 | wildtype | $1.3 \times 10^{-10}$ | 4% (1/25) | $5.0 \times 10^{-12}$ | | | 11 |
| pBBR1MCS-3 | $\Delta recJ \Delta exoX$ | $6.7 \times 10^{-10}$ | 30% (26/87) | $2.0 \times 10^{-10}$ | 2.5× | $1.4 \times 10^{-10} \pm 3.1 \times 10^{-10}$ | 22 |
| pBBR1MCS-3 | wildtype | $3.0 \times 10^{-10}$ | <1% (0/100) | $<3.0 \times 10^{-12}$ | | | 21 |
| R16a | $\Delta recJ \Delta exoX$ | $4.3 \times 10^{-10}$ | 30% (6/20) | $1.3 \times 10^{-10}$ | 1.6× | $2.4 \times 10^{-11} \pm 1.5 \times 10^{-10}$ | 10 |
| R16a | wildtype | $2.3 \times 10^{-10}$ | <7% (0/14) | $<1.7 \times 10^{-11}$ | | | 10 |
| pK71-77-1-NDM | $\Delta recJ \Delta exoX$ | $6.4 \times 10^{-10}$ | 27% (8/30) | $1.7 \times 10^{-10}$ | 2.1× | $1.7 \times 10^{-10} \pm 3.3 \times 10^{-10}$ | 10 |
| RN3 | $\Delta recJ \Delta exoX$ | $1.4 \times 10^{-11}$ | 50% (1/2) | $7.0 \times 10^{-12}$ | 0.09× | $1.4 \times 10^{-11} \pm 2.8 \times 10^{-11}$ | 6 |
| RN3 | wildtype | $<4.0 \times 10^{-12}$ | n.a. | n.a. | | | 6 |
| R388 | $\Delta recJ \Delta exoX$ | $1.0 \times 10^{-10}$ | 19% (3/16) | $1.9 \times 10^{-11}$ | 0.24× | $2.5 \times 10^{-11} \pm 7.5 \times 10^{-11}$ | 8 |
| R388 | wildtype | $4.2 \times 10^{-11}$ | 20% (1/5) | $8.4 \times 10^{-12}$ | | | 6 |

n.a., not applicable
\* values estimates from n experiments as indicated in Fig. 4.

Carriage of plasmids pBBR1MCS-3, R16a and pK71-77-1-NDM resulted in small (approximately two-fold) changes in the calculated SPDIR frequency of the *A. baylyi* Δ*recJ* Δ*exoX* strain, and no SPDIR mutants were observed when the wildtype strain carried these plasmids (Table 5). Thus, these three plasmids likely have no quantifiable effects on SPDIR.

In contrast, carriage of the IncN plasmid RN3 decreased the calculated SPIDR frequency of *A. baylyi* Δ*recJ* Δ*exoX* more than 10-fold, and in the wildtype no His^+^ mutants were encountered in six experiments. The IncW plasmid R388 exhibited an ambiguous behaviour as SPDIR mutants were detected in experiments with the wildtype strain but the calculated SPDIR frequency of *A. baylyi* Δ*recJ* Δ*exoX* decreased roughly four-fold (Table 5).

We further calculated the median SPDIR frequencies for the plasmid-carrying derivatives of *A. baylyi* Δ*recJ* Δ*exoX*. A comparison against the plasmid-free strain revealed significant differences (Kruskal-Wallis rank sum test, d.f. = 7, p = 4.61×10^-6^, followed by Dunn many-to-one post hoc test) for plasmids pQLICE and pRK415 (p < 0.02, Fig. 4). This result strongly suggests that the effect on SPDIR is plasmid-specific, and that both pQLICE and pRK415 increase SPDIR in *A. baylyi* Δ*recJ* Δ*exoX*.

**Fig. 4.**
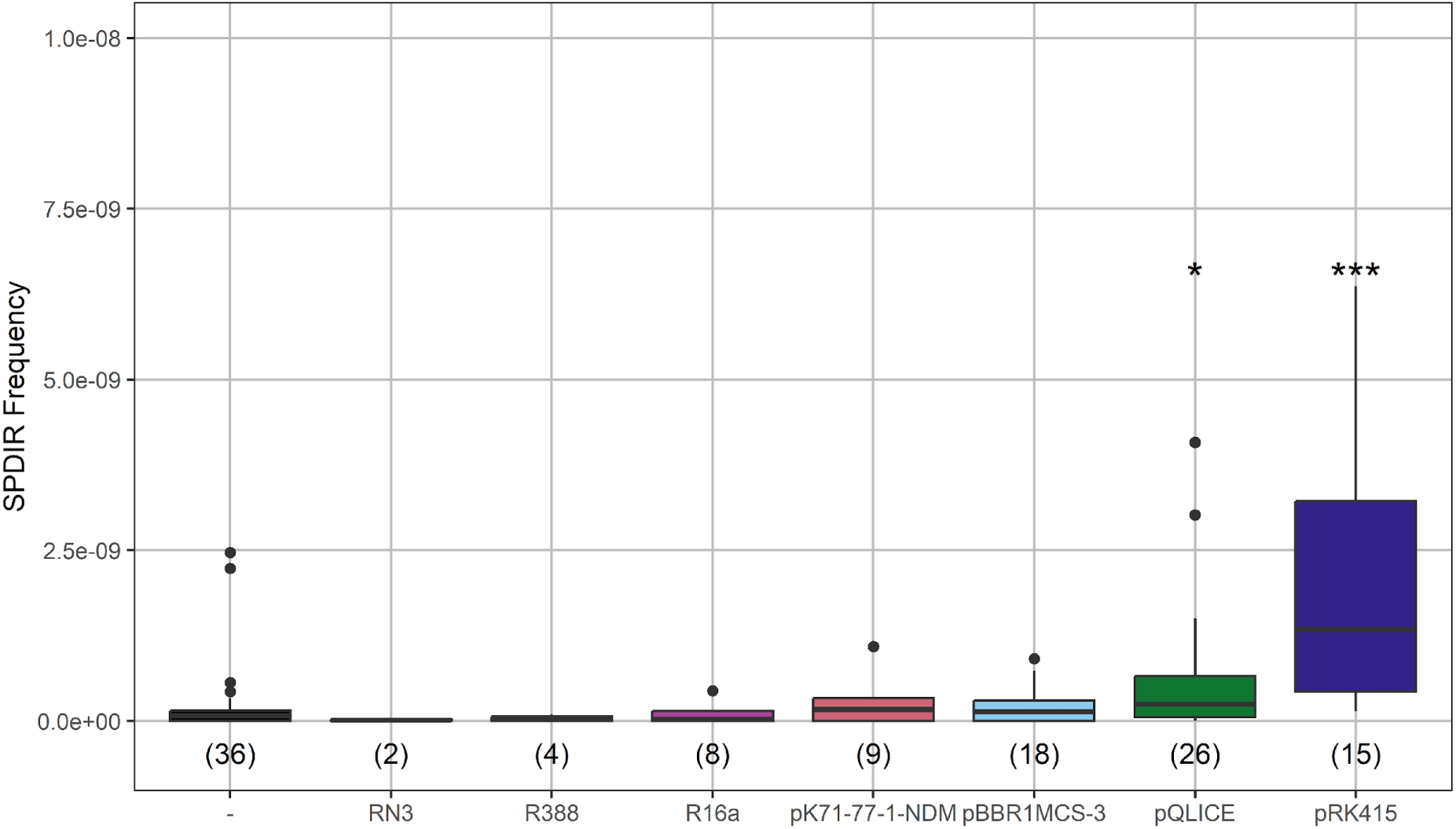
Effect of plasmid carriage on SPDIR in *A. baylyi ΔrecJ* Δ*exoX*. SPDIR frequencies were determined only for experiments where His^+^ mutants were detected. The number of experiments is indicated in parenthesis for each plasmid; - indicates the plasmid-free strain. Significant results of a Kruskal-Wallis rank sum test followed by Dunn many-to-one post hoc test (with the plasmid-free strain as reference group) are indicated as * and *** for p = 0.019 and p = 1.89×10^-6^.

### pQLICE but not pRK415 adds to the pool of templating DNA for SPDIR mutations

In this study, we recovered 114 different SPDIR mutations, unique or recurrent (Supplemental files S1 and S2). Of these mutations, 101 were produced with chromosomal DNA and 13 with DNA originating from plasmids. Among the plasmids that we found to significantly stimulate SPDIR mutations, we recovered 12 SPDIR mutations formed with ectopic DNA from pQLICE but none from pRK415. In addition, one SPDIR was formed with DNA from pK71-77-1-NDM.

The chromosome of *A. baylyi* ADP1 (3.6 Mbp; Barbe et al., 2004) is a much larger resource for templating DNA than the extrachromosomal plasmids (pQLICE: 10.5 kbp; pRK415: 10.7 kbp; pK71-77-1-NDM: 145 kbp; Table 6). We calculated the molecular frequency (density) of SPDIR events formed with the different genomic molecules in relevant *A. baylyi* Δ*recJ* Δ*exoX* strains. SPDIR with templating pQLICE DNA resulted in a density of 6.6×10^-2^ SPDIR mutations per experiment per kbp of pQLICE DNA. This value exceeds the density of SPDIR events formed with templating chromosomal DNA in the same experiments (7.7×10^-4^) by 85-fold, and in the plasmid-free *A. baylyi* control experiments by 214-fold (Table 6). For SPDIR mutations templating with pK71-77-1-NDM DNA (altogether one SPDIR recovered), the density is somewhat higher than the density with chromosomal DNA (Table 6). With pRK415 DNA, the SPDIR-causing density was below detection limit (Table 6). Taken together, carriage of both pQLICE or pRK415 increase the SPDIR frequency. However only with pQLICE, the plasmid DNA is frequently used as templating source for SPDIR mutations, exceeding the relative usage of chromosomal templating DNA. Even when accounting for an elevated plasmid copy number (10 to 12 for IncQ plasmid RSF1010; Meyer 2009), pQLICE DNA produces at least seven times more SPDIR mutations than the chromosome. Moreover, the differences between pQLICE and pRK415 suggest different mechanisms for causing SPDIR mutations.

**Table 6.** Templating density of different molecules in *A. baylyi* Δ*recJ* Δ*exoX hisC*::’ND5i’ with and without plasmids.

| Plasmid carriage | templating DNA [bp] | templating density [events per kbp per experiment] | SPDIR events | n |
| --- | --- | --- | --- | --- |
| none | chromosome<br>(3,598,621) | $3.1 \times 10^{-4}$ | 43 | 39 |
| pQLICE | pQLICE (10,546) | $6.6 \times 10^{-2}$ | 18 | 26 |
| pQLICE | chromosome<br>(3,598,621) | $7.7 \times 10^{-4}$ | 72 | 26 |
| pK71-77-1-NDM | pK71-77-1-NDM<br>(145,272) | $6.9 \times 10^{-4}$ | 1 | 10 |
| pK71-77-1-NDM | chromosome<br>(3,598,621) | $1.9 \times 10^{-4}$ | 7 | 10 |
| pRK415 | pRK415 (10,690) | $<5.5 \times 10^{-3}$ | 0 | 17 |
| pRK415 | chromosome<br>(3,598,621) | $7.0 \times 10^{-4}$ | 43 | 17 |

### The replication mechanism of pQLICE can explain the contribution to SPDIR mutations

pQLICE (Fig. 5A) belongs to the IncQ family of plasmids (Meyer. 2009) which replicate via the characteristic strand displacement mechanism: both DNA strands (plus and minus) are replicated as leading strands starting from the *oriV* (Fig. 5B). As a result, the displaced strands spend part of their life cycle as single-strand until they are replicated as final stretches of the leading strands, and the single-stranded intermediates are thought to be the substrates for SPDIR mutations (Fig. 5C).

**Fig. 5.**
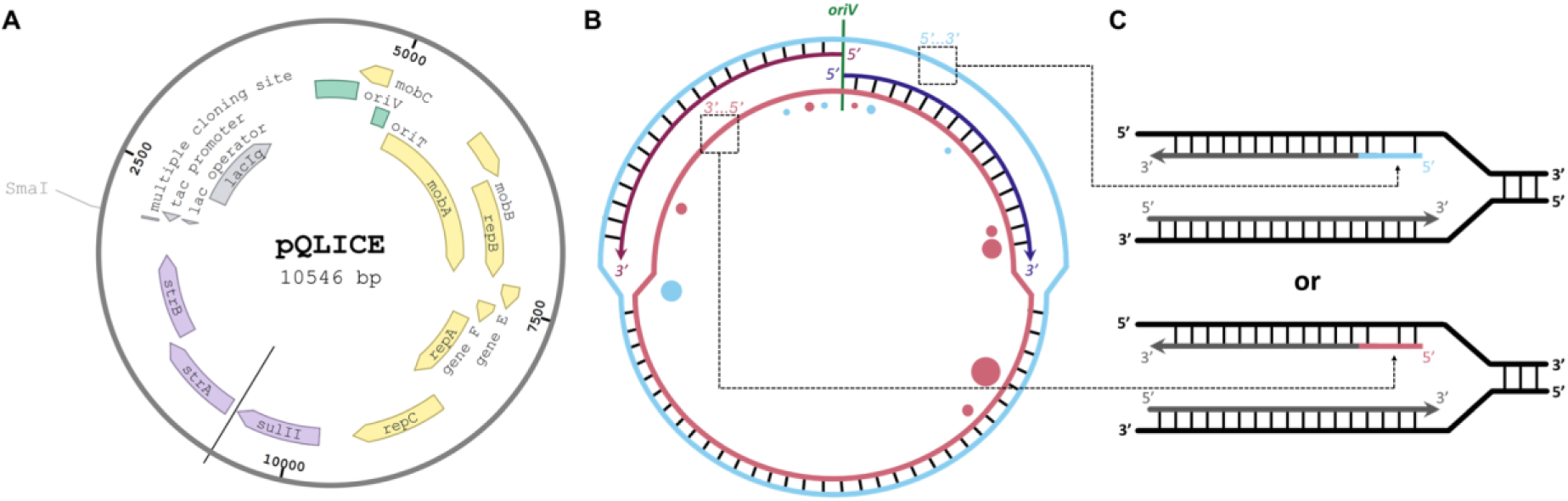
Illustration of pQLICE strand-displacement replication and mechanistic implication for SPDIR formation. A. Genetic map of pQLICE. Genes/loci are color coded as: green, origins (*oriV*, replication at the top of the map; *oriT*, transfer); yellow, replication and mobilization; purple, antibiotic resistance; gray, cloning/expression. Black line indicates the start of the plasmid annotation (GenBank: EF189157). The map was drawn with Benchling [Biology Software] (2024), retrieved from https://benchling.com. B. Model of the strand-displacement replication mechanism of pQLICE. Circles indicate the positions of the experimentally found templating ssDNA segments; the colors represent the respective DNA strands templating the SPDIR mutation (light blue: plus strand; light pink: minus strand) and the size is proportional to the count of independent events. C. Chromosomal replication fork; the dashed arrows from B to C indicate the ssDNA from pQLICE (light blue: plus strand; light pink: minus strand) annealing with the discontinuously replicated strand. The annealed strand is then integrated in the nascent discontinuously replicated strand, resulting in a chromosomal SPDIR mutation.

We investigated whether we could explain the observed pattern of templating DNA from pQLICE (filled circles in Fig. 5B). In wildtype *A. baylyi* grown under benign conditions, templating DNA for SPDIR mutations preferentially stemmed from a chromosomal section around the terminus of replication (Harms et al., 2016), and we investigated whether the templating segments of pQLICE-derived plasmids displayed a comparable bias. We found that half of the unique SPDIR templates originated instead from a region surrounding the plasmid origin of replication (*oriV*) with apparent diminishing occurrence as the distance to *oriV* increased (Fig. 5B, Suppl. Fig. S1). This result alone, however, cannot explain the full mutation pattern observed.

We hypothesized that the time each molecule remained single-stranded contributed to the SPDIR events. Each leading strand is replicated 5’ to 3’, and therefore the section 3’-oriented of *oriV* is exposed as ssDNA longest. However, the templating ssDNA sources are distributed equally over the 3’- and 5’- segments of *oriV*, suggesting no preference for the time a DNA was rendered as single-strand.

The formation of SPDIR mutations depends on the stability of hybridization of complementary DNA single-strands at one or more (extended) microhomologies (Harms et al., 2016). As a proxy for the stability of annealed single-strands, we calculated the minimal Free Energies of Hybridization (ΔG^0^_min_) for each SPDIR event (Supplemental File S1). We found three hotspots for SPDIR events (identical mutations found more than five times in 72 independent experiments), and these were among the group with the lowest ΔG^0^_min_ values reported here (indicating highest hybridization affinity; Suppl. Fig. S1). While the ΔG^0^_min_ values alone cannot explain the frequency of the SPDIR events generated with DNA from pQLICE-derived vectors, there is a significant correlation between the Free Energies of Hybridization and the occurrence of SPDIR events (Spearman’s rank correlation, p = 0.02, rho = -0.65), suggesting ΔG^0^_min_ likely as a contributing factor.

In summary, this result confirms our previous experimental observations that ssDNA can cause SPDIR mutations (Harms et al., 2016; Liljegren et al., 2024). While single-stranded DNA molecules are normally rare in healthy cells, they are frequently generated as part of the replication cycle of IncQ plasmids. This trait can explain the observed mutagenicity of pQLICE for SPDIR mutations.

### Plasmid-carriage does not influence the templating chromosomal DNA pattern for SPDIR

We finally investigated whether plasmids could affect the patterns of SPDIR events generated with chromosomal DNA. When considering all experiments, we observed 101 different events generated with chromosomal DNA (a total of 375 individual SPDIR mutation events), of which 13 were observed both in experiments with and without plasmids. The occurrence of these events follows similar trends among experiments (Fig. 6A). The same trend is observed with pQLICE-derived vectors that contribute to most of the events observed in the presence of plasmids (Fig. 6B), and is independent of the gene cloned (Suppl. Fig. S2B). The 12 unique events generated with chromosomal DNA in the absence of plasmids are very rare (median = 1), as are those generated in the presence of plasmids (median = 1) (Fig. 6C). In conclusion, although plasmids modulate the frequency of SPDIR, the source of templating chromosomal DNA used in recombination remains largely unchanged.

**Fig. 6.**
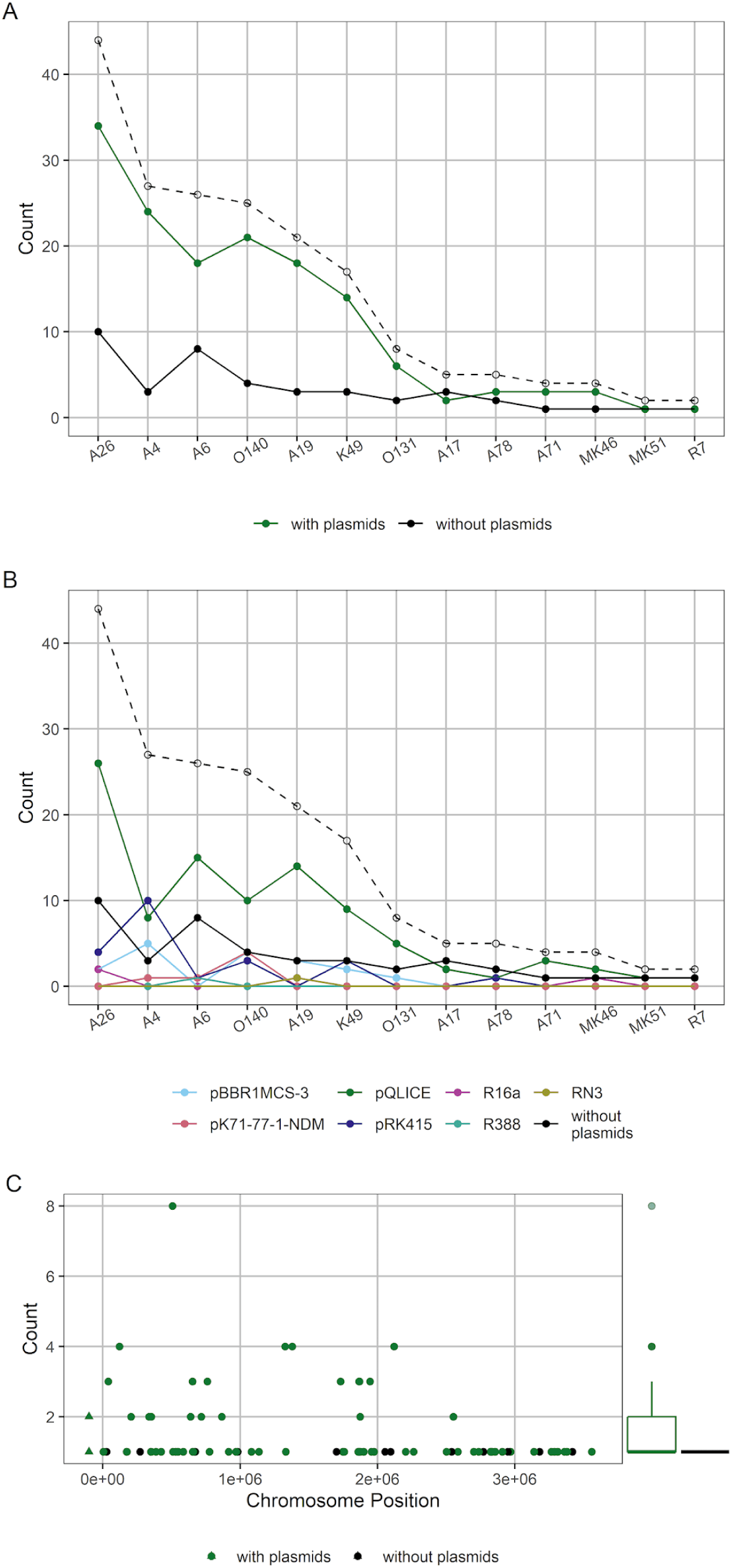
Number of SPDIR events detected in *A. baylyi* independently of plasmid carriage. A: SPDIR events in experiments with vs. without plasmids. B: SPDIR events among experiments with different plasmids. The x-axis in A and B denotes the individual SPDIR mutations found both in strains with and without plasmids. In both A and B the dashed line indicates the total number of events. C: SPDIR events only detected either in experiments with or without plasmids. Triangles at negative chromosome positions indicate rRNA loci (seven different loci in *A. baylyi* that cannot be located precisely).

## Discussion

Plasmids have an important role in bacterial adaptation and the present work expands on the means of how these elements accelerate bacterial evolution. Here, we demonstrate experimentally that carriage of plasmid DNA can be directly mutagenic to the host cell. We observed this finding by investigating the role of plasmids in Short-Patch Double Illegitimate Recombination (SPDIR), exploiting the ability of our model organism *A. baylyi* to revert to a His^+^ phenotype through microindel mutations caused by ssDNA. This mutagenic effect is categorically different from mutagenic traits encoded by plasmids, or from increased mutability occurring during transfer (see introduction).

According to Harms et al. (2016), SPDIR mutations occur by ssDNA molecules in the cytoplasm annealing at (simple or extended) microhomologies with the lagging strand at a replication fork. While experimental evidence for this model exists for DNA taken-up from the environment during natural transformation (confirmed with sequence-tagged heteroduplex DNA; Harms et al., 2016), the sources for SPDIR events formed with intragenomic DNA remained unclear. Our study with pQLICE strongly indicates that ssDNA produced in the course of DNA replication can cause SPDIR mutations directly, similar to taken-up DNA single-strands. The characteristic strand-displacement replication of the IncQ plasmid vector pQLICE generates ssDNA intermediates that can directly interact with the chromosomal DNA through hybdridization at microhomologies. Moreover, most of the SPDIR mutations were found in a strain lacking the 5’-ssDNA exonuclease RecJ and the 3’-ssDNA exonuclease ExoX. These two exonucleases clear the cytoplasm from linear ssDNA remnants (Overballe-Petersen et al., 2013; Harms et al., 2016) but do not affect circular ssDNA. This result suggests that the ssDNA molecules causing SPDIR mutations are susceptible to the ssDNA exonucleases, and we propose a model that the SPDIR-causing pQLICE molecules have been previously damaged by ssDNA breaks. Taken together, it can be concluded that ssDNA molecules causing SPDIR in general are cytoplasmic ssDNA remnants from double-stranded parental molecules.

Besides pQLICE, we determined an increase of SPDIR mutations associated with carriage of pRK415. In contrast to pQLICE, pRK415 DNA did not directly cause SPDIR mutations. However, carriage of either pQLICE or pRK415 increased the frequency of SPDIR mutations originating from chromosomal DNA compared with the isogenic, plasmid-free host (Table 6). These results indicate a different, more indirect mutagenic effect of plasmid carriage on SPDIR, and this effect is not universal, as evident from our experiments with other plasmids. What causes this secondary effect? We speculate that the increase may be a result of interactions between the plasmid and the chromosomal DNA, such as homologous or site-specific recombination attempts, interference of plasmid functions with genomic DNA maintenance, such as error-prone polymerases or attacks by plasmid-encoded functions (see introduction). The expression level of DNA-interacting plasmid genes may also be proportional to the plasmid copy number. While the cloning vectors used here are thought to be low-copy number plasmids, the natural plasmids are monocopy or near-monocopy plasmids. The lower gene dosage for the monocopy plasmids can plausibly explain the lack of increase in SPDIR frequency, however the possibility of them encoding specific (interacting) genes cannot be discarded.

An increase in SPDIR mutation frequency, similar to the effect of pRK415 or pQLICE in this study, has been observed previously with taken-up isogenic DNA during natural transformation (Harms et al., 2016). This observation was explained by interactions of the taken-up DNA with the chromosomal DNA, resulting in abortive recombination, and it is conceivable that the corresponding intermediates or remnants from these recombination attempts are the substrates for causing SPDIR mutations in our experiments.

A single SPDIR mutation was caused by pK71-77-1-NDM DNA in a strain carrying that plasmid. pK71-77-1-NDM is one of the largest plasmids in our study (approximately equivalent to 4% of the *A. baylyi* genome; Table 1), and when calculating the SPDIR density (as SPDIR events caused with this DNA per kbp per experiment), it was somewhat higher than the density of the chromosome. The plasmid (belonging to the IncA/C_2_ group) plausibly replicates by theta replication like the chromosome (Johnson & Lang, 2012), and we hypothesize that the single SPDIR event found is the result of stochastic distribution of SPDIR-causing templating DNA over the entire genome of the plasmid-carrying host. Not all plasmids that modulate the frequency of SPDIR mutations display an incremental effect. Remarkably, carriage of the IncN plasmid RN3 appeared to decrease SPDIR frequency both in the Δ*recJ* Δ*exoX* and wildtype strain of *A. baylyi*. This is the first report of an experimental decrease of SPDIR frequency below wildtype levels and also an interesting example for a plasmid acting as an “anti-mutagen”. Previous experimental setups (using knockout mutants, applying genotoxic stress, adding exogenous DNA for transformation or, in this study, presence of certain plasmids) all resulted in SPDIR increase, and the reason for the decrease associated with RN3 carriage is unclear. We speculate that RN3-encoded functions such as genes for ssDNA-binding proteins that could potentially decrease the free cytoplasmic ssDNA level below wildtype level may be the reason. We reiterate that our initial intention of this study was the investigation of overexpression of genome maintenance functions, including the single-strand-binding protein SSB. While the use of the overexpression vector pQLICE rendered that experimental approach futile, plasmid-free overexpression of *ssb* or of potential gene candidates of RN3 may give further insight into the SPDIR mutation mechanism and its suppression. In conclusion, we show that plasmids can modulate the rate of generation of microindel mutations, which expands their role in promoting bacterial evolution. One of the mechanisms by which plasmids increase SPDIR frequency is through production of ssDNA during replication by strand-displacement. Therefore, future work can delve into how other strategies that produce ssDNA (e.g., rolling-circle replication or conjugative ssDNA-processing) affect SPDIR mutation frequency. The remaining mechanisms, that either increase or decrease SPDIR frequency, speculatively derive from specific gene interactions, which sets up the field for extensive genetic screens. Lastly, like plasmids, other mobile genetic elements may contribute their host cell’s ability to acquire microindel mutations and accelerate bacterial evolution, priming future research avenues.

## Funding

Authors M.M.L. and K.H. were funded by a grant by the Norwegian Research Council (grant number 275672).

## Notes

### Competing Interest Statement

The authors have declared no competing interest.

