## Supplemental file 1 for "Plasmids modulate microindel mutations in *Acinetobacter baylyi* ADP1"

### I. Short-patch double illegitimate recombinant (SPDIR) nucleotide alignments

This file lists the DNA sequences of all individual SPDIR mutations experimentally found in this study with the *hisC*::'ND5i' detection allele (Overballe-Petersen et al., 2013) in *Acinetobacter baylyi* ADP1. The mutations were assigned labels ('MK9' etc.) on their first discovery, and those labels were kept when identical mutations were discovered in subsequent independent experiments. The labeling was continued from Harms et al. (2016) and Liljegren et al. (2024).

Abbreviations:
**A** - ancestral His- DNA sequence (with two consecutive stop codons in **red**).
**R** - recombinant/mutant His+ sequence.
**P** - sequence of templating DNA patch that caused the double illegitimate recombination event (usually ADP1 chromosome); when applicable, **ND** (chromosomally inserted 'ND5i' DNA), **pQ** (pQLICE plasmid DNA) or **pK** (pK71-77-1-NDM plasmid DNA) are used instead.
Vertical bars indicate identical nucleotides. Extended microhomologies are indicated in **blue and bold** typeface. The illegitimate crossovers are highlighted in yellow. When notable, stop and start codons are underlined.
**Pos** - position of the extended microhomology in the respective templating DNA (GenBank numbers: ***A. baylyi***: NC_005966; **ND**: EU153450; **pQ**: EF189157; **pK**: CP040884)
**R translated** or **R tl** - deduced amino acid sequence of the recombinant joint (**lowercase bold and in blue**: changed codons).
$\Delta G^0_{min}$ - minimal Free Energy of Hybridization, calculated for each extended microhomology (Wetmur, 2006).

I. chromosomal templating DNA

**MK9.**

```
                                   |-- net 39 bp loss ---------------------------------------------|
A   ACTACTAGGTCTTCTCTTAGCCTCAGCAGGAAAATCAGCCCAATTTGGACTTCATCCGTGACTTCCATCAGCTAGTGAAGGCCCTACCCCAGTATCAGCACTCCTTCATTCCAGTACAATAGTTATAGCTGGAGTATTTACCCTCATCCGCTTTTATCCATTAATAGAAAACAACCTCAC
    |||||||||||||||||||||||||||||||| ||||||                                       | |||  | || ||||||| |||||||||||||||||||||||||||||||||||||||||||||||||||||||||||||||||||||||||||||||||||
R   ACTACTAGGTCTTCTCTTAGCCTCAGCAGGAATATCAGC---------------------------------------ACGGCGAT-CCTCAGTATCCGCACTCCTTCATTCCAGTACAATAGTTATAGCTGGAGTATTTACCCTCATCCGCTTTTATCCATTAATAGAAAACAACCTCAC
       |    |           | | | |||||||||||||                                       |||||||| |||||||||||||
P   TTCATCGTGCTGGTCAGATCCATGAACAGGAATATCAGC---------------------------------------ACGGCGAT-CCTCAGTATCCGCTAAAAGTGGTCGGTGAGACATCTACCACGGGTACTACTGTACGTTTCTGGCCAAGTGGGGATACTTTTAGCCAGAC
Pos         <-4595------------------------------------------------------------------4628->
R translated            ...S  R  n  I  S                                              t  a  i   l  S  I  r  T...
```

$\Delta G^0_{min}$ = -13.74 kcal/mol + -13.38 kcal/mol (average: -13.56 kcal/mol)

**MK144.**

```
                                                                                      |---| no net loss/gain
A   CTACTAGGTCTTCTCTTAGCCTCAGCAGGAAAATCAGCCCAATTTGGACTTCATCCGTGACTTCCAT-CAGCTAGTGAAGGCCCTACCCCAGTATCAGCACTCCTTCATTCCAGTACAATAGTTATAGCTGGAGTATTTACCCTCATCCGCTTTTATCCATTAATAGAAAACAACCTCAC
    ||||||||||||||||||||||||||||||||||||||||||||||||||||||||||||||||||||| |||| |   ||||||||||||||||||||||||||||||||||||||||||||||||||||||||||||||||||||||||||||||||||||||||||||||||||||||
R   CTACTAGGTCTTCTCTTAGCCTCAGCAGGAAAATCAGCCCAATTTGGACTTCATCCGTGACTTCCAT-CAGCGAACAAAGGCCCTACCCCAGTATCAGCACTCCTTCATTCCAGTACAATAGTTATAGCTGGAGTATTTACCCTCATCCGCTTTTATCCATTAATAGAAAACAACCTCAC
     |     |  |     | |  ||| | |  ||               ||         |      ||| |||||||||||||   | || |       || |   |     |                    ||       | |                |     | |
P   ATTGCTCGGTTTGCTCTGGAATCAACGGCCAAGCTTCTTGTGCAGTGAAGCTCAAGATAGGGCTCATCCAGCGAACAAAGGCTTGAACCAAATGATACAACGCAGTTTGTGCTGAATGGCGAGCTGCTGAATCTGCCTTGGTGGTGTACTGACGATCTTTAATAATATCGAGATAGAAAT
Pos                                                               27277<---------------->27294
R translated                                                      ...S  I   S  e  q  R  P...
```

$\Delta G^0_{min}$ = -10.86 kcal/mol

**MK67.**

```
                                                                                     |----| no net loss/gain
A   ACTACTAGGTCTTCTCTTAGCCTCAGCAGGAAAATCAGCCCAATTTGGACTTCATCCGTGACTTCCATCAGCTAGTGAAGGCCCTACCCCAGTATCAGCACTCCTTCATTCCAGTACAATAGTTATAGCTGGAGTATTTACCCTCATCCGCTTTTATCCATTAATAGAAAACAACCTCAC
    |||||||||||||||||||||||||||||||||||||||||||||||||||||||||||||||||||||| ||    ||||||||||||||||||||||||||||||||||||||||||||||||||||||||||||||||||||||||||||||||||||||||||||||||||||||
R   ACTACTAGGTCTTCTCTTAGCCTCAGCAGGAAAATCAGCCCAATTTGGACTTCATCCGTGACTTCCATCAGGTACATAAGGCCCTACCCCAGTATCAGCACTCCTTCATTCCAGTACAATAGTTATAGCTGGAGTATTTACCCTCATCCGCTTTTATCCATTAATAGAAAACAACCTCAC
      |   |    |||    || |||    |||   | |             |   |   | |  | ||||||||||||              | | ||| |   |        |||          |     |     |       | | |          |   |
P   TTTTAGAATAATTCATGCAGTCTCCTTCGGATTTTGATCTTTGGAGAGTTTCTTTTTCTTATCTTCATCAGGTACATAGTTGTTGGCAATAATCTCACCGAACGGCCCAGTCAGATTTGGTTGTGCCAGTCTTTCACGCGTTTTAAGCGGAAGCAGTGGAAATACCAGTTCAGCAAAACG
Pos                                                                40845<-----------> 40859
R translated                                                       ...S  I  r  y  I  R...
```

$\Delta G^0_{min}$ = -3.23 kcal/mol

**B11.**

```
                                                                                       |--------------------| net 9 bp loss
A   ACTACTAGGTCTTCTCTTAGCCTCAGCAGGAAAATCAGCCCAATTTGGACTTCATCCGTGACTTCCATCAGCTAGTGAAGGCCCTACCCCAGTATCAGCACTCCTTCATTCCAGTACAATAGTTATAGCTGGAGTATTTACCCTCATCCGCTTTTATCCATTAATAGAAAACAACCTCAC
    |||||||||||||||||||||||||||||||||||||||||||||||||||||||||||||||||||||||||  | |   |    ||||||  | |||||||||||||||||||||||||||||||||||||||||||||||||||||||||||||||||||||||||||||||||||||
R   ACTACTAGGTCTTCTCTTAGCCTCAGCAGGAAAATCAGCCCAATTTGGACTTCATCCGTGACTTCCATCAGCT--T-ATT-CA--ACCCCA--A-CAGCACTCCTTCATTCCAGTACAAAAGGTATAGCTGGAGTATTTACCCTCATCCGCTTTTATCCATTAATAGAAAACAACCTCAC
                         |  |          |  |       || |      | |  |  |||||||  | ||| ||  ||||||  | |||||||||            |    | |     | |        || |  |  ||   |     || ||          |
P   CACTGGTTCAAGAGGTAGGTACGGAAGTGACCTTGCGTCTGGTCAATGATTATGGGGCTTATGTAAATCAGCT--T-ATT-CA--ACCCCA--A-CAGCACTCCAATTAAGGTAGGCTGATGATGGTCAAGCATAGCCGCCCTTTGTGGGCACATTCATTTTTATCTGTGGTGATTCACT
Pos                                                                    <-121897----------------------121925->
R translated                                                         ...I  S  l     f   n    P  n     S  T  P...
```

$\Delta G^0_{min}$ = -7.55 kcal/mol + -12.65 kcal/mol (average: -10.10 kcal/mol)

```
R7.                                                                      |---------------------| net 3 bp loss
A  ACTACTAGGTCTTCTCTTAGCCTCAGCAGGAAAATCAGCCCAATTTGGACTTCATCCGTGACTTCCATCAGCTAGTGAAGGCCCTACCCCAGTATCAGCACTCCTTCATTCCAGTACAATAGTTATAGCTGGAGTATTTACCCTCATCCGCTTTTATCCATTAATAGAAAACAACCTCAC
R  ACTACTAGGTCTTCTCTTAGCCTCAGCAGGAAAATCAGCCCAATTTGGACTTCATCCGTGACTTCCATCACCCACTTCT---CATGTTGCAGCATCAGCACTCCTTCATTCCAGTACAATAGTTATAGCTGGAGTATTTACCCTCATCCGCTTTTATCCATTAATAGAAAACAACCTCAC
P  CATTACTTAATCATCAATTATCTGGATTTGGTGCAGACGTTTTGGATCAGGAACCACCAGCAAAAGATCACCCACTTCT---CATGTTGCAGCATCAGCACCCAAATGTTCTGATTACTGGGCATATTGCTTGGGGAACAGAAGAGGCTCAACAGCGTTTATTTGATATTTTGCAAGATA
Pos                                                                  <-137928-------------------137897->
R translated                                                      ...I  t  h  f  s     c  c  S  I  S  T...
```

$\Delta G^0_{min}$ = -2.89 kcal/mol + -13.67 kcal/mol (average: -8.33 kcal/mol)

```
MK146.                                                                      |-| no net loss/gain
A  ACTACTAGGTCTTCTCTTAGCCTCAGCAGGAAAATCAGCCCAATTTGGACTTCATCCGTGACTTCCATCAGCTAGTGAAGGCCCTACCCCAGTATCAGCACTCCTTCATTCCAGTACAATAGTTATAGCTGGAGTATTTACCCTCATCCGCTTTTATCCATTAATAGAAAACAACCTCAC
R  ACTACTAGGTCTTCTCTTAGCCTCAGCAGGAAAATCAGCCCAATTTGGACTTCATCCGTGACTTCCATCAGCTGGGGAAGGCCCTACCCCAGTATCAGCACTCCTTCATTCCAGTACAATAGTTATAGCTGGAGTATTTACCCTCATCCGCTTTTATCCATTAATAGAAAACAACCTCAC
P  AGAAAAAAATGATTATAATAAATGGGTGAAGAAAATAAAAAATAATTGTGAAGGAAAAATAAGTCATATAGCTGGGGAAGGTGCAGGTCTAGTTAGAAATCAATGTTATAGAGATGAGTATTTATCTAGATTATCATTTTTAACAACTGGAAAAACTTCGAAAGAAGAAAATATTCATGG
Pos                                                               172627<---------->172644
R translated                                                         ...S  w  g  R...
```

$\Delta G^0_{min}$ = -11.45 kcal/mol

```
MK82.                           |-- net 87 bp loss --------------------------------------------------------------------------------------------|
A  ACTACTAGGTCTTCTCTTAGCCTCAGCAGGAAAATCAGCCCAATTTGGACTTCATCCGTGACTTCCATCAGCTAGTGAAGGCCCTACCCCAGTATCAGCACTCCTTCATTCCAGTACAATAGTTATAGCTGGAGTATTTACCCTCATCCGCTTTTATCCATTAATAGAAAACAACCTCAC
R  ACTACTAGGTCTTCTCTTAGCCTCAGCAGTGCAAATAAGTCCGATAGCGCCCGAA---------------------------------------------------------------------------------------CTCATCCGCTTTTATCCATTAATAGAAAACAACCTCAC
P  TTATTGAAATTTCTGGAAAATGTCAGCAGTGCAAATAAGTCCGATAGCGCCCGAA---------------------------------------------------------------------------------------CTCATCCGCCTTGAAAATATTTATGTTCATCGCGACGA
Pos                      <-176930--------------------------------------------------------------------------------------------------------------------176880->
R translated         ...L  S  s  a  n  k  s  d  s  a  r  t                                                                                         H  P  L  L...
```

$\Delta G^0_{min}$ = -8.42 kcal/mol + -15.04 kcal/mol (average: -11.73 kcal/mol)

```
MK64.                                                                   |- net 12 bp inserted |
A  ACTACTAGGTCTTCTCTTAGCCTCAGCAGGAAAATCAGCCCAATTTGGACTTCATCCGTGACTTCCATC--AGCT-AGT---GA--A-G---GCCTACCCCAGTATCAGCACTCCTTCATTCCAGTACAATAGTTATAGCTGGAGTATTTACCCTCATCCGCTTTTATCCATTAATAGAAA
R  ACTACTAGGTCTTCTCTTAGCCTCAGCAGGAAAATCAGCCCAATTTGGACTTCATCCGTGACTTCCATCGCAGTTAATTCCCTATCAGGATCGCCTACCCCAGTATCAGCACTCCTTCATTCCAGTACAATAGTTATAGCTGGAGTATTTACCCTCATCCGCTTTTATCCATTAATAGAAA
P  AAAGATGCTTGCTCAGCTTCAAACCCATACCGATTTTATTGTGACACCCGATGATCCCGCTTATCCATCGCAGTTAATTCCCTATCAGGATCGCCCTCCCATTCTATTTGGTCAAGGCTCAATACAAAGCCTGATCCAACCACAAATTGCAGTCGTGGGTAGTCGTAAACCCAGCCCTCAT
Pos                                                               <-205715----------------205679->
R translated                                                   ...S  I  a  v  n  s  l  s  g  s  P...
```

$\Delta G^0_{min}$ = -5.72 kcal/mol + -4.60 kcal/mol (average: -5.16 kcal/mol)

```
A78.                                                                       |---------------| net 12 bp loss
A  ACTACTAGGTCTTCTCTTAGCCTCAGCAGGAAAATCAGCCCAATTTGGACTTCATCCGTGACTTCCATCAGCTAGTGAAGGCCCTACCCCAGTATCAGCACTCCTTCATTCCAGTACAATAGTTATAGCTGGAGTATTTACCCTCATCCGCTTTTATCCATTAATAGAAAACAACCTCAC
R  ACTACTAGGTCTTCTCTTAGCCTCAGCAGGAAAATCAGCCCAATTTGGACTTCATCCGTGACTTCCATCAGCCAGTA------------CAGTATCAACACTCCTTCATTCCAGTACAATAGTTATAGCTGGAGTATTTACCCTCATCCGCTTTTATCCATTAATAGAAAACAACCTCAC
P  CTTCGGTTGTACAATCTCACTCAAATCATCAACATCTGGCAAAGCCCGTACATTTAACTGGCAATCATCAGCCAGTA------------CAGTATCAACACCTGTAATGATTGCGCCAGAACGTGCTCGCCAGTATTGCACATCGAGCCTTGCATCTACACCAGTAATCCACTTGGATTC
Pos                                                                 <-247271--------------------247248->
R translated                                                      ...I  S  q  y              S  I  n  T...
```

$\Delta G^0_{min}$ = -11.59 kcal/mol + -12.31 kcal/mol (average: -11.95 kcal/mol)

```
MK5.                                                                        |-----| no net gain/loss
A  ACTACTAGGTCTTCTCTTAGCCTCAGCAGGAAAATCAGCCCAATTTGGACTTCATCCGTGACTTCCATCAGCTAGTGAAGGCCCTACCCCAGTATCAGCACTCCTTCATTCCAGTACAATAGTTATAGCTGGAGTATTTACCCTCATCCGCTTTTATCCATTAATAGAAAACAACCTCAC
R  ACTACTAGGTCTTCTCTTAGCCTCAGCAGGAAAATCAGCCCAATTTGGACTTCATCCGTGACTTCCATCAGCTTTCCAATGCCCTACCCCAGTATCAGCACTCCTTCATTCCAGTACAATAGTTATAGCTGGAGTATTTACCCTCATCCGCTTTTATCCATTAATAGAAAACAACCTCAC
P  AGCCTTATCGTTTGGCACGGGCAATGTCTTTCACCAATCCATGGGCAGATCCGTTAGGCACAGGTTATCAGCTTTCCAATGCCCTGATGGCCTTTGGGCGAGGTGAATGGTTTGGCACAGGTTTAGGTCACAGTGTTCAAAAACTCTCGTATTTACCCGAAGCGCATACCGACTTTATGT
Pos                                                                  <-272208---272226->
R translated                                                      ...I  S  f  p  m  P  Y...
```

$\Delta G^0_{min}$ = -13.32 kcal/mol

**MK69.**

```
                                          |-- net 42 bp loss ------------------------------------|
A  ACTACTAGGTCTTCTCTTAGCCTCAGCAGGAAAATCAGCCCAATTTGGACTTCATCCGTGACTTCCATCAGCTAGTGAAGGCCCTACCCCAGTATCAGCACTCCTTCATTCCAGTACAATAGTTATAGCTGGAGTATTTACCCTCATCCGCTTTTATCCATTAATAGAAAACAACCTCAC
R  ACTACTAGGTCTTCTCTTAGCCTCAGCAGGAAAATCAGACCA------------------------------------------GAAACCAGCATCAGCACTCCTTCATTCCAGTACAATAGTTATAGCTGGAGTATTTACCCTCATCCGCTTTTATCCATTAATAGAAAACAACCTCAC
P  AAGATTTTATAAATGGTTCTTGCCATAGAAGGCCCCAGACCA------------------------------------------GAAACCAGCATCAGCTCCTTTTCCGAAGCTTTGAGTACACCCTGGATACCACCAAAATGAGTCAATAAGTCACGACGACGCTTAGGCCCTAATCCT
Pos                                       <-337614---------------------------------------------------337593->
R translated                           ...I  s  P                                              e  t  S  I  S...
```

$\Delta G^0_{min}$ = -6.72 kcal/mol + -12.36 kcal/mol  (average: -9.54 kcal/mol)

**K148.**

```
                                                                            |--| no net loss/gain
A  ACTACTAGGTCTTCTCTTAGCCTCAGCAGGAAAATCAGCCCAATTTGGACTTCATCCGTGACTTCCATCAGCTAGTGAAGGCCCTACCCCAGTATCAGCACTCCTTCATTCCAGTACAATAGTTATAGCTGGAGTATTTACCCTCATCCGCTTTTATCCATTAATAGAAAACAACCTCAC
R  ACTACTAGGTCTTCTCTTAGCCTCAGCAGGAAAATCAGCCCAATTTGGACTTCATCCGTGACTTCCATCAGCTTTTCAAGGCCCTACCCCAGTATCAGCACTCCTTCATTCCAGTACAATAGTTATAGCTGGAGTATTTACCCTCATCCGCTTTTATCCATTAATAGAAAACAACCTCAC
P  TCTTTTTCATTGAGTACAGGATGAGGGCGCACTGCAGCAACAACGCCCTGATGAAAAGGAAGGCCAGCCAGCTTTTCAAGACTGTCACGACTCGCCTTTTGTACACTGATACCAAAAGGTTCCGCCAGCTCCAGAATCTTTTGCAAGCGCTGGTCATCACGACCTTTTAAAGTAAACAGT
Pos                                                                  350408<------------>350394
R translated                                                          ...I  S  f  s  R  P...
```

$\Delta G^0_{min}$ = -7.02 kcal/mol

**R1523.**

```
                                           |-- net 39 bp loss ------------------------------------|
A  ACTACTAGGTCTTCTCTTAGCCTCAGCAGGAAAATCAGCCCAATTTGGACTTCATCCGTGACTTCCATCAGCTAGTGAAGGCCCTACCCCAGTATCAGCACTCCTTCATTCCAGTACAATAGTTATAGCTGGAGTATTTACCCTCATCCGCTTTTATCCATTAATAGAAAACAACCTCAC
R  ACTACTAGGTCTTCTCTTAGCCTCAGCAGGAAAATCAGCGCAATC---AAT--ATCC---------------------AGG-------------TCAGCACTCCTTCATTCCAGTACAATAGTTATAGCTGGAGTATTTACCCTCATCCGCTTTTATCCATTAATAGAAAACAACCTCAC
P  TCAGGTAGAAGCTGGGTATCAAAGTCAATAAAGGTCAGCGCAATC---AAT--ATCC---------------------AGG-------------TCAGCACTAAACCGAAAAGCATGTGTAGTGTCGCACCAAATACCACAACCACCAGCAGCGAACAAAGCATGGTCAGTAACTCAACC
Pos                                   <-352932---------------------------------------------------352960->
R translated                       ...I  S  a  I     n    i  q                       v              S  T...
```

$\Delta G^0_{min}$ = -11.27 kcal/mol + -10.22 kcal/mol  (average: -10.75 kcal/mol)

**R18.**

```
                                                                         |-----| no net gain/loss
A  ACTACTAGGTCTTCTCTTAGCCTCAGCAGGAAAATCAGCCCAATTTGGACTTCATCCGTGACTTCCATCAGCTAGTGAAGGCCCTACCCCAGTATCAGCACTCCTTCATTCCAGTACAATAGTTATAGCTGGAGTATTTACCCTCATCCGCTTTTATCCATTAATAGAAAACAACCTCAC
R  ACTACTAGGTCTTCTCTTAGCCTCAGCAGGAAAATCAGCCCAATTTGGACTTCATCCGTGACTTCCATCATGGAGGTAAGGCCCTACCCCAGTATCAGCACTCCTTCATTCCAGTACAATAGTTATAGCTGGAGTATTTACCCTCATCCGCTTTTATCCATTAATAGAAAACAACCTCAC
P  GGAAGATACATTGCAATAACCAGTCCACCGACTAAAATGCCCAAAATCGCCATAATCAGCGGTTCCATCATGGAGGTAAGGCCATCGACCGCATGATCGACTTCACTTTCATAATAATTTGCGACTTTATCCAGCATTTCATCGAGTGCTCCAGATTCTTCACCAATCGCTACCATTTGG
Pos                                                             <-353463-----353483->
R translated                                                 ...T  S  I  m  e  v  R  P...
```

$\Delta G^0_{min}$ = -17.64 kcal/mol

**MK125.**

```
                                                  |-- net 6 bp loss ---------------------------|
A  ACTACTAGGTCTTCTCTTAGCCTCAGCAGGAAAATCAGCCCAATTTGGACTTCATCCGTGACTTCCATCAGCTAG-TGAAGGCC-CTACCCCAGTATCAGCACTCCTTCATTCCAGTACAATAGTTATAGCTGGAGTATTTACCCTCATCCGCTTTTATCCATTAATAGAAAACAACCTCAC
R  ACTACTAGGTCTTCTCTTAGCCTCAGCAGGAAAATCAGCCCAATTTGGACTTCA-CC----CTT--ACCAGCCAGCTCAATACCACT-CACCAGAACCAACACTCCTTCATTCCAGTACAATAGTTATAGCTGGAGTATTTACCCTCATCCGCTTTTATCCATTAATAGAAAACAACCTCAC
P  TTAACTCTCAAATAACGTGGGTCAACCCATTCTCTCAACTGTTTAACTGATTCA-CC----CTT--ACCAGCCAGCTCAATACCACT-CACCAGAACCAACACACTTTTGGCATTTTTCTTTCCTGCACTATTTTTAAGCTGGTATTGTGCTTCGATCAAATCCAGGGATAAATGATCATGA
Pos                                                <-386930-------------------------------------386886->
R translated                                    ...T  S  p     l   p  a  s  s  i  P  l   t  r  t  n   T...
```

$\Delta G^0_{min}$ = -3.33 kcal/mol + -12.06 kcal/mol  (average: -7.70 kcal/mol)

**MK75.**

```
                                                                              |--| no net loss/gain
A  ACTACTAGGTCTTCTCTTAGCCTCAGCAGGAAAATCAGCCCAATTTGGACTTCATCCGTGACTTCCATCAGCTAGTGAAGGCCCTACCCCAGTATCAGCACTCCTTCATTCCAGTACAATAGTTATAGCTGGAGTATTTACCCTCATCCGCTTTTATCCATTAATAGAAAACAACCTCAC
R  ACTACTAGGTCTTCTCTTAGCCTCAGCAGGAAAATCAGCCCAATTTGGACTTCATCCGTGACTTCCATCAGCTACTCCAGGCCCTACCCCAGTATCAGCACTCCTTCATTCCAGTACAATAGTTATAGCTGGAGTATTTACCCTCATCCGCTTTTATCCATTAATAGAAAACAACCTCAC
P  TACAGCGTCAGGCACCAGCCGTTCCTTTCAGCGCAATTCAACAGCAAGTGGAAAAAGAGCTCGGTAAACCGCTACTCCAGGCCTTTCAGTCTTTTGATCAACAACCTTTTGCCGCAGCATCAATTGGTCAGGTACATCGGGCTGTTTTGCCAGACGGACAAGAAGTTGTGGTCAAAGTAC
Pos                                                                    424549<----------->424537
R translated                                                             ...S  y  s  R  P...
```

$\Delta G^0_{min}$ = -10.64 kcal/mol

**O131.**

```
                                                                                   |---| no net gain/loss
A  ACTACTAGGTCTTCTCTTAGCCTCAGCAGGAAAATCAGCCCAATTTGGACTTCATCCGTGACTTCCATCAGCTAGTGAAGGCCCTACCCCAGTATCAGCACTCCTTCATTCCAGTACAATAGTTATAGCTGGAGTATTTACCCTCATCCGCTTTTATCCATTAATAGAAAACAACCTCAC
R  ACTACTAGGTCTTCTCTTAGCCTCAGCAGGAAAATCAGCCCAATTTGGACTTCATCCGTGACTTCCATCAGCAACTTAAGGCCCTACCCCAGTATCAGCACTCCTTCATTCCAGTACAATAGTTATAGCTGGAGTATTTACCCTCATCCGCTTTTATCCATTAATAGAAAACAACCTCAC
P  ATGTGTTACCTAATTTTCGTGCGCATCATCCAGAACTTAGAATTGAACTCCATAGTTTAAATAACGATCAGCAACTTAAGGCGCTTAAACGTGGTGAACTCGATTTATGTTTTATCCGAGATAATATTCAGCATCGTGATGGTATCCATAGTCAATTGGTATTAAGTGAACCACTCATTT
Pos                                                                  <-444536---444518->
R translated                                                     ...S  I  S  n  l  R  P  Y...
```

$\Delta G^0_{min}$ = -16.89 kcal/mol

**K52.**

```
                                                                                 |----------| no net gain/loss
A  ACTACTAGGTCTTCTCTTAGCCTCAGCAGGAAAATCAGCCCAATTTGGACTTCATCCGTGACTTCCATCAGCTAGTGAAGGCCCTACCCCAGTATCAGCACTCCTTCATTCCAGTACAATAGTTATAGCTGGAGTATTTACCCTCATCCGCTTTTATCCATTAATAGAAAACAACCTCAC
R  ACTACTAGGTCTTCTCTTAGCCTCAGCAGGAAAATCAGCCCAATTTGGACTTCATCCGTGACTTCCATCGGCAAATCAGCACCCTACCCCAGTATCAGCACTCCTTCATTCCAGTACAATAGTTATAGCTGGAGTATTTACCCTCATCCGCTTTTATCCATTAATAGAAAACAACCTCAC
P  AAGCTTTCTTAAAATCATTGCTGTTCCAAACACAAATAATTTATACGGATTTCTCATGACAACGACATCGGCAAATCAGCACCCTACCCGTCGCGATCTTCTAAAATGGTTTACTGGTATTCCTTTTTTACCACTCGGGGCAATGGCAACAGCAGCAACACTGGCAGGATGCAATGACAG
Pos                                                                <-508476--------508453->
R translated                                                     ...S  I  g  k  s  a  P  Y
```

$\Delta G^0_{min}$ = -17.32 kcal/mol

**MK59.**

```
                                                 |-- net 93 bp loss ---------------------------------------------------------------------------------|
A  ACTACTAGGTCTTCTCTTAGCCTCAGCAGGAAAATCAGCCCAATTTGGACTTCATCCGTGACTTCCATCAGCTAGTGAAGGCCCTACCCCAGTATCAGCACTCCTTCATTCCAGTACAATAGTTATAGCTGGAGTATTTACCCTCATCCGCTTTTATCCATTAATAGAAAACAACCTCAC
R  ACTACTAGGTCTTCTCTTAGCCTCAGCAGGAAAATCAGCCCAATAG-----------------------------------------------------------------------------------------AATTTAATCCGCTATTATCCATTAATAGAAAACAACCTCAC
P  GTTGCTGCATTTTTATTTTTAAGTCTAAATGGTCGAAGCCCAATAG-----------------------------------------------------------------------------------------AATTTAATCCGCTATGTTGTTATTTCAGAACAGTGTTTGAC
Pos                                      <-511159---------------------------------------------------------------------------------------------511183->
R translated                         ...S  P  I  e                                                                                       f  n  P  L  L...
```

$\Delta G^0_{min}$ = -10.38 kcal/mol + -8.04 kcal/mol (average: -9.21 kcal/mol)

**MK13.**

```
                                                                                   |-- net 12 bp loss ----|
A  ACTACTAGGTCTTCTCTTAGCCTCAGCAGGAAAATCAGCCCAATTTGGACTTCATCCGTGACTTCCATCAGCTAGTGAAGGCCCTACCCCAGTATCAGCACTCCTTCATTCCAGTACAATAGTTATAGCTGGAGTATTTACCCTCATCCGCTTTTATCCATTAATAGAAAACAACCTCAC
R  ACTACTAGGTCTTCTCTTAGCCTCAGCAGGAAAATCAGCCCAATTTGGACTTCATCCGTGACTTCCATCAGCAAGT------CC-A-TGC--TA--AGCACTCCTTCATTCCAGTACAATAGTTATAGCTGGAGTATTTACCCTCATCCGCTTTTATCCATTAATAGAAAACAACCTCAC
P  AAATATGCGGCACGGGTTTTTGACTACGTGCAGTCACACGTTCGGTCACTTTTACTAATATCGGATTCAGCAAGT------CC-A-TGC--TA--AGCACTCCTTTTACAACTGGATAAGCTGGCAAACAAAATAGACCAAATGATTTCATGATCATTTGAAAAAAATAAATAATATTA
Pos                                                                  <-532435----------------------532409->
R translated                                                       ...I  S  f  s        m   l      S  T  P  S...
```

$\Delta G^0_{min}$ = -7.40 kcal/mol + -15.36 kcal/mol (average: -11.38 kcal/mol)

**MK104.**

```
                                                                   |-- net 75 bp loss -------------------------------------------------------------------------------------|
A  ACTACTAGGTCTTCTCTTAGCCTCAGCAGGAAAATCAGCCCAATTTGGACTTCATCCGTGACTTCCATCAGCTAGTGAAGGCCCTACCCCAGTATCAGCACTCCTTCATTCCAGTACAATAGTTATAGCTGGAGTATTTACCCTCATCCGCTTTTATCCATTAATAGAAAACAACCTCAC
R  ACTACTAGGTCTTCTCTTAGCCTCAGCAGGAAAATCAGCCCAATTTGGACTTCATCCGGGACAGTGCAACCAGTA--------------------------------------------------------------------------AAGTAAATGGCAACATCAAAACAACCTCAC
P  CACCAATAAATGCACCAACCAGAACATCACTTGGGTAATGTAGCCCCAAAATCATCCGGGACAGTGCAACCAGTA--------------------------------------------------------------------------AAGTAAATGGCAACATCAAAACAACCAATA
Pos                                                        <-548165---------------------------------------------------------------------------------------------------------548116->
R translated                                                                                                                                                  m  a  t  s  K  Q  P...
```

$\Delta G^0_{min}$ = -13.46 kcal/mol + -10.55 kcal/mol (average: -12.01 kcal/mol)

**MK124.**

```
                                                                                |-----------------| no net loss/gain
A  ACTACTAGGTCTTCTCTTAGCCTCAGCAGGAAAATCAGCCCAATTTGGACTTCATCCGTGACTTCCATCAGCTAG-TGAAGGCCCTACCCCAGTATCAGCACTCCTTCATTCCAGTACAATAGTTATAGCTGGAGTATTTACCCTCATCCGCTTTTATCCATTAATAGAAAACAAC
R  ACTACTAGGTCTTCTCTTAGCCTCAGCAGGAAAATCAGCCCAATTTGGACTTCATCCGTGACTTCCATAACCCAGCTCAATCCCCC-CCCCAGTATCAGCACTCCTTCATTCCAGTACAATAGTTATAGCTGGAGTATTTACCCTCATCCGCTTTTATCCATTAATAGAAAACAAC
P  TTTTGATCCATCTGTTTCGGCCTGTTCATCAAAATAGTGAATAAAAACCAGGTCCAGATCAGAACCATAACCCAGCTCAATCCCCC-CCAGCTTCCCATAACCAATCACTGCAAAACCTGTATGCTCCAGTGAGCAACGTTCACCACTGGCATCTTTGGGATAACCATGTTTTTTA
Pos                                                           586394<------------------->586368
R translated                                                     ...S  I  t  q  l  n  p  p   P...
```

$\Delta G^0_{min}$ = -12.27 kcal/mol

**K109.**

```
                                                                       |------------------| no net gain/loss
A  ACTACTAGGTCTTCTCTTAGCCTCAGCAGGAAAATCAGCCCAATTTGGACTTCATCCGTGACTTCCATCAGCTAGTGA--AGGCCCTACCCCAGTATCAGCACTCCTTCATTCCAGTACAATAGTTATAGCTGGAGTATTTACCCTCATCCGCTTTTATCCATTAATAGAAAACAACCTC
R  ACTACTAGGTCTTCTCTTAGCCTCAGCAGGAAAATCAGCCCAATTTGGACTTCATCCGTGAC-GACATCAGAT-TTGATCACGCCCAACCCCAGTATCAGCACTCCTTCATTCCAGTACAATAGTTATAGCTGGAGTATTTACCCTCATCCGCTTTTATCCATTAATAGAAAACAACCTC
P  TGAGCTTCAAGTTCAGCCAGTTCTTCAGGCGTTGGTACATGAATACGTGATAGATCCGTGAC-GACATCAGAT-TTGATCACGCCCAACATGTTGACGAAGAGATTAAATGCTTCTTTTTTGTATTCTTGCTCTGGATTCTTCTGTGCGTAGCCACGTAAATGAATTCCCTGACGTAAGT
Pos                                                   <-639281--------------------639248->
R translated                                   ...S  S  V  T  t  s  d  l  i  t  P  n  P...
```

$\Delta G^0_{min}$ = -24.65 kcal/mol

**MK109.**

```
                                                                                                                      |-- net 9 bp inserted -------------|
A  CCTCAGCAGGAAAATCAGCCCAATTTGGACTTCATCCGTGACTTCCATCAGCTAGTGAAGGCCCTACCCCAGTATCAGCACTCCTTCATTCCAGTACAATAGTTATAGCTGGAGTATTTAC-CC------T-CATCCG-CTTTTATCCATTAATAGAAAACAACCTCACTATTCAAACTT
R  CCTCAGCAGGAAAATCAGCCCAATTTGGACTTCATCCGTGACTTCCATCAGCTAGTGAAGGCCCTACCCCAGTATCAGCACTCCTTCATTCCAGTACAATAGTTATAGCTGGAGTTTTACACCCAAGGTTGTATAAGAGGTCTAGCCATGAATAGAAAACAACCTCACTATTCAAACTT
P  TTGTTCCTATTTAATTTTTTGATCTTTAAAGAGGCTAGAGATTAGCCTCTTTTTTTATGTATAAAATGTAGCGACGTACTAAATTATCTCTATGATTTATAGTAGAATGATGAGTTTTTACACCCAAGGTTGTATAAGAGGTCTAGCCATGAATAGTGAAC-AGCTCACACAAATTTTAC
Pos                                                                                                                  <-653745-----------------------------------------653801->
R translated                                                                                                                                               m  N  R  K  Q  P  H...
```

$\Delta G^0_{min}$ = -8.47 kcal/mol + -20.53 kcal/mol (average: -14.50 kcal/mol)

**MK123.**

```
                                                             |-- net 105 bp loss -------------------------------------------------------------------------------------------|
A  TCTTAGCCTCAGCAGGAAAATCAGCCCAATTTGGACTTCATCCGTGACTTCCATCAGCTAGTGAAGGCCCTACCCCAGTATCAGCACTCCTTCATTCCAGTACAATAGTTATAGCTGGAGTATTTACCCTCATCCGCTTTTATCCATTAATAGAAAACAACCTCACTATTCAAACTTCAACACT
R  TCTTAGCCTCAGCAGGAAAATCAGCCCAATTTGGACTTCATCCGTGACTTCTG-----------------------------------------------------------------------------------------------------CACCCATTTCTTCAAACTTCAACACT
P  ACGTTTGCCCATCATATCCAATTCGATCCAGTCTTCACCACGCGTGACTTCTG-----------------------------------------------------------------------------------------------------CACCCATTTCTTCAAACTTATCAAGA
Pos                                              <-654868-------------------------------------------------------------------------------------------------------------654835->
R translated                                 ...S  S  V  T  S  a                                                                                                     p  i  s  S  N  F...
```

$\Delta G^0_{min}$ = -13.93 kcal/mol + -9.82 kcal/mol (average: -11.88 kcal/mol)

**MK98.**

```
                                                     |-- net 6 bp loss --------------------------|
A  ACTACTAGGTCTTCTCTTAGCCTCAGCAGGAAAATCAGCCCAATTTGGACTTCA-TCCGTGACTTCCATCAGCTAGTGAAGGCCCTACCC-CAGTATCAGCACTCCTTCATTCCAGTACAATAGTTATAGCTGGAGTATTTACCCTCATCCGCTTTTATCCATTAATAGAAAACAACCTC
R  ACTACTAGGTCTTCTCTTAGCCTCAGCAGGAAAATCAGCACTCCTT-CATTCCAGTACAATAGTT--AT-AGCT-G-G-AGTATTTACCC-CAGTATCAGCACTCCTTCATTCCAGTACAATAGTTATAGCTGGAGTATTTACCCTCATCCGCTTTTATCCATTAATAGAAAACAACCTC
ND ACTTCCATCAGCTAGTGAAGGCCCTACCCCAGTATCAGCACTCCTT-CATTCCAGTACAATAGTT--AT-AGCT-G-G-AGTATTTACCCTCATCCGCTTTTATCCATTAATAGAAAACAACCTCACTATTCAAACTTCAACACTTCTGGTTCTGGCTCTAATGTCACTAATGAAAAAAT
Pos                                   <-109 of 'ND5i' insert -----------------------------161->
R translated                      ...K  I  S  t  p  s  f  q  y  n  s  y  S  w  s  I  Y  P  S  I  S  T  P  S  F  Q  Y  N  S  Y  S  W  S  I  Y  P  H...
```

$\Delta G^0_{min}$ = -6.84 kcal/mol + -6.04 kcal/mol (average: -6.44 kcal/mol)

Note: the templating DNA originated from sequence downstream of stop codon, inside 'ND5i' DNA (orange). Resulting in a direct repeat *in frame*, separated by two nucleotides.

**MK116.**

```
                                                          |-- net 12 bp loss ---------------------------------------------------------------|
A  ACTACTAGGTCTTCTCTTAGCCTCAGCAGGAAAATCAGCCCAATTTGGACTTCATCCGTGACT-TCCATCAGCTAGTGAAGGCCCTACCCCAGTAT-CAGCACTCCTTCATTCCAGTACAATAGTTATAGCTGGAGTATTTACCCTCATCCGCTTTTATCCATTAATAGAAAACAACCTCA
R  ACTACTAGGTCTTCTCTTAGCCTCAGCAGGAAAATCAGCCCAATTTACCCT-CATCCGCTTTTATCCATTAA-TAGAAAA--CA--ACCTCACTATTCAA-ACT--T-CAA-C-ACTTC--TGGTTCTGGCTGGAGTATTTACCCTCATCCGCTTTTATCCATTAATAGAAAACAACCTCA
ND ATCAGCACTCCTTCATTCCAGTACAATAGTTATAGCTGGAGTATTTACCCT-CATCCGCTTTTATCCATTAA-TAGAAAA--CA--ACCTCACTATTCAA-ACT--T-CAA-C-ACTTC--TGGTTCTGGCTCTAATGTCACTAATGAAAAAATGCGTTTCTGGAGTCCAGAGGTTCGTGA
Pos                                             <-151 of 'ND5i' ----------------------------------------------------------------226->
R translated                                ...I  y  p  h  p  l  l  s  i  n  r  k  q  p  h  y  s  n  f  n  t  s  g  s  g  W...
```

$\Delta G^0_{min}$ = -10.25 kcal/mol + -4.15 kcal/mol (average: -7.20 kcal/mol)

**K49.**

```
                                                                                      |----| no net loss/gain
A  ACTACTAGGTCTTCTCTTAGCCTCAGCAGGAAAATCAGCCCAATTTGGACTTCATCCGTGACTTCCATCAGCTAGTGAAGGCCCTACCCCAGTATCAGCACTCCTTCATTCCAGTACAATAGTTATAGCTGGAGTATTTACCCTCATCCGCTTTTATCCATTAATAGAAAACAACCTCAC
R  ACTACTAGGTCTTCTCTTAGCCTCAGCAGGAAAATCAGCCCAATTTGGACTTCATCCGTGACTTCCATCAGGTACATAAGGCCCTACCCCAGTATCAGCACTCCTTCATTCCAGTACAATAGTTATAGCTGGAGTATTTACCCTCATCCGCTTTTATCCATTAATAGAAAACAACCTCAC
P  AGGCGAAGGTGGATACGGATTTTCATTGGTATTGAGTTTAAGTAAGTTCTGGATTTTAGGTTGCTCACCAGGTACATAAGGCTCTAACTCACGAACCTCTGGACTCCAGAAACGCATTTTTTCATTAGTGACATTCATCTTTCTATTCTCAAAAAAGGGCTGAAAATCAGCCCCGTAGAG
Poa                                                                 <-657707-----657687->
R translated                                                     ...S  I  r  y  i  R  P  Y...
```

$\Delta G^0_{min}$ = -12.55 kcal/mol

```
A4.                             |-- net 42 bp loss ---------------------------------|
A  ACTACTAGGTCTTCTCTTAGCCTCAGCAGGAAAATCAGCCCAATTTGGACTTCATCCGTGACTTCCATCAGCTAGTGAAGGCCCTACCCCAGTATCAGCACTCCTTCATTCCAGTACAATAGTTATAGCTGGAGTATTTACCCTCATCCGCTTTTATCCATTAATAGAAAACAACCTCAC
R  ACTACTAGGTCTTCTCTTAGCCTCACCAGG------------------------------------------TACATAAGGCCCTACCCCAGTATCAGCACTCCTTCATTCCAGTACAATAGTTATAGCTGGAGTATTTACCCTCATCCGCTTTTATCCATTAATAGAAAACAACCTCAC
P  TAAGTTCTGGATTTAGGTTGCTCACCAGG------------------------------------------TACATAAGGCTCTAACTCACGAACCTCTGGACTCCAGAAACGCATTTTTTCATTAGTGACATTCATCTTTCTATTCTCAAAAAAGGGCTGAAAATCAGCCCCGTAGAG
Pos                     <-657709-----------------------------------------------657687->
R translated         ...L  t  R                                          y  i  R  P  Y...
```

$\Delta G^0_{min}$ = -9.21 kcal/mol + -8.64 kcal/mol  (average: -8.93 kcal/mol)

```
MK2.                                                                    |-- net 6 bp loss ------------|
A  ACTACTAGGTCTTCTCTTAGCCTCAGCAGGAAAATCAGCCCAATTTGGACTTCATCCGTGACTTCCATCAGCTAGTGAAGGCCCTACCCCAGTATCAGCACTCCTTCATTCCAGTACAATAGTTATAGCTGGAGTATTTACCCTCATCCGCTTTTATCCATTAATAGAAAACAACCTCACT
R  ACTACTAGGTCTTCTCTTAGCCTCAGCAGGAAAATCAGCCCAATTTGGACTTCATCCGTGACTTCCATCAGC-ATTCTTGCCGTTAACCCC----CA-CACCACTTCATTCCAGTACAATAGTTATAGCTGGAGTATTTACCCTCATCCGCTTTTATCCATTAATAGAAAACAACCTCACT
P  AATGAAGGCGGATTCAGTAAAAATGGCCACTCTTTCTGATAACCCAATCCATAATTAAATAAAGAAATCAGC-ATTCTTGCCGTTAACCCC----CA-CACCACTTCATCTTCGACCTGTAAACTTGGGAAATAAAGCGTTTGTTTGATGTATTGTACTTTGTAAGGGGTTGGTGGCGCGT
Pos                                                                 <-669516----------------------------669480->
R translated                                                     ...I  S  t  l  a  v  n  p     h  t  t  S  F...
```

$\Delta G^0_{min}$ = -5.84 kcal/mol + -8.81 kcal/mol  (average: -7.33 kcal/mol)

```
MK72.                                                                   |---| no net loss/gain
A  ACTACTAGGTCTTCTCTTAGCCTCAGCAGGAAAATCAGCCCAATTTGGACTTCATCCGTGACTTCCATCAGCTAGTGAAGGCCCTACCCCAGTATCAGCACTCCTTCATTCCAGTACAATAGTTATAGCTGGAGTATTTACCCTCATCCGCTTTTATCCATTAATAGAAAACAACCTCAC
R  ACTACTAGGTCTTCTCTTAGCCTCAGCAGGAAAATCAGCCCAATTTGGACTTCATCCGTGACTTCCATCAGCCAGAAAAGGCCCTACCCCAGTATCAGCACTCCTTCATTCCAGTACAATAGTTATAGCTGGAGTATTTACCCTCATCCGCTTTTATCCATTAATAGAAAACAACCTCAC
P  TAAAATAGCAGGCACCAACATTAGGCGTGGGTTGTTGGTTCGCTGTGACAGAAAAATACCAACGATCAAAGCCAGAAAAGGCCATAACGCGTAGCTCACCAGACCTTGAGAGTTAAATTGTCCGCCATCCCCTGCAACCATCCAGCTAAACAGTGTATTGGCAGCCCCCCCTAATACTCC
Pos                                                           672958<------------>672945
R translated                                                    ...S  q  k  R  P...
```

$\Delta G^0_{min}$ = -11.55 kcal/mol

```
MK44.                                                          |-----------| no net loss/gain
A  ACTACTAGGTCTTCTCTTAGCCTCAGCAGGAAAATCAGCCCAATTTGGACTTCATCCGTGACTTCCATCAGCTAGTGAAGGCCCTACCCCAGTATCAGCACTCCTTCATTCCAGTACAATAGTTATAGCTGGAGTATTTACCCTCATCCGCTTTTATCCATTAATAGAAAACAACCTCAC
R  ACTACTAGGTCTTCTCTTAGCCTCAGCAGGAAAATCAGCCCAATTTGGACTTCATCCGTGACTACCATTAATCACCGAAGGCCCTACCCCAGTATCAGCACTCCTTCATTCCAGTACAATAGTTATAGCTGGAGTATTTACCCTCATCCGCTTTTATCCATTAATAGAAAACAACCTCAC
P  ATCCACGTCAGCCATAACCACGTCAATGGTCGCCAAATAAATGATCAAGTCCGAAACCAGACTACCATTAATCACCGAAGGCATTTGCTGCAAGTGGGTGAAGGTTGGCGTACGAATACGTGTACGATAGCTCATGGTTGACTTGTCAGATGTGAGGTAATACGTGCTCGCACCTTTTAC
Pos                                                         <-717313-------717291->
R translated                                           ...V  T  t  I  n  h  r  R  P...
```

$\Delta G^0_{min}$ = -13.04 kcal/mol

```
S75.                                   |-- net 90 bp loss ------------------------------------------------------------------------|
A  ACTACTAGGTCTTCTCTTAGCCTCAGCAGGAAAATCAGCCCAATTTGGACTTCATCCGTGACTTCCATCAGCTAGTGAAGGCCCTACCCCAGTATCAGCACTCCTTCATTCCAGTACAATAGTTATAGCTGGAGTATTTACCCTCATCCGCTTTTATCCATTAATAGAAAACAACCTCAC
R  ACTACTAGGTCTTCTCTTAGCCTCAGCAGGAAAA------------------------------------------------------------------------------------------ATAGCTGGAGTATTTACCCTCATCCGCTTTTATCCATTAATAGAAAACAACCTCAC
P  AGCATGCAAACTAAGCCATGCAGAAGCAGGAAAA------------------------------------------------------------------------------------------ATAGCCGAAGCAAACAATAGCCCCTTGGGATTAAGCAGCGTTGCACAAAACAGTTC
Pos                        <-760735--------------------------------------------------------------------------------------------760716->
R translated            ...S  R  K  n                                                                                      S  W  S...
```

$\Delta G^0_{min}$ = -13.32 kcal/mol + -6.69 kcal/mol  (average: -10.01 kcal/mol)

```
MK73.                                                                   |--------| no net loss/gain
A  ACTACTAGGTCTTCTCTTAGCCTCAGCAGGAAAATCAGCCCAATTTGGACTTCATCCGTGACTTCCATCAGCTAGTGAAGGCCCTACCCCAGTATCAGCACTCCTTCATTCCAGTACAATAGTTATAGCTGGAGTATTTACCCTCATCCGCTTTTATCCATTAATAGAAAACAACCTCAC
R  ACTACTAGGTCTTCTCTTAGCCTCAGCAGGAAAATCAGCCCAATTTGGACTTCATCCGTGACTTCCATCAGCCAGCAAACGTCCTACCCCAGTATCAGCACTCCTTCATTCCAGTACAATAGTTATAGCTGGAGTATTTACCCTCATCCGCTTTTATCCATTAATAGAAAACAACCTCAC
P  ACACAATACTTTCTGATGATCAGAATAAGGAACGCTGGTCGAACCGATAAAACCAGCAAGATCTGAAACAGCCAGCAAACGTCCCCGAATATTGGCCAGACCTTTAAACCATTGCTGAGTATTCGGGACAGGAATACAAACAGGCGGATATATCACTTCTGCTACCTCACCTAATGGTAC
Pos                                                          776226<-------------->776211
R translated                                                   ...I  S  q  q  t  s  Y...
```

$\Delta G^0_{min}$ = -10.34 kcal/mol

**MK49.**

```
                                                                                                                                            net 1 bp inserted |---------|
A  ACTACTAGGTCTTCTCTTAGCCTCAGCAGGAAAATCAGCCCAATTTGGACTTCATCCGTGACTTCCATCAGCTAGTGAAGGCCCTACCCCAGTATCAGCACTCCTTCATTCCAGTACAATAGTTATAGCTGGAGTATTTACCCTCATCCGCTTT-TATCCATTAATAGAAAACAACCTCAC
   ||||||||||||||||||||||||||||||||||||||||||||||||||||||||||||||||||||||||||||||||||||||||||||||||||||||||||||||||||||||||||||||||||||||||||||||||||| |    ||| |||||||||||||||||||||||
R  ACTACTAGGTCTTCTCTTAGCCTCAGCAGGAAAATCAGCCCAATTTGGACTTCATCCGTGACTTCCATCAGCTAGTGAAGGCCCTACCCCAGTATCAGCACTCCTTCATTCCAGTACAATAGTTATAGCTGGAGTATTTACCCTCATCAGGAGAATATTCATTAATAGAAAACAACCTCAC
   |   ||     |  |  |          |       |  |     | || |  | |         |      |      |       || ||   | |  |  || |||  |   ||       |  |    |||||||||||||||||||||   || |  |   |    |
P  ATGGCTTTACGTCTTGCTGGTGAGTTGCTTGATGCAGCTGAAGGTAAAGGTGCAGCAATCAAAAAACGTGAAGATGTGCATCGTATGGCTGAAGCCAACAAAGCCTTCTCTCACTACCGTTTCTAAGCGGATAAAACAGTCCCTCATCAGGAGAATATTCATGCGTACAGCAGCTCGATTC
Pos                                                                                                                                                   <-865908------865929->
R translated, new start:                                                                                                          i  v  i  a  g  v  f  t  p  i  r  r  i  f  I  N  R  K  Q  P...
                                                                                                                                  |     |                             |
                                                                                                                                  potential ATA start codons?----------
```

$\Delta G^0_{min}$ = -8.18 kcal/mol

**MK54.**

```
                                                                        |-- net 15 bp loss ----------------------------------|
A  GCTAGTGAAGGCCCTACCCCAGTATCAGCACTCCTTCATTCCAGTACAATAGTTATAGCTGGAGTATTTACCCTCATCCGCTTTTATCCATTAATAGAAAACAACCTCACTATTCAAACTTCAACACTTCTGGTTCTGGCTCTAATGTCACTAATGAAAAAATGCGTTTCTGGAGTCCAG
   |||||||||||||||||||||||||||||||||||||||||||||||||||||||||||||||||||||||| |     |  |  |    |   ||| ||  | ||| | |  ||||||  |      ||||||||||||||||||||||||||||||||||||||||||||||||||||||||
R  GCTAGTGAAGGCCCTACCCCAGTATCAGCACTCCTTCATTCCAGTACAATAGTTATAGCTGGAGTATTTACCTTAAAAGGAATAGAA--A--AAT-GAGCA-AACTT-A--ATTCAA--TG----ACTTCTGGTTCTGGCTCTAATGTCACTAATGAAAAAATGCGTTTCTGGAGTCCAG
     |    |      ||       ||  ||      | | | |        |  || |    ||||||||||||||||||||||||||  |  |  ||| ||||| ||||| |  ||||||  ||    ||||||| | | | |     |       |                 |  | |
P  TATTTGCACTGTAATAAATTGAGATTTGCGCCGTCTAACTTTATGTACCTCTGTACAAACCGAGTATTTACCTTAAAAGGAATAGAA--A--AAT-GAGCA-AACTT-A--ATTCAA--TG----ACTTCAGTTGCAGCATTATTGGCATTATCAGTCGCTGGTGTAACTCAGGTGGCTG
Pos                                                             <-916228-----------------------------------------------------916283->
R translated                                                                                  m   s  k   l   n    s  m        T  S  G  S  G  S  N...
```

$\Delta G^0_{min}$ = -12.46 kcal/mol + -7.12 kcal/mol (average: -9.79 kcal/mol)

**B18.**

```
                                                                          |--| no net loss/gain
A  ACTACTAGGTCTTCTCTTAGCCTCAGCAGGAAAATCAGCCCAATTTGGACTTCATCCGTGACTTCCATCAGCTAGTGAAGGCCCTACCCCAGTATCAGCACTCCTTCATTCCAGTACAATAGTTATAGCTGGAGTATTTACCCTCATCCGCTTTTATCCATTAATAGAAAACAACCTCAC
   |||||||||||||||||||||||||||||||||||||||||||||||||||||||||||||||||||||||||| || |||||||||||||||||||||||||||||||||||||||||||||||||||||||||||||||||||||||||||||||||||||||||||||||||||
R  ACTACTAGGTCTTCTCTTAGCCTCAGCAGGAAAATCAGCCCAATTTGGACTTCATCCGTGACTTCCATCAGATATTGCAGGCCCTACCCCAGTATCAGCACTCCTTCATTCCAGTACAATAGTTATAGCTGGAGTATTTACCCTCATCCGCTTTTATCCATTAATAGAAAACAACCTCAC
            | || |  | ||  |    | | |  |         |  | |     | | ||   | ||||||||||||     |    |  |        |      ||    |   |  | || | | ||      |     |  ||                     |   |
P  TGCGGGTTCTATTTTTCTGGCACCTCTGGCAGACAAAATTGGTCGCCGTTTACTGATTTAATTTGTTTAGCTATTGCAGGCATCAGCATGCTCGCCTGTGGTTTGGTGCATAGCCACAGTATGTTGGCAGCATTACGCTTTGTGACAGGTATTGGGGTGGGGGTATTTTAGCCAGTAG
Pos                                                                  968658 <-----------> 968670
R translated                                                             ...S  y  c  R  P...
```

$\Delta G^0_{min}$ = -12.91 kcal/mol

**MK150.**

```
                                                                         |--| no net gain/loss
A  ACTACTAGGTCTTCTCTTAGCCTCAGCAGGAAAATCAGCCCAATTTGGACTTCATCCGTGACTTCCATCAGCTAGTGAAGGCCCTACCCCAGTATCAGCACTCCTTCATTCCAGTACAATAGTTATAGCTGGAGTATTTACCCTCATCCGCTTTTATCCATTAATAGAAAACAACCTCAC
   ||||||||||||||||||||||||||||||||||||||||||||||||||||||||||||||||||||||| |  ||||||||||||||||||||||||||||||||||||||||||||||||||||||||||||||||||||||||||||||||||||||||||||||||||||||||
R  ACTACTAGGTCTTCTCTTAGCCTCAGCAGGAAAATCAGCCCAATTTGGACTTCATCCGTGACTTCCATCAGCGAAAGAAGGCCCTACCCCAGTATCAGCACTCCTTCATTCCAGTACAATAGTTATAGCTGGAGTATTTACCCTCATCCGCTTTTATCCATTAATAGAAAACAACCTCAC
   |  |  |     |         | |   |   |||   | |||       |       || || |||||||||||||        |  |     |   |       | ||  |    |  ||     |   ||  |    ||      |
P  AGCAGAACAAAATGGTGCGCATTTACATCTACTTGCAGGTGCAATTGCTGGGATTGATGGAATTGCAGCAGCGAAAGAAGGCGGCTTGGAAAAAGTGACCTACAAAGGCTGCAAAAGTCCAAATAGTTGGCGTGGCAGTTATGCCGAGCAGTTAATTGATTTGGATCAGGTACATACAGT
Pos                                                        982222 <------------------> 982241
R translated                                                 ...T  S  I  S  e  r  R  P...
```

$\Delta G^0_{min}$ = -16.15 kcal/mol

**MK48.**

```
                 |-- net 84 bp loss ------------------------------------------------------------------------------------|
A  TAGAAAGATGAAACACTACACTCCCACTACTAGGTCTTCTCTTAGCCTCAGCAGGAAAATCAGCCCAATTTGGACTTCATCCGTGACTTCCATCAGCTAGTGAAGGCCCTACCCCAGTATCAGCACTCCTTCATTCCAGTACAATAGTTATAGCTGGAGTATTTACCCTCATCCGCTTTTAT
   ||||||||||||||||   |                                                                                |     ||| ||||||||||||||||||||||||||||||||||||||||||||||||||||||||||||||||||||||
R  TAGAAAGATGAAACACACCTGCTT----------------------------------------------------------------------------CACGTGCAGCATCAGCACTCCTTCATTCCAGTACAATAGTTATAGCTGGAGTATTTACCCTCATCCGCTTTTAT
   |  |  |||||||||||||||||                                                                               ||||||||||||||||             | |                | | ||    | |  |  |    |  |
P  TCAAGTGATGAAACACACCTGCTT----------------------------------------------------------------------------CACGTGCAGCATCAGCCTGGAAAGTACGGATTTTTCATCAGCATCAGCAGCTAATTCAGTGTTATCGTACTCA
Pos     <-1081237-----------------------------------------------------------------------------------------1081204->
R tl   ...M  K  H  t  c  f                                                                               t  c  a  S  A...
```

$\Delta G^0_{min}$ = -11.77 kcal/mol + -9.93 kcal/mol (average: -10.85 kcal/mol)

**MK117.**

```
                                                   |-- net 12 bp loss ---------------|
A  ACTACTAGGTCTTCTCTTAGCCTCAGCAGGAAAATCAGCCCAATTTGGACTTCATCCGTGACTTCCATCAGCTAGTGAAGGCCCTACCCCAGTATCAGCACTCCTTCATTCCAGTACAATAGTTATAGCTGGAGTATTTACCCTCATCCGCTTTTATCCATTAATAGAAAACAACCTCAC
   |||||||||||||||||||||||||||||||||||||||||||||     |||    | ||| || ||| ||| | ||| |||||||||||||||||||||||||||||||||||||||||||||||||||||||||||||||||||||||||||||||||||||||||||||||||||||
R  ACTACTAGGTCTTCTCTTAGCCTCAGCAGGAAAATCAGCCAATTT-----TCA--GAT-ACTGCC-TCA-CTA-T-AAGTCCCTACCCCAGTATCAGCACTCCTTCATTCCAGTACAATAGTTATAGCTGGAGTATTTACCCTCATCCGCTTTTATCCATTAATAGAAAACAACCTCAC
      |    |    |       |  | |     |   | |||||||     |||  ||| |||||| ||| ||| | ||||||||| |          |          || |     | ||     ||      | |||  |    |    |                 |  |  | |
P  TTCACGGCGGTAACCGGGGTTCGAATCCCCGTAGGGACGCCAATTT-----TCA--GAT-ACTGCC-TCA-CTA-T-AAGTCCCTATCGTCTAGAGGCCTAGGACATCGCCCTTTCACGGCGGTAACCGGGGTTCGAATCCCCGTAGGGACGCCAAATTCGATCACTATGATCATCAGTA
Pos                                          <-1136013-----------------------------1135979->
R translated                             ...P  I  f      r   y   c  l   t   i   s  P  Y...
```

$\Delta G^0_{min}$ = -6.55 kcal/mol + -16.18 kcal/mol (average: -11.37 kcal/mol)

**MK7.**

```
                                                                                         |-----------| net 3 bp inserted
A  ACTACTAGGTCTTCTCTTAGCCTCAGCAGGAAAATCAGCCCAATTTGGACTTCATCCGTGACTTCCATCAGCTAGT-GAAG-GCC-CTACCCCAGTATCAGCACTCCTTCATTCCAGTACAATAGTTATAGCTGGAGTATTTACCCTCATCCGCTTTTATCCATTAATAGAAAACAACCT
   ||||||||||||||||||||||||||||||||||||||||||||||||||||||||||||||||||||||||||| |  | | |||  ||||||||||||||||||||||||||||||||||||||||||||||||||||||||||||||||||||||||||||||||||||||||||||
R  ACTACTAGGTCTTCTCTTAGCCTCAGCAGGAAAATCAGCCCAATTTGGACTTCATCCGTGACTTCCATCAGCTACTTCATGCGCCATTACCCCAGTATCAGCACTCCTTCATTCCAGTACAATAGTTATAGCTGGAGTATTTACCCTCATCCGCTTTTATCCATTAATAGAAAACAACCT
          |                                                   |  |   |||  ||||||||||||||||||||||||||    |
P  GGCTAAAGCGAGTAAACCTTGTCCCTTAAAGGCTTCTTGTGAACGACTGTAATGACGTTGTGTAACATGAGCTACTTCATGCGCCATTACCCCAGCCACCTCATCAATATTTCGTGCATTTGCGATTAAACCTGTATTTAAGGCAAATAACCCACCTGGAACAGCAAATGCATTAATTTG
Pos                                                                  <-1326406------------1326377->
R translated                                                    ...S  I  S  y  f  m  r  h  Y  P  S...
```

$\Delta G^0_{min}$ = -20.34 kcal/mol

**MK16.**

```
                                                                                         |-------| no net gain/loss
A  ACTACTAGGTCTTCTCTTAGCCTCAGCAGGAAAATCAGCCCAATTTGGACTTCATCCGTGACTTCCATCAGCTAGTGAAGGCCCTACCCCAGTATCAGCACTCCTTCATTCCAGTACAATAGTTATAGCTGGAGTATTTACCCTCATCCGCTTTTATCCATTAATAGAAAACAACCTCAC
   ||||||||||||||||||||||||||||||||||||||||||||||||||||||||||||||||||||||||||     |||  ||||||||||||||||||||||||||||||||||||||||||||||||||||||||||||||||||||||||||||||||||||||||||||||||
R  ACTACTAGGTCTTCTCTTAGCCTCAGCAGGAAAATCAGCCCAATTTGGACTTCATCCGTGACTTCCATCAGCTACATGAGGTTCTACCCCAGTATCAGCACTCCTTCATTCCAGTACAATAGTTATAGCTGGAGTATTTACCCTCATCCGCTTTTATCCATTAATAGAAAACAACCTCAC
     |       | |  ||        |          |||     |       |||    |      ||   |||||||||||||||||||||   ||  ||    |         |     |          |         |         |          ||           ||||
P  TGTGAAGACACATGGCTAGCTTAGATTTACCTGATCATTTAATCCACACCTTATCACAAGTTTTAAATCAGCTACATGAGGTTCTACCAAAGATTCGTCAAGAACCTGATTTCTCAGCACCTGCATTTAAATGGCAAGATAAACAACTTAAGCCCATTTTTGACCCCAAAAAATTTGATT
Pos                                                                    <-1331952----1331931->
R translated                                                       ...I  S  y  m  R  m  Y  P...
```

$\Delta G^0_{min}$ = -14.25 kcal/mol

**K86.**

```
                                                                                      |------| no net loss/gain
A  ACTACTAGGTCTTCTCTTAGCCTCAGCAGGAAAATCAGCCCAATTTGGACTTCATCCGTGACTTCCATCAGCTAGTGAAGGCCCTACCCCAGTATCAGCACTCCTTCATTCCAGTACAATAGTTATAGCTGGAGTATTTACCCTCATCCGCTTTTATCCATTAATAGAAAACAACCTCAC
   |||||||||||||||||||||||||||||||||||||||||||||||||||||||||||||||||||||||  ||| | ||||||||||||||||||||||||||||||||||||||||||||||||||||||||||||||||||||||||||||||||||||||||||||||||||||
R  ACTACTAGGTCTTCTCTTAGCCTCAGCAGGAAAATCAGCCCAATTTGGACTTCATCCGTGACTTCCATCAGTAAGTTATGGCCCTACCCCAGTATCAGCACTCCTTCATTCCAGTACAATAGTTATAGCTGGAGTATTTACCCTCATCCGCTTTTATCCATTAATAGAAAACAACCTCAC
    | ||    | | |    |      |       |||||  | |   ||   |   ||  ||     ||||||||||||||||||       | |          || |        |    | | |    |      ||  |           | |  ||     |      || ||
P  CCAACGTTATTTGCAAAAACATTATGATCACCAATCACACAATCATGTGCAATATGAGTATTGACCATCAGTAAGTTATGGCTACCGATTTTGGTTAAACTGTTATCTTGTATCGTCCCACGATGCAACTGCAATGTTCACGAATAAGATTATAGTCTCCAATCTCTAACCAGGTCTCT
Pos                                                                  <----------------> 1378419-1378402
R translated                                                    ...S  I  S  k  l  w  P...
```

$\Delta G^0_{min}$ = -12.72 kcal/mol

**MK35.**

```
                                                                               |-- net 63 bp loss ---------------------------------------------------------------|
A  ACTACTAGGTCTTCTCTTAGCCTCAGCAGGAAAATCAGCCCAATTTGGACTTCATCCGTGACTTCCATCAGCTAGTGAAGGCCCTACCCCAGTATCAGCACTCCTTCATTCCAGTACAATAGTTATAGCTGGAGTATTTACCCTCATCCGCTTTTATCCATTAATAGAAAACAACCTCAC
   |||||||||||||||||||||||||||||||||||||||||||||||||||||||||||||||||||    |||| |                                                               |   ||| ||| ||||||||||||||||||||||||||||||
R  ACTACTAGGTCTTCTCTTAGCCTCAGCAGGAAAATCAGCCCAATTTGGACTTCATCCGTGACTTCCCCAGCGA---------------------------------------------------------------TCTGCCTTATCTGCTTTTATCCATTAATAGAAAACAACCTCAC
        |  |     |                |||    |                   | | ||||||||||||                                                               ||||||||||||| |  ||    |   |||      |  |
P  TTGTTTTTGCAGCTTTGGCCATAATCGGACCAAAATTTCTGGCGGCTTTACGATAAGTTAAATTCCCCCAGCGA---------------------------------------------------------------TCTGCCTTATCTGCATAAATGAATGCATAATCGGCTTCAAGTG
Pos                                                               <-1701316----------------------------------------------------------------------1701291->
R translated                                                   ...T  S  p  s  d                                                      l  P  y  l  L...
```

$\Delta G^0_{min}$ = -8.12 kcal/mol + -7.50 kcal/mol  (average: -7.81 kcal/mol)

**A19.**

```
                                                                                      |--------------------| net 6 bp loss
A  ACTACTAGGTCTTCTCTTAGCCTCAGCAGGAAAATCAGCCCAATTTGGACTTCATCCGTGACTTCCATCAGCTAGTGAAGGCCCTACCCCAGTATCAGCACTCCTTCATTCCAGTACAATAGTTATAGCTGGAGTATTTACCCTCATCCGCTTTTATCCATTAATAGAAAACAACCTCAC
   |||||||||||||||||||||||||||||||||||||||||||||||||||||||||||||||||||||| ||   |   | | ||       | ||||||||||||||||||||||||||||||||||||||||||||||||||||||||||||||||||||||||||||||||||
R  ACTACTAGGTCTTCTCTTAGCCTCAGCAGGAAAATCAGCCCAATTTGGACTTCATCCGTGACTTCCATCAGTTATCGTCGTCACT------GGATCAGCACTCCTTCATTCCAGTACAATAGTTATAGCTGGAGTATTTACCCTCATCCGCTTTTATCCATTAATAGAAAACAACCTCAC
      |  | |    |    | |     |  ||      ||        | |  |   |           ||||||||||||||||||||      ||||||||||| |         |   |    |  |  || |       |     |    |     ||     |    |  |||     ||       |   |
P  TGAAAGATGCGATGCATCAATGACTTCAAACTGATTGCATTGAATATGCTGTTGCATAAATTCACCATCAGTTATCGTCGTCACT------GGATCAGCACTACCCGCGATAATCAATGTCGGAATTGAGATTGATGCCAATTTTTCACGTACATCCGCTTTAGCCAAAGCACGGCAAGC
Pos                                                                  <-1713833--------------------1713802->
R translated                                                    ...S  I  S  y  r  r  h  w        I  S  T  P...
```

$\Delta G^0_{min}$ = -9.20 kcal/mol + -11.56 kcal/mol  (average: -10.38 kcal/mol)

**R17.**

```
                                                                                  |-- net 3 bp loss ---------|
A  ACTACTAGGTCTTCTCTTAGCCTCAGCAGGAAAATCAGCCCAATTTGGACTTCATCCGTGACTTCCATCAGC-TAGTGAAGGCCCTACCCCAGTATCAGCACTCCTTCATTCCAGTACAATAGTTATAGCTGGAGTATTTACCCTCATCCGCTTTTATCCATTAATAGAAAACAACCTCAC
   ||||||||||||||||||||||||||||||||||||||||||||||||||||||||||||||||||||||| || |||| ||| || |       || |||||||||||||||||||||||||||||||||||||||||||||||||||||||||||||||||||||||||||||||||||||
R  ACTACTAGGTCTTCTCTTAGCCTCAGCAGGAAAATCAGCCCAATTTGGACTTCATCCGTGACTTCCACCTGCGTAGTAAAGACCATTT----GTTTCAGCACTCCTTCATTCCAGTACAATAGTTATAGCTGGAGTATTTACCCTCATCCGCTTTTATCCATTAATAGAAAACAACCTCAC
         |    |    |     | |           |     ||   ||||||     || |   ||||||||||||||||||||||||     |||||||||||  |  |   |      |  | |  |       ||||| |      |    |   |  |  |  | | |            | |
P  GTATAAATTATTAACATCCAGATAAAATAATCCTAAACGTAAAGCCGGACTTACATTGTAATAGCCACCTGCGTAGTAAAGACCATTT----GTTTCAGCACTTTGAAACGCCCATTCACCGCTGAAATCCTGATATTTTCATAATTGGTCGAGTCGCGCTGATTCACCTTGTCTATAAA
Pos                                                                  <-1730059---------------------1730093->
R translated                                                    ...S  t  c  v  v  k  t  i  c     f  S  T...
```

$\Delta G^0_{min}$ = -10.32 kcal/mol + -10.58 kcal/mol  (average: -10.45 kcal/mol)

```
A60.                                                                                                |-| no net loss/gain
A  ACTACTAGGTCTTCTCTTAGCCTCAGCAGGAAAATCAGCCCAATTTGGACTTCATCCGTGACTTCCATCAGCTAGTGAAGGCCCTACCCCAGTATCAGCACTCCTTCATTCCAGTACAATAGTTATAGCTGGAGTATTTACCCTCATCCGCTTTTATCCATTAATAGAAAACAACCTCAC
   |||||||||||||||||||||||||||||||||||||||||||||||||||||||||||||||||||||||||||||||||||||| | ||||||||||||||||||||||||||||||||||||||||||||||||||||||||||||||||||||||||||||||||||||||||||||||||||||||
R  ACTACTAGGTCTTCTCTTAGCCTCAGCAGGAAAATCAGCCCAATTTGGACTTCATCCGTGACTTCCATCAGCTATTTAAGGCCCTACCCCAGTATCAGCACTCCTTCATTCCAGTACAATAGTTATAGCTGGAGTATTTACCCTCATCCGCTTTTATCCATTAATAGAAAACAACCTCAC
      |   |    |  | |  |        |   |      ||         |         ||   |    | | || | |  |||||||||||    |      |       | |  |     |         |  |  |     |        |        |             |             |  ||   ||      | |
P  TATGGTTTTTTGTTTGCTTTTAAAATTATATTTATTTTACTTAGGCTATTTTATTTTTTAATTTGCTTAGGCTATTTAAGGCTACAAAGTAAATGAATCGATTTGCCCAAAAGTCAAAGTAAACTTGCAAAAAAGACGCAGATAATCAAACAACGGATGTCTCCTAACCAAATTAATGAA
Pos                                                                                     <----------> 1747667-1747656
R translated                                                                        ...S  y  l  R  P...
```

$\Delta G^{0}_{min}$ = -9.87 kcal/mol

```
MK134.                                                                                         |-----| no net loss/gain
A  ACTACTAGGTCTTCTCTTAGCCTCAGCAGGAAAATCAGCCCAATTTGGACTTCATCCGTGACTTCCATCAGCTAGTGAAGGCCCTACCCCAGTATCAGCACTCCTTCATTCCAGTACAATAGTTATAGCTGGAGTATTTACCCTCATCCGCTTTTATCCATTAATAGAAAACAACCTCAC
   |||||||||||||||||||||||||||||||||||||||||||||||||||||||||||||||||||||||||||||||||| | ||  ||||||||||||||||||||||||||||||||||||||||||||||||||||||||||||||||||||||||||||||||||||||||||||||||||||||
R  ACTACTAGGTCTTCTCTTAGCCTCAGCAGGAAAATCAGCCCAATTTGGACTTCATCCGTGACTTCCATCAACCAGCAAAGGCCCTACCCCAGTATCAGCACTCCTTCATTCCAGTACAATAGTTATAGCTGGAGTATTTACCCTCATCCGCTTTTATCCATTAATAGAAAACAACCTCAC
       |    | ||  |    |    ||                       |  || |  |   |      ||||||||||||||    |     |    |   ||  |      |            |       ||      ||||||          |     ||  ||   || |   ||| |  |
P  TAATTTTGATTTTAGCCCCGTTGCAAGTTTTTGTGGGCGATGCACATGGTTTAAACACTTTAGAGCATCAACCAGCAAAACTCGCGGCTATGGAAGGTCATTGGGAAACCAACCATGATCATGCTATGCCTCTTTATTTATTCGGTATTCCAGACATGCAGGAAGAACGAACCAAATATG
Pos                                                                          1757779<------------>1757767
R translated                                                               ...S  I  n  q  q  R...
```

$\Delta G^{0}_{min}$ = -6.61 kcal/mol

```
MK46.                                                                                            |---| no net gain/loss
A  ACTACTAGGTCTTCTCTTAGCCTCAGCAGGAAAATCAGCCCAATTTGGACTTCATCCGTGACTTCCATCAGCTAGTGAAGGCCCTACCCCAGTATCAGCACTCCTTCATTCCAGTACAATAGTTATAGCTGGAGTATTTACCCTCATCCGCTTTTATCCATTAATAGAAAACAACCTCAC
   |||||||||||||||||||||||||||||||||||||||||||||||||||||||||||||||||||||||||||||||||||||| | ||||||||||||||||||||||||||||||||||||||||||||||||||||||||||||||||||||||||||||||||||||||||||||||||||||
R  ACTACTAGGTCTTCTCTTAGCCTCAGCAGGAAAATCAGCCCAATTTGGACTTCATCCGTGACTTCCATCAGCCACTTAAGGCCCTACCCCAGTATCAGCACTCCTTCATTCCAGTACAATAGTTATAGCTGGAGTATTTACCCTCATCCGCTTTTATCCATTAATAGAAAACAACCTCAC
      |   |        ||   ||    |  | ||  | |     |   |     |       ||    |||||||||||||||   |       | |    |            |    |                     |   |         |        | |  |    |
P  TTGAAATGAGTAATGGTTCATCTGTCGAACATAAAGAACTTGCCTCTAAAGCATTTGAAATTTTGACACAGCCACTTAAGGCCTAATACTTAAAACTAGCCCAAAGAAGAGTCGAGAGCTTTGGGCTAAATTTTTGCAATAATATAGAGGTGAAATATGAAGTTTGCCACAATAAAAAAT
Pos                                                                             1816041<------------->1816055
R translated                                                                     ...I  S  h  l  S  P...
```

$\Delta G^{0}_{min}$ = -14.01 kcal/mol

```
MK51.                                                                                   |--------------------------| no net gain/loss
A  ACTACTAGGTCTTCTCTTAGCCTCAGCAGGAAAATCAGCCCAATTTGGACTTCATCCGTGACTTCCATCAGCTAGTGAAGGCCCTACCCCAGTATCAGCACTCCTTCATTCCAGTACAATAGTTATAGCTGGAGTATTTACCCTCATCCGCTTTTATCCATTAATAGAAAACAACCTCAC
   |||||||||||||||||||||||||||||||||||||||||||||||||||||||||||||||||||||||||||||||||| |||          ||  ||   |  ||||     ||||||||||||||||||||||||||||||||||||||||||||||||||||||||||||||||||||||||||||
R  ACTACTAGGTCTTCTCTTAGCCTCAGCAGGAAAATCAGCCCAATTTGGACTTCATCCGTGACTTCCACCAGAACCGCCAGCACCACCACCAGCGCCAGCACTCCTTCATTCCAGTACAATAGTTATAGCTGGAGTATTTACCCTCATCCGCTTTTATCCATTAATAGAAAACAACCTCAC
   || ||   |   | ||     | |  |||     |     || | | |||    ||   |         |    |||||||||||||||||||||||||||||||||||||   |       |      |   |     | |    || |    |   |    |        |      |  || ||       |
P  ACCACCTGAGCCTCCAGCACCACCAGATCCACCTGCACCGCCATTACCACCGCCTGACCCAGAACCACCAGAACCGCCTGCACCACCACCAGCGCCAGCACCGCCATTGGCACCCGAACCACCATCACCGCCAGAACCGCCATTAGCACCGACACAGTTAATAGTATTGCCTGAAACTGT
Pos                                                                               <-1854121-------------------1854157->
R translated                                                               ...S  t  r  t  a  c  t  t  t  S  a  S  T...
```

$\Delta G^{0}_{min}$ = -15.14 kcal/mol

```
A6.                                                                                              |----| no net gain/loss
A  ACTACTAGGTCTTCTCTTAGCCTCAGCAGGAAAATCAGCCCAATTTGGACTTCATCCGTGACTTCCATCAGCTAGTGAAGGCCCTACCCCAGTATCAGCACTCCTTCATTCCAGTACAATAGTTATAGCTGGAGTATTTACCCTCATCCGCTTTTATCCATTAATAGAAAACAACCTCAC
   |||||||||||||||||||||||||||||||||||||||||||||||||||||||||||||||||||||||||||||||||||||| |  ||||||||||||||||||||||||||||||||||||||||||||||||||||||||||||||||||||||||||||||||||||||||||||||||||||
R  ACTACTAGGTCTTCTCTTAGCCTCAGCAGGAAAATCAGCCCAATTTGGACTTCATCCGTGACTTCCATCAGCCATAGCAGGCCCTACCCCAGTATCAGCACTCCTTCATTCCAGTACAATAGTTATAGCTGGAGTATTTACCCTCATCCGCTTTTATCCATTAATAGAAAACAACCTCAC
   |     ||  |   |     ||  ||      |    ||| |   |   | ||          |      || |||||||||||||||||       ||  | |       ||| |   |       |   | || ||| ||| ||         |  | |  |||      |   |      |        ||
P  ATCTGTATTTGGACATGAAGAATCTTTGACAGGGTCACCAAAGACTAGATGCATCGATGGGTCATCACCAGCCATAGCAGGCCCTTTGTATCTACAACCTTGATAACATGCTGTTTGTGATGAGATTGCTTGAGAATAAGATACAGTTTGTTCATATGTCCAAGAATGCCAGCCAGGGAC
Pos                                                                                   <-1854425--1854444->
R translated                                                               ...S  I  S  h  s  R  P  Y...
```

$\Delta G^{0}_{min}$ = -16.79 kcal/mol

```
O140.               |-- net 129 bp loss -------------------------------------------------------------------------------------------------------------------|
A  GGTCTTCTCT-TAGCCTCAGCAGGAAAATCAGCCCAATTTGGACTTCATCCGTGACTTCCATCAGCTAGTGAAGGCCCTACCCCAGTATCAGCACTCCTTCATTCCAGTACAATAGTTATAGCTGGAGTATTTACCCTCATCCGCTTTTATCCATTAATAGAAAACAACCTCACTATTCAA
   |||||||||| ||||| || ||||  |                                                                                                                           |||||||||||||||||||||
R  GGTCTTCTCT-TAGCCCCAACAGGTTAGCC-----------------------------------------------------------------------------------------------------------------------------AGAAAACAACCTCACTATTCAA
   |     |||| ||||||||||||||||||                                                                                                                             ||||||||||| |   | |  |
P  GTAGGGCTCTATAGCCCCAACAGGTTAGCC-----------------------------------------------------------------------------------------------------------------------------AGAAAACAACCACCTCAGTTAT
Pos     <-1855272-----------------------------------------------------------------------------------------------------------------------------------------------1855306->
R tl ...S  L   S  p  n  R  l  a                                                                                                                              R  K  Q  P...
```

$\Delta G^{0}_{min}$ = -17.87 kcal/mol + -14.05 kcal/mol  (average: -15.96 kcal/mol)

**MK114.**

```
                                                                                                                                                  |-----------| net 1 bp inserted
A  TTCATCCGTGACTTCCATCAGCTAGTGAAGGCCCTACCCCAGTATCAGCACTCCTTCATTCCAGTACAATAGTTATAGCTGGAGTATTTACCCTCATCCGCTTTTATCCATTAATAGAAAACAACCTCACTATTCA-AACTTCAACACTTCTGGTTCTGGCTCTAATGTCACTAATGAAA
R  TTCATCCGTGACTTCCATCAGCTAGTGAAGGCCCTACCCCAGTATCAGCACTCCTTCATTCCAGTACAATAGTTATAGCTGGAGTATTTACCCTCATCCGCTTTTATCCATTAATAGAAAACAACCTGACGATGAATAATTTCAACACTTCTGGTTCTGGCTCTAATGTCACTAATGAAA
P  AGCAATTTAGTTGAAAATTTGATATAAGGTTTTGATAATTAAAATTTTATTAAATGTTATATTTTTGTGGATTAACCGTATATTATTCATTAATATACAAGCTATTCTGTATGCAAAATAGAAACCTGACGATGAATAATTTCTCATTTACTCAATCAGAGGACGCTCGTCATGAAGCGT
Pos                                                                                                                                         <-1859219---1859199->
R translated                                                                                                                                          m  n  n  F...
```

$\Delta G^0_{min}$ = -9.72 kcal/mol

**MK106.**

```
                                                       |-- net 195 bp loss -------------------------------------------------------------------------|
A  CACGTTCTGCACGTTATCGTTATCAGTAAGTATTAATATTTTAGCAAGATCAGAGCTTAGTAACTGCTCTACGGGGCTGATTTTCAGCCCTTTTTTGAGAATAGAAAGATGAAACACTACACTCCCACTACTAG(120bp)CTGGAGTATTTACCCTCATCCGCTTTTATCCATTAATAGA
R  CACGTTCTGCACGTTATCGTTATCAGTAAGTATTAATATTTTAGCAAGATCAGGACAGT--------------------------------------------------------------------------------TATGCGTATTTACCCTCATCCGCTTTTATCCATTAATAGA
P  AACAAGTCTATAATGCGCTTGTTTCTGGCTCTTTGGTTATTGCAAAAGATCAGGACAGT--------------------------------------------------------------------------------TATGCGTATTTACCTAAACCACTTGCAGATAAAATCAATC
Pos                                                <-1863255----------------------------------------------------------------------------------------1863282->
R translated                                                                                               |                                          m  r  I  Y  P...
```

$\Delta G^0_{min}$ = -8.18 kcal/mol + -10.30 kcal/mol  (average: -9.24 kcal/mol)

**MK97.**

```
                                                                                 |-- net 6 bp loss -------|
A  ACTACTAGGTCTTCTCTTAGCCTCAGCAGGAAAATCAGCCCAATTTGGACTTCATCCGTGACTTCCATCAGCTAGTGAAGGCCCTACCCCAGTATCAGCACTCCTTCATTCCAGTACAATAGTTATAGCTGGAGTATTTACCCTCATCCGCTTTTATCCATTAATAGAAAACAACCTCAC
R  ACTACTAGGTCTTCTCTTAGCCTCAGCAGGAAAATCAGCCCAATTTGGACTTCATCCGTGACTTCCATC--C-A-TG-TCGCAATA-CGCACTAGCAGCACTCCTTCATTCCAGTACAATAGTTATAGCTGGAGTATTTACCCTCATCCGCTTTTATCCATTAATAGAAAACAACCTCAC
P  ATTAATAATGCAATCGATTGTAAACTCTTTGAAATTACGAAAATTGCCAACCAGATGCACGCTTCCATC--C-A-TG-TCGCAATA-CGCACTAGCAGCATCCTTTTGCCAGTCTTATCTGCCACCCAGCCCCATATAAATGGCGCAAAAAATCGTGTAATGATGGCAATCGAAGAAAGT
Pos                                                            <-1865765--------------------1865802->
R translated                                                ...T  S  I   h   v   a  I   r  t  s  S  T...
```

$\Delta G^0_{min}$ = -8.90 kcal/mol + -12.17 kcal/mol  (average: -10.54 kcal/mol)

**MK105.**

```
            |-- net 93 bp loss ------------------------------------------------------------------------------------------------------|
A  TTTGGACTTCATCCGTGACTTCCATCAGCTAGTGAAGGCCCTACCCCAGTATCAGCACTCCTTCATTCCAGTACAATAGTTATAGCTGGAGTATTTACCCTCATCCGCTTTTATCCATTAATAGAAAACAACCTCACTATTCAAACTTCAACACTTCTGGTTCTGGCTCTAATGTCAC
R  TTTGGACTTCAGCCGTTACCATGCGACTTTGGACAGAATGCTTT------------------------------------------------------------------------------CCTGTCGCTGGAGGGCTGATTGGCTCTGACTCTAATGTCAC
P  GGCATACTTCAGCCGTTACCATGCGACTTTGGACAGAATGCTTT------------------------------------------------------------------------------CCTGTCGCTGGAGGGCTGATTGGCTCTGACTCTAATTTACG
Pos     <-1868500-----------------------------------------------------------------------------------------------------------------1868576->
tl   ...T  S  a  V  T  m  r  l  w  t  e  c  f                                                                          p  v  a  g  g  l  i  G  S  d  S  N  V...
```

$\Delta G^0_{min}$ = -14.75 kcal/mol + -16.94 kcal/mol  (average: -15.85 kcal/mol)

**MK141.**

```
                                                                                 |-------| no net loss/gain
A  ACTACTAGGTCTTCTCTTAGCCTCAGCAGGAAAATCAGCCCAATTTGGACTTCATCCGTGACTTCCATCAGCTAGTGAAGGCCCTACCCCAGTATCAGCACTCCTTCATTCCAGTACAATAGTTATAGCTGGAGTATTTACCCTCATCCGCTTTTATCCATTAATAGAAAACAACCTCAC
R  ACTACTAGGTCTTCTCTTAGCCTCAGCAGGAAAATCAGCCCAATTTGGACTTCATCCGTGACTTCCATCTTCCAGTGCAGGCCCTACCCCAGTATCAGCACTCCTTCATTCCAGTACAATAGTTATAGCTGGAGTATTTACCCTCATCCGCTTTTATCCATTAATAGAAAACAACCTCAC
P  GTTGTCTGGTTTCATTTTATAAATCCAGCCTTTGCCATAAGGTTCGTCATTGATAAAGTCTGGATCATCTTCCAGTGCAGTATTGACTTCAACCACTGTACCCGATACTGGTGCATGAATGTCTGAAGCTGTTTTTACAGACTCAACCACGCCAGCTTGTGTGCCAGCAGTCATATGGCT
Pos                                                          1870223<------------->1870209
R translated                                                   ...S  I  f  q  c  R...
```

$\Delta G^0_{min}$ = -7.90 kcal/mol

**MK113.**

```
            |-- net 114 bp loss ------------------------------------------------------------------------------------------------------|
A  TTCATCCGTGACTTCCATCAGCTAGTGAAGGCCCTACCCCAGTATCAGCACTCCTTCATTCCAGTACAATAGTTATAGCTGGAGTATTTACCCTCATCCGCTTTTATCCATTAATAGAAAACAACCTCACTATTCAAACTTCAACACTTCTGGTTCTGGCTCTAATGTCACTAATGAA
R  TTCATCCGTGACTTCCATCGACCAAATGGCTTTCCA------------------------------------------------------------------------------------ACATCACACCTCGAACATCACTAATGAA
P  AAATCTTCTGATTTCCATCGACCAAATGGCTTTCCA------------------------------------------------------------------------------------ACATCACACCTCGAACATCACTTTCAAT
Pos          <-1870516------------------------------------------------------------------------------------------------------1870565->
R      ...V  T  S  I  d  q  m  a  f  q                                                                                   h  h  t  S  N  i  T...
```

$\Delta G^0_{min}$ = -10.65 kcal/mol + -7.70 kcal/mol  (average: -9.18 kcal/mol)

**MK107.**

```
                                                                                                                                                           |-- net 30 bp deleted ----------------------|
A   CTTCTCTTAGCCTCAGCAGGAAAATCAGCCCAATTTGGACTTCATCCGTGACTTCCATCAGCTAGTGAAGGCCCTACCCCAGTATCAGCACTCCTTCATTCCAGTACAATAGTTATAGCTGGAGTATTTACCCTCATCCGCTTTTATCCATTAATAGAAAACAACCTCACTATTCAAACT
R   CTTCTCTTAGCCTCAGCAGGAAAATCAGCCCAATTTGGACTTCATCCGTGACTTCCATCAGCTAGTGAAGGCCCTACCCCAGTATCAGCACTCCTTCATTCCAGTACAATAGTTCAAAAGA------------------------------TCCGATGAAACAACCTCACTATTCAAACT
P   TGCGTGGAGCAAATGATGGACCAAATTATGATATTCAGTCCATTTCGAATATTAGAGCAAAACATAAGGGCGAATTACCTGCGCTTGTAATTGACTGTAGTCATGGCAATAGTTCAAAAGA------------------------------TCCGATGAAACAACCGCAAGTACTTCGAC
Pos                                                                                                                                              <-1870632-----------------------------------------1870597->
R translated                                                                                                                                ...Y  N  S  s  k  d                                  p  m  K  Q  P  H...
```

$\Delta G^0_{min}$ = -7.27 kcal/mol + -12.36 kcal/mol (average: -9.82 kcal/mol)

**MK112.**

```
                                                                                                              |-- net 6 bp loss -|
A   AGCTAGTGAAGGCCCTACCCCAGTATCAGCACTCCTTCATTCCAGTACAATAGTTATAGCTGGAGTATTTACCCTCATCCGCTTTTATCCATTAATAGAAAACAACCTCACTATTCAAACTTCAACACTTCTGGTTCTGGCTCTAATGTCACTAATGAAAAAATGCGTTTCTGGAGTCC
R   AGCTAGTGAAGGCCCTACCCCAGTATCAGCACTCCTTCATTCCAGTACAATAGTTATAGCTGGAGTATTTACCCTCATCCGCTTTTATCCATTAATAGAAAACAACCA-ATGAT-CAA----CAACGCTTCTGGTTCTGGCTCTAATGTCACTAATGAAAAAATGCGTTTCTGGAGTCC
P   AAAAACGCTAAAAATCGTCGATAAACGCTTGTCAAATACTCAAACAACGCGTATGTTAGTTAACAAGATACGTTTTATCTGATTGAAACAGACATGTTAAAACAACCA-ATGAT-CAA----CAACGCTTACTTTTATTTTTGGCAATTTAGTTAGTTGCCTGACTGCATTCGGGCAAC
Pos                                                                                                    <-1871738-------------1871713->
R translated                                                                                                       m  i  n     n  a  S  G...
```

$\Delta G^0_{min}$ = -11.55 kcal/mol + -6.55 kcal/mol (average: -9.05 kcal/mol)

**MK110.**

```
                                                                   |-- net 9 bp loss ---------------------|
A   ACTACTAGGTCTTCTCTTAGCCTCAGCAGGAAAATCAGCCCAATTTGGACTTCATCCGTGACTTCCATCAGCTAGTGAAGGCCCTACCCCAGTATCAGCACTCCTTCATTCCAGTACAATAGTTATAGCTGGAGTATTTACCCTCATCCGCTTTTATCCATTAATAGAAAACAACCTCAC
R   ACTACTAGGTCTTCTCTTAGCCTCAGCAGGAAAATCAGCCCAATTTGGACTTCATCCGTGAC-ACCTTCA-C---CGACTTCAATA--ACAATACCAG--CGCCTTCATTCCAGTACAATAGTTATAGCTGGAGTATTTACCCTCATCCGCTTTTATCCATTAATAGAAAACAACCTCAC
P   CATTCACCACATTCGGCAGTATAAAGCGGAATGACATGATCGCCAGGTTGTACACTGGTGAC-ACCTTCA-C---CGACTTCAATA--ACAATACCAG--CGCCTTCATGTCCTAAAATTGCGGGAAAAACACCTTCAGGATCATCACCAGATAAGGTAAAAGCATCTGTATGGCATACA
Pos                                                         <-1872871--------------------------------1872913->
R translated                                             ...V  T  p  s  p    t  s  I    t  I  p  a    P  S  F...
```

$\Delta G^0_{min}$ = -6.11 kcal/mol + -12.15 kcal/mol (average: -9.13 kcal/mol)

**MK90.**

```
                                                                            |-- net 9 bp loss ------------------|
A   ACTACTAGGTCTTCTCTTAGCCTCAGCAGGAAAATCAGCCCAATTTGGACTTCATCCGTGACTTCCATCAGCTAGTGAAGGCCCTACCCCAGTATCAGCACTCCT-TC-ATTCCAGTACAATAGTTATAGCTGGAGTATTTACCCTCATCCGCTTTTATCCATTAATAGAAAACAACCTCAC
R   ACTACTAGGTCTTCTCTTAGCCTCAGCAGGAAAATCAGCCCAATTTGGACTTCATCCGTGACTTCCATCAGC-AGTGAA-----TA-CTTTG-AT--GC-CAGCTATCAATTCCAGTACAATAGTTATAGCTGGAGTATTTACCCTCATCCGCTTTTATCCATTAATAGAAAACAACCTCAC
P   AATCACCGCATGCATAACAAATTGATTATTGCGGACAACACAATTGCGGTGACTGGAGGACGCAATATCAGC-AGTGAA-----TA-CTTTG-AT--GC-CAGCTATCAATTCCAGTTTACAGATCTCGATATTTTGTTTTATGGTACAGCCGCATCTCAGGCAGAGAAAGTCTTTCAAACT
Pos                                                                   <-1874200--------------------------------1874239->
R translated                                                       ...I  S  s  e     y  f  d     a  s  y  q  F  Q  Y...
```

$\Delta G^0_{min}$ = -13.69 kcal/mol + -8.91 kcal/mol (average: -11.30 kcal/mol)

**MK71.**

```
                                                                        |-- net 6 bp loss -------------|
A   ACTACTAGGTCTTCTCTTAGCCTCAGCAGGAAAATCAGCCCAATTTGGACTTCATCCGTGACTTCCATCAGCTAGTGAAGGCCCTACC-CCAGTATCAGCACTCCTTCATTCCAGTACAATAGTTATAGCTGGAGTATTTACCCTCATCCGCTTTTATCCATTAATAGAAAACAACCTCA
R   ACTACTAGGTCTTCTCTTAGCCTCAGCAGGAAAATCAGCCCAATTTGGACTTCATCCGTGACTTCCA-GACCTAAT-AA-G-CC-ACCACCA--ATCACCACTCCTTCATTCCAGTACAATAGTTATAGCTGGAGTATTTACCCTCATCCGCTTTTATCCATTAATAGAAAACAACCTCA
P   ATCCGATCCATCAAATGCAATACCGTCACATTCATACCGCGCTGTTTTAAACCATAGGCGGCTTCCA-GACCTAAT-AA-G-CC-ACCACCA--ATCACCACAGCATGCTTTTTCTGAGATGCGTAATCCACCATGGTATTGACATCATAGATATCCCGAAAGCTAATGACACCTTTTAA
Pos                                                              <-1898975-------------------------1899010->
R translated                                                  ...T  S  r   p  n  k   p  p  p   I  t  T...
```

$\Delta G^0_{min}$ = -21.89 kcal/mol

**A26.**

```
                                                                        |--------| no net gain/loss
A   ACTACTAGGTCTTCTCTTAGCCTCAGCAGGAAAATCAGCCCAATTTGGACTTCATCCGTGACTTCCATCAGCTAGTGAAGGCCCTACCCCAGTATCAGCACTCCTTCATTCCAGTACAATAGTTATAGCTGGAGTATTTACCCTCATCCGCTTTTATCCATTAATAGAAAACAACCTC
R   ACTACTAGGTCTTCTCTTAGCCTCAGCAGGAAAATCAGCCCAATTTGGACTTCATCCGTGACTTCCACCTGCAGCATAAGGCCCTACCCCAGTATCAGCACTCCTTCATTCCAGTACAATAGTTATAGCTGGAGTATTTACCCTCATCCGCTTTTATCCATTAATAGAAAACAACCTC
P   ATTGACGCCTCCCTGAGCATTTACAGCTTTCATATAATCGATAAATCCACCAAAAAAGCCCGTTCCACCTGCAGCATAAGGCCCTACACGATAACTTTGTAATGGAAAATATTGAGCATTTTTATCAGACAAAACTTTTTCAGCGTTCGATTTATCTGTGTTTGTTTGATCTTGAGAC
Pos                                                                 <-1928206---------1928232->
R translated                                                     ...T  S  t  c  s  i  R  P  Y...
```

$\Delta G^0_{min}$ = -16.91 kcal/mol

**MK30.**

```
                                                         |-- net 9 bp loss ---------------------|
A  ACTACTAGGTCTTCTCTTAGCCTCAGCAGGAAAATCAGCCCAATTTGGACTTCATCCGTGACTTCCATCAGCTAGTGAAGGCCCTACCCCAGTATCAGCACTCCTTCATTCCAGTACAATAGTTATAGCTGGAGTATTTACCCTCATCCGCTTTTATCCATTAATAGAAAACAACCTCAC
R  ACTACTAGGTCTTCTCTTAGCCTCAGCAGGAAAATCAGCCCAATTTGGACTTCATCCGTGAC---------CACAGCATCTGTTTCGGCCCAGTTCGGCACACCTTCATTCCAGTACAATAGTTATAGCTGGAGTATTTACCCTCATCCGCTTTTATCCATTAATAGAAAACAACCTCAC
P  AAAGGCTGTCCTTCGGTTAAACGGATTTCAATTGGTGGTTGATCTGGCACATAAATCGTGAC---------CACAGCATCTGTTTCGGCCCAGTTCGGCACACCTTCATAAATAAATGCAAACAACACCACACGGCGAATCTGCGACCAGTGCTGTCCATTAATCCTAAGCCACTCACCC
Pos                                                        <-1944867-----------------------------------1944910->
R translated                                           ...S  V  T        t  a  s  v  s  a  q  f  g  T  P  S  F...
```

$\Delta G^0_{min}$ = -7.18 kcal/mol + -16.21 kcal/mol (average: -11.70 kcal/mol)

**MK135.**

```
                                                                 |--------------| net 12 bp loss
A  ACTACTAGGTCTTCTCTTAGCCTCAGCAGGAAAATCAGCCCAATTTGGACTTCATCCGTGACTTCCATCAGCTAGTGAAGGCCCTACCCCAGTATCAGCACTCCTTCATTCCAGTACAATAGTTATAGCTGGAGTATTTACCCTCATCCGCTTTTATCCATTAATAGAAAACAACCTCAC
R  ACTACTAGGTCTTCTCTTAGCCTCAGCAGGAAAATCAGCCCAATTTGGACTTCATCCGTGACTTCCATCA-----------CCC-GCCCCAGTATCAGCACTCCTTCATTCCAGTACAATAGTTATAGCTGGAGTATTTACCCTCATCCGCTTTTATCCATTAATAGAAAACAACCTCAC
P  AAAAACGACTTTTTGGTATTGTTGCCCTAGGTCATTGAGCTTCACGATAATTTGTTCATCATCGCCATCA-----------CCC-GCCCCAGTACGATTATCACCCGTTAACCAGATATGTCCAGAAGAATGCTTTTGATTATTAAAAAAGATAACATCACTATTTTCTAAAGTGGCATT
Pos                                                                <-1956396------------1956413->
R translated                                                   ...S  I  t             r  P  S  I...
```

$\Delta G^0_{min}$ = -5.87 kcal/mol + -14.07 kcal/mol (average: -9.97 kcal/mol)

**MK79.**

```
                                                          |-- net 21 bp loss -------------------------|
A  ACTACTAGGTCTTCTCTTAGCCTCAGCAGGAAAATCAGCCCAATTTGGACTTCATCCGTGACTTCCATCAGCTAGTGAAGGCCCTACCCCAGTATCAGCACTCCTTCATTCCAGTACAATAGTTATAGCTGGAGTATTTACCCTCATCCGCTTTTATCCATTAATAGAAAACAACCTCAC
R  ACTACTAGGTCTTCTCTTAGCCTCAGCAGGAAAATCAGCCCAATTTGGACTTCATCCGTGACTGCAA-CAG--A-TCATG-C---A-----GT--CA-CT-TC-T---TTCCAGTACAATAGTTATAGCTGGAGTATTTACCCTCATCCGCTTTTATCCATTAATAGAAAACAACCTCAC
P  CGTGCAGTTGTGATTAGTACCGGCATTCAAACCTATTTTGGTGCGCTGGCTCATCGCGTGACTGCAA-CAG--A-TCATG-C---A-----GT--CA-CT-TC-T---TTCCAGTTGGGTGTCAATAAAGTCAGTTGGTTATTGATTCGCTTTATGTTGGTCATGGCACCAGTCGTCTTG
Pos                                                        <-1975009-----------------------------------------1974966->
R translated                                           ...S  V  T  a  t  d    h  a          v    t    s     F  Q  Y...
```

$\Delta G^0_{min}$ = -8.89 kcal/mol + -7.84 kcal/mol (average: -8.37 kcal/mol)

**MK138.**

```
                                                                  |--------------| net 3 bp loss
A  ACTACTAGGTCTTCTCTTAGCCTCAGCAGGAAAATCAGCCCAATTTGGACTTCATCCGTGACTTCCATCAGCTAGTGAAGGCCCTACCCCAGTATCAGCACTCCTTCATTCCAGTACAATAGTTATAGCTGGAGTATTTACCCTCATCCGCTTTTATCCATTAATAGAAAACAACCTCAC
R  ACTACTAGGTCTTCTCTTAGCCTCAGCAGGAAAATCAGCCCAATTTGGACTTCATCCGTGACTTCCATCAG---GTACAGGAACAATCCCAGTATCAGCACTCCTTCATTCCAGTACAATAGTTATAGCTGGAGTATTTACCCTCATCCGCTTTTATCCATTAATAGAAAACAACCTCAC
P  AAATTCCATTGTGGTAGGTAAGATTCGCTGAGGTTTTACAATTGCTGCAAGTTTGTCTAAGCGTGCATCAG---GTACAGGAACAATCCCAGTATTTGGCTCAATGGTACAAAATGGAAAGTTTTCTGCTGCAATCGCTGCTTTGGTCAATGCATTAAACAGAGTTGATTTACCAACGTT
Pos                                                              <-2054979----------------2054949->
R translated                                                 ...T  S  I  r     y  r  n  n  P  S  I...
```

$\Delta G^0_{min}$ = -16.93 kcal/mol

**MK140.**

```
                                                                        |-| no net loss/gain
A  ACTACTAGGTCTTCTCTTAGCCTCAGCAGGAAAATCAGCCCAATTTGGACTTCATCCGTGACTTCCATCAGCTAGTGAAGGCCCTACCCCAGTATCAGCACTCCTTCATTCCAGTACAATAGTTATAGCTGGAGTATTTACCCTCATCCGCTTTTATCCATTAATAGAAAACAACCTCAC
R  ACTACTAGGTCTTCTCTTAGCCTCAGCAGGAAAATCAGCCCAATTTGGACTTCATCCGTGACTTCCATCAGCTATTCAAGGCCCTACCCCAGTATCAGCACTCCTTCATTCCAGTACAATAGTTATAGCTGGAGTATTTACCCTCATCCGCTTTTATCCATTAATAGAAAACAACCTCAC
P  ATGGCACGGTAAAACGAACATGATGATATGCCTGACTTAGAAAAGCACAATATTGTTCCAGTGAAAGACGGCTATTCAAGGCATCATCAATCAGTGAAGTCGCAAACAATTTTTGTCTTTCTTGAGCGGTTGCTTGTTGAAGCTGTTGAAAAAATGACATAAATTTCTTCCTTTAAGCTG
Pos                                                            2095646<------------>2095659
R translated                                                     ...I  S  y  s  R  P...
```

$\Delta G^0_{min}$ = -11.86 kcal/mol

**MK74.**

```
                                                                        |-| no net loss/gain
A  ACTACTAGGTCTTCTCTTAGCCTCAGCAGGAAAATCAGCCCAATTTGGACTTCATCCGTGACTTCCATCAGCTAGTGAAGGCCCTACCCCAGTATCAGCACTCCTTCATTCCAGTACAATAGTTATAGCTGGAGTATTTACCCTCATCCGCTTTTATCCATTAATAGAAAACAACCTCAC
R  ACTACTAGGTCTTCTCTTAGCCTCAGCAGGAAAATCAGCCCAATTTGGACTTCATCCGTGACTTCCATCAGCTATTTAAGGCCCTACCCCAGTATCAGCACTCCTTCATTCCAGTACAATAGTTATAGCTGGAGTATTTACCCTCATCCGCTTTTATCCATTAATAGAAAACAACCTCAC
P  TTGGTAGGCATATCCAGACTCGAACTGGAAACCTCTACGATGTCAACGTAGTGTTTAAGCTTATAATTCAGCTATTTAAGAAGTATGGGTGTCAAATTGGTGTCGATAAATTTATGTAAGGAGAAGTTTCATATCTAAAAGGTAGAGAGAATAATGATCTAATTGTTATTGAGCTATATT
Pos                                                           2120356<----------->2120368
R translated                                                     ...I  S  y  l  S...
```

$\Delta G^0_{min}$ = -10.35 kcal/mol

**A17.**

```
                                                                         |----------------| net 9 bp loss
A   ACTACTAGGTCTTCTCTTAGCCTCAGCAGGAAAATCAGCCCAATTTGGACTTCATCCGTGACTTCCATCAGCTAGTGAAGGCCCTACCCCAGTATCAGCACTCCTTCATTCCAGTACAATAGTTATAGCTGGAGTATTTACCCTCATCCGCTTTTATCCATTAATAGAAAACAACCTCAC
R   ACTACTAGGTCTTCTCTTAGCCTCAGCAGGAAAATCAGCCCAATTTGGACTTCATCCGTGACTTCCATCAGCT--TGAAG-------CCGTGTATCAGCACTCCTTCATTCCAGTACAATAGTTATAGCTGGAGTATTTACCCTCATCCGCTTTTATCCATTAATAGAAAACAACCTCAC
P   CAAATATTGGCACATCGAGAGATAAAGCGGCCCATGGGCACTTTGTTCGCCGTGATATACCCGATGATCAGCT--TGAAG-------CCGTGTATCAGCACTGGCTGGCAAAACGTATTGTGAACCGCCCAGCAAGCGACATGCTGCGTGCCGGCTGGTTTTATGAAGGCATTCAGGATG
Pos                                                                     <-2152623------------------2152597->
R translated                                                         ...I  S  l    k         P  c  I  S  T  P...
```

$\Delta G^0_{min}$ = -13.48 kcal/mol + -15.82 kcal/mol  (average: -14.83 kcal/mol)

**MK6.**

```
                  |-- net 123 bp loss -----------------------------------------------------------------------------------------------------|
A   CAGGAAAATCAGCCCAATTTGGACTTCATCCGTGACTTCCATCAGCTAGTGAAGGCCCTACCCCAGTATCAGCACTCCTTCATTCCAGTACAATAGTTATAGCTGGAGTATTTACCCTCATCCGCTTTTATCCATTAATAGAAAACAACCTCACTATTCAAACTTCAACACTTCTGGTTCTG
R   CAGGAAAATCAGCGCAA------------------------------------------------------------------------------------------------------------AAAAACCTATGCCGCGTTCAAACTTCAACACTTCTGGTTCTG
P   AGCAAAAATCAGCGCAA------------------------------------------------------------------------------------------------------------AAAAACCTATGCCGCGTTCAAACTTCATAAATCTGTCATAAA
Pos     <-2204361-----------------------------------------------------------------------------------------------------------------------2204322->
R     ...K  I  S  a  k                                                                                                     K  p  m  p  r  S  N  F  N...
```

$\Delta G^0_{min}$ = -15.68 kcal/mol + -13.55 kcal/mol  (average: -14.62 kcal/mol)

**MK100.**

```
                                                                        |-- net 3 bp inserted ------------|
A   ACTACTAGGTCTTCTCTTAGCCTCAGCAGGAAAATCAGCCCAATTTGGACTTCATCCGTGACTTCCATCAGCTAGTGAAGGCCCTA---CCCCAG-TATCAGCACTCCTTCATTCCAGTACAATAGTTATAGCTGGAGTATTTACCCTCATCCGCTTTTATCCATTAATAGAAAACAACCTCAC
R   ACTACTAGGTCTTCTCTTAGCCTCAGCAGGAAAATCAGCCCAATTTGGACTTCATCCGTGACTTCCATCAGCCAATGCAG-CAATAGATCCCCATACATCGCCACTGCTTCATTCCAGTACAATAGTTATAGCTGGAGTATTTACCCTCATCCGCTTTTATCCATTAATAGAAAACAACCTCAC
P   GCTTGGACCTAAGCGGGCAAACTTGATCATCAAATACATTTCAATAACTAAAAGAACCGTATAGAAAGCAGCCAATGCAG-CAATAGATCCCCATACATCGCCACTGCTTAAACTAGATGTTGAAAGATGTGTTGGAAGAACTTCGCCAATCGTCCACGGTTGACGGCCACCTTCTGCAACGTA
Pos                                                                     <-2263122-------------------------2263080->
R translated                                                         ...I  S  q  c  s   n  r  s  p  y  i  a  t  a  S...
```

$\Delta G^0_{min}$ = -19.02 kcal/mol

**MK17.**

```
                                                             |-- net 162 bp loss -----------------------------------------------|
A   CTACACTCCCACTACTAGGTCTTCTCTTAGCCTCAGCAGGAAAATCAGCCCACCAATTTGGACTTCATCCGTGACTTCCATCAGCTAGTGAAGG(120xN)TCTGGCTCTAATGTCACTAATGAAAAAATGCGTTTCTGGAGTCCAGAGGTTCGTGAGTTAGAGCCTTATGTACCTGGTG
R   CTACACTCCCACTACTAGGTCTTCTCTTAGCCTCAGCAGGAAAATCAGCCCAC-------------------------------------------------TGTGTACCCTCAAGCAACGTGAAAAAATGCGTTTCTGGAGTCCAGAGGTTCGTGAGTTAGAGCCTTATGTACCTGGTG
P   TCAAAGTATCAAGATGGGCTTACCCGATGATATGCAAACCGAATACGGCCCAC-------------------------------------------------TGTGTACCCTCAAGCAACGTGAAAAAATCCAGCAAGTGGTGCAGCGTTCAGTTGAACAAGGTGCAAAACTGATCACTG
Pos                                                       <-2502313------------------------------------------------------------2502276->
R translated                                           ...I  S  P  f                                             c  t  l  k  q  r  E  K  M  R...
```

$\Delta G^0_{min}$ = -9.25 kcal/mol + -9.95 kcal/mol  (average: -9.60 kcal/mol)

**MK21.**

```
                    |-- net 123 bp loss -----------------------------------------------------------------------------------------------------|
A   TCAGCCCAATTTGGACTTCATCCGTGACTTCCATCAGCTAGTGAAGGCCCTACCCCAGTATCAGCACTCCTTCATTCCAGTACAATAGTTATAGCTGGAGTATTTACCCTCATCCGCTTTTATCCATTAATAGAAAACAACCTCACTATTCAAACTTCAACACTTCTGGTTCTGGCTCTAA
R   TCAGCCCAATTTGGACGTC-----------------------------------------------------------------------------------------------------------CGACAATAGCGACAAAACCACTTCTGGTTCTGGCTCTAA
P   TTTCCAACATTTGGACGTC-----------------------------------------------------------------------------------------------------------CGACAATAGCGACAAAACCACTTCTGAAGTCGGAAGGAA
Pos         <-2538818-----------------------------------------------------------------------------------------------------------------------2538854->
R tl     ...I  W  T  S                                                                                                        d  n  s  d  k  t  T  S  G...
```

$\Delta G^0_{min}$ = -11.23 kcal/mol + -9.51 kcal/mol  (average: -10.37 kcal/mol)

**MK70.**

```
                                                                         |-- net 9 bp inserted ---------|
A   ACTACTAGGTCTTCTCTTAGCCTCAGCAGGAAAATCAGCCCAATTTGGACTTCATCCGTGACTTCCATCA-GC-TAG-TGAAGGC--CC-T--A-CCCCAGTATCAGCACTCCTTCATTCCAGTACAATAGTTATAGCTGGAGTATTTACCCTCATCCGCTTTTATCCATTAATAGAAAA
R   ACTACTAGGTCTTCTCTTAGCCTCAGCAGGAAAATCAGCCCAATTTGGACTTCATCCGTGACTTCCATCAACCATGGACCAAACCAACCTTGCATACCAAAAATCAGCACTCCTTCATTCCAGTACAATAGTTATAGCTGGAGTATTTACCCTCATCCGCTTTTATCCATTAATAGAAAA
P   ACGGTAATAAAAATGGTTGCAATTACAATCGCAGGTACGGCGTATAAAATCTTAATGTCATGATCCATCAACCATGGACCAAACCAACCTTGCATACCAAAAATCAGCACTAACATTAAACCTGCGATGACAGGTGAAACTGAAAATGGCAAATCAATCAGCGTGGTCAGAAAAGATTTA
Pos                                                                    <-2552426------------------------------2552379->
R translated                                                        ...S  I  n  h  g  p  n  q  p  c  i  P  k  I  S  T...
```

$\Delta G^0_{min}$ = -7.66 kcal/mol + -11.53 kcal/mol  (average: -9.60 kcal/mol)

```
MK119.                                                                     |- net 12 bp inserted-|
A  ACTACTAGGTCTTCTCTTAGCCTCAGCAGGAAAATCAGCCCAATTTGGACTTCATCCGTGACTTCCATCAGCTAGTG-AAG--------G-CC--CTACCCCAGTATCAGCACTCCTTCATTCCAGTACAATAGTTATAGCTGGAGTATTTACCCTCATCCGCTTTTATCCATTAATAGAA
R  ACTACTAGGTCTTCTCTTAGCCTCAGCAGGAAAATCAGCCCAATTTGGACTTCATCCGTGACTTCCATCAGCAGGTGCAAGATTACATTGTCCTTCTACACCAGTATCAGCACTCCTTCATTCCAGTACAATAGTTATAGCTGGAGTATTTACCCTCATCCGCTTTTATCCATTAATAGAA
P  TAAAAGTCGAGGCTTAAAGTATCCTAAAAACGCTTTCGTGGGGATACTGTATGGGCACAACAATCCATCAGCAGGTGCAAGATTACATTGTCCTTCTACACCAGTTCCTGCAAAAAGTTTGTTTGAGCCAATATTAATATTCGGGTAGTTAGAATTCTTTGTGGCGCAGTCATTATTAACT
Pos                                                                 <-2584138------------------------2584179->
R translated                                                     ...S  I  S  r  c  k  I  t  l  s  f  Y  t  I...
```

**ΔG⁰min = -16.07 kcal/mol + -10.20 kcal/mol (average: -13.14 kcal/mol)**

```
R1522.                                                                     |-- net 3 bp inserted --------|
A  ACTACTAGGTCTTCTCTTAGCCTCAGCAGGAAAATCAGCCCAATTTGGACTTCATCCGTGACTTCCATCAGCTAGTGA---AG---GCCCTACCCCAGTATCAGCACTCCTTCATTCCAGTACAATAGTTATAGCTGGAGTATTTACCCTCATCCGCTTTTATCCATTAATAGAAAACA
R  ACTACTAGGTCTTCTCTTAGCCTCAGCAGGAAAATCAGCCCAATTTGGACTTCATCCGTGACTTCCATCAGCAAGTGGCACAGATTGCCAAAC---AGTATCTGCACTCCTTCATTCCAGTACAATAGTTATAGCTGGAGTATTTACCCTCATCCGCTTTTATCCATTAATAGAAAACA
P  TACTTTCGATTTTTGGATTTATATTAATTCAATTATTAGCATTTTTGTGTCCTTGGCTCATTGGCGATCAGCAAGTGGCACAGATTGCCAAAC---AGTATCTGCACTTTGTTGCTTTTGGTGTACCAGGTTTAGTTTATATGACACTATTTCGCTTCGTTAGCGAAGGTCATAGTTTA
Pos                                                                   <-2700471------------------------2700433->
R translated                                                       ...I  S  k  w  h  r  l  p  n     S  I  c  T...
```

**ΔG⁰min = -10.41 kcal/mol + -11.91 kcal/mol (average: -11.16 kcal/mol)**

```
MK3.                                                                      |-------| no net gain/loss
A  ACTACTAGGTCTTCTCTTAGCCTCAGCAGGAAAATCAGCCCAATTTGGACTTCATCCGTGACTTCCATCAGCTAGTGAAGGCCCTACCCCAGTATCAGCACTCCTTCATTCCAGTACAATAGTTATAGCTGGAGTATTTACCCTCATCCGCTTTTATCCATTAATAGAAAACAACCTCAC
R  ACTACTAGGTCTTCTCTTAGCCTCAGCAGGAAAATCAGCCCAATTTGGACTTCATCCGTGACTTCCATCAGTTATCAAATGCCCTACCCCAGTATCAGCACTCCTTCATTCCAGTACAATAGTTATAGCTGGAGTATTTACCCTCATCCGCTTTTATCCATTAATAGAAAACAACCTCAC
P  AGTCTCAATCTAAACTTGAGTCATTCTTTAGATGACTGGGAAAAGGATCGTAATCAAAGTTATTACATCAGTTATCAAATGCCCTTTGGATACTACGATTTTGGTCTAAGTTATAACTGGTATCGCTATGATCAGTATGTTGCGGGCTTTAATGCACCGATTCATTATCAGGGTGAAAGC
Pos                                                              <-2736164-----2736142->
R translated                                                 ...T  S  I  S  f  q  m  P  Y...
```

**ΔG⁰min = -16.30 kcal/mol**

```
MK121.                                                                                                                  |-- net 17 bp insertion -------------------|
A  AGCTAGTGAAGGCCCTACCCCAGTATCAGCACTCCTTCATTCCAGTACAATAGTTATAGCTGGAGTATTTACCCTCATCCGCTTTTATCCATTAATAGAAAACAACCTCACTATTCAAACTTCAACACTTC--T--GGT-T-CTG--GC-T---C-T----AATGTCACTAATGAAAAAATG
R  AGCTAGTGAAGGCCCTACCCCAGTATCAGCACTCCTTCATTCCAGTACAATAGTTATAGCTGGAGTATTTACCCTCATCCGCTTTTATCCATTAATAGAAAACAACCTCACTATTCATCCAGCAACTCTTGAGTACGGAATACTGCAGCATGATCCTGCACAATGTCACTAATGAAAAAATG
P  TACAGGTTTTCAACCTCCAAAGCCTCTACACGTGCAGTGTTAAATACCTTAGATTTGTCTTTTAAGTAAATATTGCGAACTCGTGGCTCTAGCGCAAGAATTTGTCTTACACCTTCATCCAGCAACTCTTGAGTACGGAATACTGCAGCATGATCCTGCACAATGTCACGAATCGCATCTGC
Pos                                                                                                             <-2821959----------------------------------------2822018->
R translated                                                                                                                                        m  i  l  h  N  V  T  N...
```

**ΔG⁰min = -5.95 kcal/mol + -11.63 kcal/mol (average: -8.79 kcal/mol)**

```
MK24.                                                                       |--------------| net 3 bp loss
A  ACTACTAGGTCTTCTCTTAGCCTCAGCAGGAAAATCAGCCCAATTTGGACTTCATCCGTGACTTCCATCAGCTAGTGAAGGCCCTACCCCAGTATCAGCACTCCTTCATTCCAGTACAATAGTTATAGCTGGAGTATTTACCCTCATCCGCTTTTATCCATTAATAGAAAACAACCTCAC
R  ACTACTAGGTCTTCTCTTAGCCTCAGCAGGAAAATCAGCCCAATTTGGACTTCATCCGTGACTTCCATCAGCTCGTGCATG---TTTGGCAGTATCAGCACTCCTTCATTCCAGTACAATAGTTATAGCTGGAGTATTTACCCTCATCCGCTTTTATCCATTAATAGAAAACAACCTCAC
P  TTAATATTGCGGATTGGTGTGATGATTTTTCGACACAATCGATTGAGAAACAGTCTTGGCTGATTCATGAGCTCGTGCATG---TTTGGCAGTATCAGCAAGGCATAAAGTTAATCCGAAAAGGAATTTTTGAAAGAAAATATCAGTACGTTTTGCAGCAGGGAAAGCAGTTTTTGCATT
Pos                                                                 <-2826900-------------------2826867->
R translated                                                     ...I  S  s  c  m     f  g  S  I  S  T...
```

**ΔG⁰min = -11.84 kcal/mol + -14.34 kcal/mol (average: -13.09 kcal/mol)**

```
MK133.                                                                       |----| no net gain/loss
A  ACTACTAGGTCTTCTCTTAGCCTCAGCAGGAAAATCAGCCCAATTTGGACTTCATCCGTGACTTCCATCAGCTAGTGAAGGCCCTACCCCAGTATCAGCACTCCTTCATTCCAGTACAATAGTTATAGCTGGAGTATTTACCCTCATCCGCTTTTATCCATTAATAGAAAACAACCTCAC
R  ACTACTAGGTCTTCTCTTAGCCTCAGCAGGAAAATCAGCCCAATTTGGACTTCATCCGTGACTTCCATCAGCTATTCAAAGCCCTACCCCAGTATCAGCACTCCTTCATTCCAGTACAATAGTTATAGCTGGAGTATTTACCCTCATCCGCTTTTATCCATTAATAGAAAACAACCTCAC
P  GAGCATGGATTTTGTAAAAGGGTTACCTACAGGTGTGTATGTAGGGCAAACCAGAATTCAGCATCCTAAAGCTATTCAAAGCCCTTTATCAGAGCTTGAAGGTTTTAACTTAGCCTTGCATGTGAATGATGATCCGTTTCGTGTACAGCAGCATCGTATGATTTTGCTGGATGAGTTTTC
Pos                                                            2834948<--------------->2834933
R translated                                                        ...S  y  s  k  P  Y...
```

**ΔG⁰min = -15.05 kcal/mol**

**MK78.**

```
                                                                            |----------------| no net gain/loss
A  ACTACTAGGTCTTCTCTTAGCCTCAGCAGGAAAATCAGCCCAATTTGGACTTCATCCGTGACTTCCATCAGCTAGTGAAGGC-CCT-ACCCCAGTATCAGCACTCCTTCATTCCAGTACAATAGTTATAGCTGGAGTATTTACCCTCATCCGCTTTTATCCATTAATAGAAAACAACCTCAC
R  ACTACTAGGTCTTCTCTTAGCCTCAGCAGGAAAATCAGCCCAATTTGGACTTCATCCGTGACTTCCATCAGCT-GTT-CGGCAAATGACACCAGTATCAGCACTCCTTCATTCCAGTACAATAGTTATAGCTGGAGTATTTACCCTCATCCGCTTTTATCCATTAATAGAAAACAACCTCAC
P  ACGTAAACGCCAGGTGCATCGGCTTTGGCATAAGCGTCTATATTGTATTTTTTAATCAAATCAATCATCAGCT-GTT-CGGCAAATGACACCAGTGTACGTACATTCCATTTCAGGCGATTTAAATCAAACATAAAATAAATCACCATTTGACCAGGGCCATGCCAAGTGACTTGACCGCCA
Pos                                                                   <-2857228------------2857201->
R translated                                                      ...S  I  S   v  r  q  m  t  P  V...
```

$\Delta G^0_{min}$ = -17.13 kcal/mol

**MK99.**

```
                                   |-- net 48 bp loss -------------------------------------------------|
A  ACTACTAGGTCTTCTCTTAGCCTCAGCAGGAAAATCAGCCCAATTTGGACTTCATCCGTGACTTCCATCAGCTAGTGAAGGCCCTACCCCAGTATCAGCACTCCTTCATTCCAGTACAATAGTTATAGCTGGAGTATTTACCCTCATCCGCTTTTATCCATTAATAGAAAACAACCTCAC
R  ACTACTAGGTCTTCTCTTAGCCTCAGCAGTCCAAACACA------------------------------------------------GCAGGAATCACCACTCCTTCATTCCAGTACAATAGTTATAGCTGGAGTATTTACCCTCATCCGCTTTTATCCATTAATAGAAAACAACCTCAC
P  AGTGTACTCAACGCATGCCCGAGCAGCAGTCCAAACACA------------------------------------------------GCAGGAATCACCACTCTTCCACGGATCCCCGCTGTGATGGCATAAGCCCAATCGGCGCCGGGAGTTAAAATAAACAACGCGGAAACACCCCAA
Pos                           <-2909057-------------------------------------------------------------------2909023->
R translated         ...L  S  s  p  n  t                                                        a  g  I  t  T  L  S...
```

$\Delta G^0_{min}$ = -6.63 kcal/mol + -14.85 kcal/mol (average: -10.74 kcal/mol)

**A71.**

```
                                                                         |-------| no net loss/gain
A  ACTACTAGGTCTTCTCTTAGCCTCAGCAGGAAAATCAGCCCAATTTGGACTTCATCCGTGACTTCCATCAGCTAGTGAAGGCCCTACCCCAGTATCAGCACTCCTTCATTCCAGTACAATAGTTATAGCTGGAGTATTTACCCTCATCCGCTTTTATCCATTAATAGAAAACAACCTCAC
R  ACTACTAGGTCTTCTCTTAGCCTCAGCAGGAAAATCAGCCCAATTTGGACTTCATCCGTGACTTCCATCTTCAAGTGCAGGCCCTACCCCAGTATCAGCACTCCTTCATTCCAGTACAATAGTTATAGCTGGAGTATTTACCCTCATCCGCTTTTATCCATTAATAGAAAACAACCTCAC
P  ACAGTTACCACTGGTTTGAATGGCGTGCATTTGTACCGTAGCAAGCTCTGCCAAAATATCGGGTACATCTTCAAGTGCAGGCCAGTTGAGCTGAATATTTTGACGCGTAGAAACGTGTGCATAACCACGGTCATATTCTTTTGAAATTTCTGCAATTTTGCGTAATTGTTTTGAATTCAT
Pos                                                                   <----------------> 2911491-2911508
R translated                                                      ...S  I  f  k  c  R  P...
```

$\Delta G^0_{min}$ = -11.31 kcal/mol

**MK136.**

```
                          |-- net 123 bp loss ---------------------------------------------------------------------------------------------------------------------|
A  TAGAAAGATGAAACACTACACTCCCACTACTAGGTCTTCTCTTAGCCTCAGCAGGAAAATCAGCCCAATTTGGACTTCATCCGTGACTTCCATCAGCTAGTGAAGGCCCTACCCCAGTATCAGCACTCCTTCATTCCAGTACAATAGTTATAGCTGGAGTATTTACCCTCATCCGCTTTTA
R  TAGAAAGATGAAACACTACCTTTTAC---------------------------------------------------------------------------------------------------------------AAAAAAACCGTATTTACCCTCATCCGCTTTTA
P  ATAGAGGATTGAACACTACCTTTTAC---------------------------------------------------------------------------------------------------------------AAAAAAACCGTATTTACCATAATAGTAGATGT
Pos                  <-2948975---------------------------------------------------------------------------------------------------------------------2948943->
R tl      M  K  H  Y  l  l  q                                                                                                                 k  n  r  I  Y  P...
```

$\Delta G^0_{min}$ = -8.33 kcal/mol + -10.16 kcal/mol (average: -9.25 kcal/mol)

**MK66.**

```
               |-- net 105 bp loss -----------------------------------------------------------------------------------------|
A  CACTACACTCCCACTACTAGGTCTTCTCTTAGCCTCAGCAGGAAAATCAGCCCAATTTGGACTTCATCCGTGACTTCCATCAGCTAGTGAAGGCCCTACCCCAGTATCAGCACTCCTTCATTCCAGTACAATAGTTATAGCTGGAGTATTTACCCTCATCCGCTTTTATCCATTAATAGAA
R  CACTACACTCCCCATGCCCGA---------------------------------------------------------------------------------------------------CTACTGGATTGTAGCTGGAGTATTTACCCTCATCCGCTTTTATCCATTAATAGAA
P  ACAAACACTCCCCATGCCCGA---------------------------------------------------------------------------------------------------CTACTGGATTGTAGCTGGTGATCGATATAAATCGCACGTACTTTATGAGGAAACA
Pos     <-2964498------------------------------------------------------------------------------------------------------------------2964532->
tl ...Y  T  P  h  a  r                                                                                                        l  l  d  c  S  W...
```

$\Delta G^0_{min}$ = -10.60 kcal/mol + -9.36 kcal/mol (average: -9.98 kcal/mol)

**MK63.**

```
                                      |-- net 33 bp loss ---------------------------------------------|
A  ACTACTAGGTCTTCTCTTAGCCTCAGCAGGAAAATCAGCCCAATTTGGACTTCATCCGTGACTTCCATCAGCTAGTGAAGGCCCTACCCCAGTATCAGCACTCCTTCATTCCAGTACAATAGTTATAGCTGGAGTATTTACCCTCATCCGCTTTTATCCATTAATAGAAAACAACCTCAC
R  ACTACTAGGTCTTCTCTTAGCCTCAGCAGGAATATTACGAATATTGTA---------------------------------TTTCACAGCCGTATCTTCACTCCTTCATTCCAGTACAATAGTTATAGCTGGAGTATTTACCCTCATCCGCTTTTATCCATTAATAGAAAACAACCTCAC
P  TTTACCGTCTTTGAATAGAAGAAGTGCAGGAATATTACGAATATTGTA---------------------------------TTTCACAGCCGTATCTTCACATGCAGTGACGTCAACTTTTACAATTTTTACTTTACCTTCGTATTCACGAGACAAATCTTCTAAAACAGGTGCGATAGC
Pos                           <-2967553---------------------------------------------------------2967511->
R translated            ...S  R  n  i  t  n  I  v                                         f  h  s  r  I  f  T...
```

$\Delta G^0_{min}$ = -8.30 kcal/mol + -7.84 kcal/mol (average: -8.07 kcal/mol)

**MK108.**

```
                                            |-- net 45 bp loss ---------------------------------------------|
A  ACTACTAGGTCTTCTCTTAGCCTCAGCAGGAAAATCAGCCCAATTTGGACTTCATCCGTGACTTCCATCAGCTAGTGAAGGCCCTACCCCAGTATCAGCACTCCTTCATTCCAGTACAATAGTTATAGCTGGAGTATTTACCCTCATCCGCTTTTATCCATTAATAGAAAACAACCTCAC
   ||||||||||||||||||||||||||||||||||||||                                                  | | ||    |  | ||||||||||||||||||||||||||||||||||||||||||||||||||||||||||||||||||||||||||||||||||||||
R  ACTACTAGGTCTTCTCTTAGCCTCAGCAGGAAAATCAGGATGG---------------------------------------------CAAATGTCGAAATCCATTCATTCCAGTACAATAGTTATAGCTGGAGTATTTACCCTCATCCGCTTTTATCCATTAATAGAAAACAACCTCAC
     |       |   |      |    | |||  |||||||||||||                                          ||||||||||||||||||||||||||    |    |    | |    |       | | |   | |     || ||     |         | || |
P  CCTGTACCAACAATTAACGTCAGTAACAGTAAAATCAGGATGG---------------------------------------------CAAATGTCGAAATCCATTCATTCCAGATATAACCTAGAAACCCTAGGCCAGATCTTTGTTCTTGATTTTCTTGCATATCTGTTCCACTTTTT
Pos                                <-3115252------------------------------------------------------------------3115294->
R translated             ...S  R  K  I  r  m  a                                              n  v  e  i  h  S  F  Q...
```

$\Delta G^0_{min}$ = -12.04 kcal/mol + -12.53 kcal/mol  (average: -12.29 kcal/mol)

**MK129.**

```
                                  |-- net 33 bp loss -------------------------------|
A  ACTACTAGGTCTTCTCTTAGCCTCAGCAGGAAAATCAGCCCAATTTGGACTTCATCCGTGACTTCCATCAGCTAGTGAAGGCCCTACCCCAGTATCAGCACTCCTTCATTCCAGTACAATAGTTATAGCTGGAGTATTTACCCTCATCCGCTTTTATCCATTAATAGAAAACAACCTCAC
   |||||||||||||||||||||||||||| ||||                                   ||  ||||||||||||||||||||||||||||||||||||||||||||||||||||||||||||||||||||||||||||||||||||||||||||||||||||||||||||||||||
R  ACTACTAGGTCTTCTCTTAGCCTCAGCAGCAAAAAGCC---------------------------------GAAGAAAAGGCCCTACCCCAGTATCAGCACTCCTTCATTCCAGTACAATAGTTATAGCTGGAGTATTTACCCTCATCCGCTTTTATCCATTAATAGAAAACAACCTCAC
   |  |  | |    ||     |    ||||||||||||||                                ||||||||||||| ||           | |          ||          |      |    |    |    | |    |      |     |    | | |     ||
P  AAGAACAAGAGCGCTTAGCACGAGAGCAGCAAAAAGCC---------------------------------GAAGAAAAGGCCCAACGTGATCGAGAAAAACTGACTCAGGATGTGATTGGCACTTTTGCAAAAAGTGCGGCACGAACTCTTGGAGGTCCGACAGGTCAGAAACTGGTAC
Pos                           <-3138850----------------------------------------------3138882->
R translated             ...S  S  K  k  p                                       k  k  R  P  Y...
```

$\Delta G^0_{min}$ = -10.20 kcal/mol + -10.72 kcal/mol  (average: -10.46 kcal/mol)

**MK36.**

```
                                  |-- net 27 bp inserted ---------------------------------------------------------------------------|
A  ACTACTAGGTCTTCTCTTAGCCTCAGCAGGAAAATCAGCCCAATTTGGACTTCATCCGTGACTTCC--------------------------ATCAGCTAGTGAAGGCCCTACCCCAGTATCAGCACTCCTTCATTCCAGTACAATAGTTATAGCTGGAGTATTTACCCTCATCCGCTT
   |||||||||||||||||||||||||||| ||||||  |          |    |      |                              |||          || ||   |    |    ||     ||||||||||||||||||||||||||||||||||||||||||||||||
R  ACTACTAGGTCTTCTCTTAGCCTCAGCAGAAAAATCGCCAATCCCACAGCCAGACTAAATGCAGTGGTAAATCTGGAATCCACATGGCACAGCTCACACCAATCAGACCTGCCAATAAATTTCCAACGATCACATTCCAGTACAATAGTTATAGCTGGAGTATTTACCCTCATCCGCTT
         |     ||| |    | |||  ||||||||||||||||||||||||||||||||||||||||||||||||||||||||||||||||||||||||||||||||||||||||||||||||||||||||||         |
P  GGATGCAATGAATCTGTACTCATCATCAGAAAAATCGCCAATCCCACAGCCAGACTAAATGCAGTGGTAAATCTGGAATCCACATGGCACAGCTCACACCAATCAGACCTGCCAATAAATTTCCAACGATCACATTCCAGGGTTGAGCCAGGGGACTTGATGGCACAGCAAAAAGTAAA
Pos                     <-3179683------------------------------------------------------------------------------------------------3179803->
R translated       ...S  L  S  R  K  I  a  n  p  t  a  r  l  n  a  v  v  k  s  g  i  h  m  a  q  l  t  p  i  r  p  a  n  k  f  p  t  i  t  F  Q...
```

$\Delta G^0_{min}$ = -12.37 kcal/mol + -9.05 kcal/mol  (average: -10.71 kcal/mol)

**MK58.**

```
                                                                           |- net 3 bp inserted -|
A  ACTACTAGGTCTTCTCTTAGCCTCAGCAGGAAAATCAGCCCAATTTGGACTTCATCCGTGACTTCCATCAGCTAG-TGAAGGCCCTAC-CCCAGT-ATCAGCACTCCTTCATTCCAGTACAATAGTTATAGCTGGAGTATTTACCCTCATCCGCTTTTATCCATTAATAGAAAACAACCT
   ||||||||||||||||||||||||||||||||||||||||||||||||||||||||||||||||||||||||  |  |||    |  | |      | |||||||||||||||||||||||||||||||||||||||||||||||||||||||||||||||||||||||||||||||||||||||
R  ACTACTAGGTCTTCTCTTAGCCTCAGCAGGAAAATCAGCCCAATTTGGACTTCATCCGTGACTTCCATCAGCTTGAAGAAATGCGTGCGAGCTTTAATCAGCACTCCTTCATTCCAGTACAATAGTTATAGCTGGAGTATTTACCCTCATCCGCTTTTATCCATTAATAGAAAACAACCT
   |  |  |  | ||   | |   || | |  || |          |     | ||     |    ||   |||||||||||||||||||||||||||||||||||||    |      |       || |      |  |         ||    |  |      |   |||          ||  |
P  AAAAAAATCTGTTAAATGAATTTCGCGAAAAACAGTCCAAACTTAGAAATTTACAACAGTACCAACATCAGCTTGAAGAAATGCGTGCGAGCTTTAATCAGCAGCTCACACAACTTCCCAAAGAAAGCGAAATTCCTGGATTGGTCGAAGATATTAATATGACAGGCGTAAATTCTGGTC
Pos                                                                    <-3265486--------------------3265449->
R translated                                                      ...S  I  S  i  k  k  c  v  r  a  l  I  S  T...
```

$\Delta G^0_{min}$ = -9.49 kcal/mol + -7.78 kcal/mol  (average: -8.64 kcal/mol)

**MK111.**

```
                                                        |-- net 18 bp lost -----------------------------------------|
A  ACTACTAGGTCTTCTCTTAGCCTCAGCAGGAAAATCAGCCCAATTTGGACTTCATCCGTGACTTCCATCAGCTAGTGAAGGCCCTACCCCAGTATCAGCACTCCTTCATTCCAGTACAATAGTTATAGCTGGAGTATTTACCCTCATCCGCTTTTATCCATTAATAGAAAACAACCTCAC
   |||||||||||||||||||||||||||||||||||||||||||||||| |||| |||  | | | || |  | ||   ||||  |  ||  | |  | || |     |||||||||||||||||||||||||||||||||||||||||||||||||||||||||||||||||||||||||||||||||
R  ACTACTAGGTCTTCTCTTAGCCTCAGCAGGAAAATCAGCCCAATTTGGATTTCACCCG--A-TACGATTA--T-GTC--GGCCG-ATGCC--T-TG-GAACAAC-----TCCAGTACAATAGTTATAGCTGGAGTATTTACCCTCATCCGCTTTTATCCATTAATAGAAAACAACCTCAC
   |          |      | ||        ||    || || |  |||||||||||| | |||||||| | |||  ||||| ||||| | || |||||||     |||||||||     |        |         ||   | |     |           ||   |    |||   |
P  AAACAGGCATGGCAAAGCACGCTGCAACTTGAAGCACGCACACTCGGGATTTCACCCG--A-TACGATTA--T-GTC--GGCCG-ATGCC--T-TG-GAACAAC-----TCCAGTACACCCTCTGGCAAGACACGCAAAGCCTCAATACTGGTAATGGAGCAAACTAATGTTAACAAAAA
Pos                                               <-3288189-------------------------------------------------------3288136->
R translated                                   ...W  i  s  p  d     t  I  m     s     a  d   a     l     e  q  l       Q  Y  N...
```

$\Delta G^0_{min}$ = -11.67 kcal/mol + -11.22 kcal/mol  (average: -11.45 kcal/mol)

**MK43.**

```
                                                                       |-- net 6 bp loss ------|
A  ACTACTAGGTCTTCTCTTAGCCTCAGCAGGAAAATCAGCCCAATTTGGACTTCATCCGTGACTTCCATCAGCTAGTGAAGGCCCTACCCCAGTATCAGCACTCCTTCATTCCAGTACAATAGTTATAGCTGGAGTATTTACCCTCATCCGCTTTTATCCATTAATAGAAAACAACCTCAC
   ||||||||||||||||||||||||||||||||||||||||||||||||||||||||||||||||||||||| || |  ||   | | ||||||| |||||||||||||||||||||||||||||||||||||||||||||||||||||||||||||||||||||||||||||||||||||||||
R  ACTACTAGGTCTTCTCTTAGCCTCAGCAGGAAAATCAGCCCAATTTGGACTTCATCCGTGACTTCCATCAAC-AGCG-CGG--C-A-CCCAGTACCAGCACTCCTTCATTCCAGTACAATAGTTATAGCTGGAGTATTTACCCTCATCCGCTTTTATCCATTAATAGAAAACAACCTCAC
       |          | |  ||| ||      |            | |  |||| | |||||| |||| |||  | | ||||||||| ||              |  ||    ||     |        |     |       ||                || || ||    |
P  GTCGATTCAGACAACAACACACCCGCCAGAAATGCGCCCATCGCCATCGACAAACCACTGACCTGCATCAAC-AGCG-CGG--C-A-CCCAGTACCACCAGCAAGGCTGCTGCGGTCATCACTTCACGCGCTTTGGATGCGGCCAGTAATCTAAACAGTGGATTAAGCAACCAGAAACCT
Pos                                                          <-3360211-----------------------3360246->
R translated                                           ...V  T  S  I  n  s  a  a       P  S  t  S  T...
```

$\Delta G^0_{min}$ = -25.91 kcal/mol

```
MK120.                                                                     |-- net 6 bp loss  ---|
A  ACTACTAGGTCTTCTCTTAGCCTCAGCAGGAAAATCAGCCCAATTTGGACTTCATCCGTGACTTCCATCAGCTAGTGAAGGCCCTACCCCAGTATCAGCACTCCTTCATTCCAGTACAATAGTTATAGCTGGAGTATTTACCCTCATCCGCTTTTATCCATTAATAGAAAACAACCTCAC
   |||||||||||||||||||||||||||||||||||||||||||||||||||||||||||||||||||||||| | | ||    ||   |||  | ||||||||||||||||||||||||||||||||||||||||||||||||||||||||||||||||||||||||||||||||||||||
R  ACTACTAGGTCTTCTCTTAGCCTCAGCAGGAAAATCAGCCCAATTTGGACTTCATCCGTGACTTCCATCAGC-AAT-AA---TCT-GACCACCACCAGCACTCCTTCATTCCAGTACAATAGTTATAGCTGGAGTATTTACCCTCATCCGCTTTTATCCATTAATAGAAAACAACCTCAC
        | ||  ||     |  | ||| | |  ||    ||   |  |       | |  |  | || ||||| ||| ||   ||| ||||||||||||||||  | |  |  |   | |  | | |       ||   ||    |||        |     | |||    |  |   |
P  CAACATCGGCTTTAAGAAACGCCCAGTATGTGAAATTTCCACTTCAGCCACCTGTTCTGGTGTACCTTCAGC-AAT-AA---TCT-GACCACCACCAGCACCACCTTCTGGCCCAAGATCAATCACCCAGTCAGCGGTTTTAATCACATCTAGATTATGCTCAATCACCACAATGGTATT
Pos                                                                <-3379400--------------------3379363->
R translated                                                   ...S  I  S   n   n     l   t  t  t  S  T...
```

**$\Delta G^0_{min}$ = -6.30 kcal/mol + -10.13 kcal/mol (average: -8.22 kcal/mol)**

```
MK137.                  |-- net 123 bp loss -----------------------------------------------------------------------------------------------------------------------|
A  ACTTCATCCGTGACTTCCATCAGCTAGTGAAGGCCCTACCCCAGTATCAGCACTCCTTCATTCCAGTACAATAGTTATAGCTGGAGTATTTACCCTCATCCGCTTTTATCCATTAATAGAAAACAACCTCACTATTCAAACTTCAACACTTCTGGTTCTGGCTCTAATGTCACTAATGAA
   ||||||||||||||||||||| ||                                                                                                                           | |||||    |  |||||||||||||||||||
R  ACTTCATCCGTGACTTCCATCGGC---------------------------------------------------------------------------------------------------------------------------AGTTCTGTCAATTCCTCTAATGTCACTAATGAA
         ||   ||   ||||||||                                                                                                                           ||||||||||||||||||||||||||
P  GGCATGTCTAAGAGCACCATCGGC---------------------------------------------------------------------------------------------------------------------------AGTTCTGTCAATTCCTCTAATGTCACAGCACTG
Pos                <-3417866--------------------------------------------------------------------------------------------------------------------------------------------3417899->
R translated   ...S  I  g                                                                                                                             s  s  v  n  s  S  N  V  T...
```

**$\Delta G^0_{min}$ = -7.88 kcal/mol + -14.83 kcal/mol (average: -11.36 kcal/mol)**

```
MK122.                                                                    |--| no net loss/gain
A  TACTAGGTCTTCTCTTAGCCTCAGCAGGAAAATCAGCCCAATTTGGACTTCATCCGTGACTTCCAT-CAGCTAGTGAAGGCCCTACCCCAGTATCAGCACTCCTTCATTCCAGTACAATAGTTATAGCTGGAGTATTTACCCTCATCCGCTTTTATCCATTAATAGAAAACAACCTCAC
   ||||||||||||||||||||||||||||||||||||||||||||||||||||||||||||||||||||||| |  ||||||||||||||||||||||||||||||||||||||||||||||||||||||||||||||||||||||||||||||||||||||||||||||||||||||||
R  TACTAGGTCTTCTCTTAGCCTCAGCAGGAAAATCAGCCCAATTTGGACTTCATCCGTGACTTCCAT-CAGCCATAGAAGGCCCTACCCCAGTATCAGCACTCCTTCATTCCAGTACAATAGTTATAGCTGGAGTATTTACCCTCATCCGCTTTTATCCATTAATAGAAAACAACCTCAC
   |   |          |        |   | |||      |||    |     |    |  ||| || |||||||||||                ||| |       ||           |   | |  |  |  |   ||     |      |  |      |  ||  || | ||
P  TTGAACACGACGAACTGATTAAGCCTTTAGAATTTTTAAAATCAAAAGGAATTGAAAGTATTC-ATGCAGCCATAGAACAAAAAGAAGTTCAGACAGTAAAAGGCGATCAGGAGTTTGGACCAAAATATACACCACAAACTTCGCTAGCACATGTTGATCCGAACGACTACGATCTTCT
Pos                                                     3558812<---------------->3558828
R translated                                              ...T  S  I   S  h  I  R...
```

**$\Delta G^0_{min}$ = -6.37 kcal/mol**

II. chromosomal templating DNA (*rrn* operons)

**R1517 (*rrn* operon).**

```
                                                                         |----| no net gain/loss
A   ACTACTAGGTCTTCTCTTAGCCTCAGCAGGAAAATCAGCCCAATTTGGACTTCATCCGTGACTTCCATCAGCTAGTGAAGGCCCTACCCCAGTATCAGCACTCCTTCATTCCAGTACAATAGTTATAGCTGGAGTATTTACCCTCATCCGCTTTTATCCATTAATAGAAAACA
    ||||||||||||||||||||||||||||||||||||||||||||||||||||||||||||||||||||||||| |||  ||||||||||||||||||||||||||||||||||||||||||||||||||||||||||||||||||||||||||||||||||||||||||||||
R   ACTACTAGGTCTTCTCTTAGCCTCAGCAGGAAAATCAGCCCAATTTGGACTTCATCCGTGACTTCCATCAGCTTGTGCGGGCCCTACCCCAGTATCAGCACTCCTTCATTCCAGTACAATAGTTATAGCTGGAGTATTTACCCTCATCCGCTTTTATCCATTAATAGAAAACA

P   GAAAGTTTCTACTATGTCAAGGCCAGGTAAGGTTCTTCGCGTTGCATCGAATTAAACCACATGCTCCACCGCTTGTGCGGGCCCCCGTCAATTCATTTGAGTTTTAGTCTTGCGACCGTACTCCCCAGGCGGTCTACTTATCGCGTTAGCTGCGCCACTAAAGCCTCAAAGGC
Pos                                                                 19344<------------>19331
                                                                   214129<------------>214116
                                                                   646966<------------>646953
                                                                  1661628<------------>1661615
                                                                  2946435<------------>2946448
                                                                  3075532<------------>3075545
                                                                  3563854<------------>3563867
R translated                                                           ...S  c  c  g  P...
```

**$\Delta G^0_{min}$ = -13.81 kcal/mol**

**MK147 (*rrn* operon).**

```
                                                                        |---| no net gain/loss
A   ACTACTAGGTCTTCTCTTAGCCTCAGCAGGAAAATCAGCCCAATTTGGACTTCATCCGTGACTTCCATCAGCTAGTGAAGGCCCTACCCCAGTATCAGCACTCCTTCATTCCAGTACAATAGTTATAGCTGGAGTATTTACCCTCATCCGCTTTTATCCATTAATAGAAAACAACCTCAC
    |||||||||||||||||||||||||||||||||||||||||||||||||||||||||||||||||||||||| ||  |||||||||||||||||||||||||||||||||||||||||||||||||||||||||||||||||||||||||||||||||||||||||||||||||||||||||||||
R   ACTACTAGGTCTTCTCTTAGCCTCAGCAGGAAAATCAGCCCAATTTGGACTTCATCCGTGACTTCCATCAGCCAGCTAAGGCCCTACCCCAGTATCAGCACTCCTTCATTCCAGTACAATAGTTATAGCTGGAGTATTTACCCTCATCCGCTTTTATCCATTAATAGAAAACAACCTCAC

P   CTATGAGTACTATCCGGGAGACAGACTGCGGGTGCTAACGTCCGTAGTCAAGAGGAAAACAATCCAGACCGCCAGCTAAGGCCCCAAAATCATAGTTAAGTGGGAAACGATGTGGGAAGGCATAGACAGCTAGGAGGTTGGCTTAGAAGCAGCCACCCTTTAAAGAAAGCGTAATAGCTC
Pos                                                               21516<-------------->21537
                                                                 216301<-------------->216322
                                                                 649228<-------------->649249
                                                                1663800<-------------->1663821
                                                                2944173<-------------->2944152
                                                                3073270<-------------->3073249
                                                                3561682<-------------->3561661
R translated                                                         ...I  S  q  l  R  P...
```

**$\Delta G^0_{min}$ = -14.64 kcal/mol**

**MK118 (*rrn* operon).**

```
                                  |-- net 15 bp  loss ------------------------------|
A   ACTACTAGGTCTTCTCTTAGCCTCAGCAGGAAAATCAGCCCAATTTGGACTTCATCCGTGACTTCCATCAGCTAGTGAAGGCCCTACCCCAGTATCAGCACTCCTTCATTCCAGTACAATAGTTATAGCTGGAGTATTTACCCTCATCCGCTTTTATCCATTAATAGAAAACAACCTCAC
    |||||||||||||||||||||||||||||    | |  | ||| |    |||||| || | |||  | |||  |  |    |||||||||||||||||||||||||||||||||||||||||||||||||||||||||||||||||||||||||||||||||||||||||||||||||||
R   ACTACTAGGTCTTCTCTTAGCCTCAGCAG----A-CTTCTCAA-TCCACCTTCAT-CG-G-CTT--A-CAG--AACG-CTCCCCTACCCCAGTATCAGCACTCCTTCATTCCAGTACAATAGTTATAGCTGGAGTATTTACCCTCATCCGCTTTTATCCATTAATAGAAAACAACCTCAC

P   AGCATTCGCACTTCTGATACCTCCAGCAG----A-CTTCTCAA-TCCACCTTCAT-CG-G-CTT--A-CAG--AACG-CTCCCCTACCACGTAACTTAAAGTTACATCCGCAGCTTCGGCACATAGTTTTAGCCCCGTTACATCTTCCGCGCAGGCCGACTCGACTAGTGAGCTATTACG
Pos                       <-21761----------------------------------------------------21710->
                          <-216546--------------------------------------------------216495->
                          <-649473--------------------------------------------------649422->
                          <-1664045------------------------------------------------1663994->
                          <-2943928------------------------------------------------2943979->
                          <-3073025------------------------------------------------3073076->
                          <-3561437------------------------------------------------3561488->
R translated         ...L  S  r     l  l  n   p  p  s  s   a  y    I   t  l  P  y  P...
```

**$\Delta G^0_{min}$ = -6.63 kcal/mol + -9.61 kcal/mol  (average: -8.12 kcal/mol)**

III. plasmid templating DNA

```
MK126 (pK71-77-1-NDM).                                                                    |----------------| no net loss/gain
A  ACTACTAGGTCTTCTCTTAGCCTCAGCAGG-AAAATCAGCCCAATTTGGACTTCAT-CCGTGACTTCCATCAGCTAG-T-GAAGGCCCTACCCCAGTATCAGCACTCCTTCA-TTCCAGTACAATAGTTATAGCTGGAGTATTTACCCTCATCCGCTTTTATCCATTAATAGAAAACAACCTCAC
R  ACTACTAGGTCTTCTCTTAGCCTCAGCAGGAAAAATCAGCCCAATTTGGACTTCATCCCGTGACTTCCATCTGCTCGCTTGGA-TCAA-ACCCCAGTATCAGCACTCCTTCATTTCCAGTACAATAGTTATAGCTGGAGTATTTACCCTCATCCGCTTTTATCCATTAATAGAAAACAACCTCAC
pK TTTATGATACTGTAAACAATAAAAAGACACCTTGCAAGACTTTCACATGTGAAAGTTTTCTGGTTTACATCTGCTCGCTTGGA-TCAA-ACCCCAGAGCCTTGCACACGTCATCCAGGTCATCGAGCGTGAAGAATCCCAGCGTCAGCTCCCCGGAGCGCTTCGCGGGGTTATAGCGAATTTTGA
Pos                                                                 <-132434----------------132402->
R translated                                                   ...T  S  I  c  s  l  g   s  n  P  S...
ΔG0min = -13.07 kcal/mol

MK52 (pQLICE).                                                        |-- net 3 bp loss -|
A  ACTACTAGGTCTTCTCTTAGCCTCAGCAGGAAAATCAGCCCAATTTGGACTTCATCCGTGACTTCCATCAGCTAGTG-AAGGCCCTACCCCAGTATCAGCACTCCTTCATTCCAGTACAATAGTTATAGCTGGAGTATTTACCCTCATCCGCTTTTATCCATTAATAGAAAACAACCTCAC
R  ACTACTAGGTCTTCTCTTAGCCTCAGCAGGAAAATCAGCCCAATTTGGACTTCATCCGCG--TTCCA--AGCTGCGGAAAGGCCCTACCCCAGTATCAGCACTCCTTCATTCCAGTACAATAGTTATAGCTGGAGTATTTACCCTCATCCGCTTTTATCCATTAATAGAAAACAACCTCAC
pQ CGCACCTGCTTGATCGCGGCCGCGATAGCTAGATCGCGTTGCTCCTCTTCTCCATCCGCG--TTCCA--AGCTGCGGAAAGGCACCCATAAGCGTACGCCTGGTCGAGCAGGCGACGCGGATCGACGTCCAGCGCACGAGAGAATGCGTCCGCCATCTGTGCAATGCGTCTAGGATCGAGA
Pos                                                      <-1637------------------------1606->
R translated                                         ...T  S  S  a    f  q    a  a  e  R  P...
ΔG0min = -23.13 kcal/mol

MK50 (pQLICE).                                                                                 |----| no net loss/gain
A  ACTACTAGGTCTTCTCTTAGCCTCAGCAGGAAAATCAGCCCAATTTGGACTTCATCCGTGACTTCCATCAGCTAGTGAAGGC--CCTACCCCAGTATCAGCACTCCTTCATTCCAGTACAATAGTTATAGCTGGAGTATTTACCCTCATCCGCTTTTATCCATTAATAGAAAACAACCTC
R  ACTACTAGGTCTTCTCTTAGCCTCAGCAGGAAAATCAGCCCAATTTGGACTTCATCCGTGACTTCCATCAGCTTCCGATGGC--CCTACCCCAGTATCAGCACTCCTTCATTCCAGTACAATAGTTATAGCTGGAGTATTTACCCTCATCCGCTTTTATCCATTAATAGAAAACAACCTC
pQ TCCAGAACGGGAGTGCGCCTTGAGCGACACGAATTATGCAGTGATTTACGACCTGCACAGCCATACCACAGCTTCCGATGGCTGCCTGACGCCAGAAGCATTGGTGCACCGTGCAGTCGATAAGCTCCGCCGGATCAGCTTGCAATTCGCGCGCGAAGGCGAAGCGGCATTTACGTTGAC
Pos                                                                          <-2251--2269->
R translated                                                           ...I  S  f  r  w  P    Y...
ΔG0min = -5.41 kcal/mol

MK62 (pQLICE).                                                                                   |------| no net loss/gain
A  ACTACTAGGTCTTCTCTTAGCCTCAGCAGGAAAATCAGCCCAATTTGGACTTCATCCGTGACTTCCATCAGCTAGTGAAGGCCCTACCCCAGTATCAGCACTCCTTCATTCCAGTACAATAGTTATAGCTGGAGTATTTACCCTCATCCGCTTTTATCCATTAATAGAAAACAACCTCAC
R  ACTACTAGGTCTTCTCTTAGCCTCAGCAGGAAAATCAGCCCAATTTGGACTTCATCCGTGACTTCCATCAGCTATTTAACGACCTACCCCAGTATCAGCACTCCTTCATTCCAGTACAATAGTTATAGCTGGAGTATTTACCCTCATCCGCTTTTATCCATTAATAGAAAACAACCTCAC
pQ AGGGAAAGGAATTGCCGGGCTATGTGCAACGGGAATTTGAAGACCGGTAAACCAGCAATAGACATAAGCGGCTATTTAACGACCCTGCCCTGAACCGACGACCGGGTCGAATTTGCTTTCGAATTTCTGCCATTCATCCGCTTATTATCACTTATTCAGGCGTAGCAACCAGGCGTTTAA
Pos                                                                          <-3871----3856->
R translated                                                              ...S  y  l  t  t  Y
ΔG0min = -9.24 kcal/mol

MK131 (pQLICE).                                                                               |-- net 24 bp loss ---------------------|
A  ACTACTAGGTCTTCTCTTAGCCTCAGCAGGAAAATCAGCCCAATTTGGACTTCATCCGTGACTTCCATCAGCTAGTGAAGGCCCTACCCCAGTATCAGCACTCCTTCATTCCAGTACAATAGTTATAGCTGGAGTATTTACCCTCATCCGCTTTTATCCATTAATAGAAAACAACCTCAC
R  ACTACTAGGTCTTCTCTTAGCCTCAGCAGGAAAATCAGCCCAATTTGGACTTCATCCGTGACTTCCATCAGCCAGTG------------------------CCCGTCACTCCCGTACAATAGTTATAGCTGGAGTATTTACCCTCATCCGCTTTTATCCATTAATAGAAAACAACCTCAC
pQ TACAGTCTATCGCTTAGCGGAAAGTTCTTTTACCCTCAGCCGAAATGCCTGCCGTTGCTAGACATTGCCAGCCAGTG------------------------CCCGTCACTCCCGTACTAACTGTCACGAACCCCTGCAATAACTGTCACGCCCCCCTGCAATAACTGTCACGAACCCCTG
Pos                                                                          <-4174------------------------------------4198->
R translated                                                                I  S  i  c                        P  S  l  p  Z
ΔG0min = -8.73 kcal/mol + -13.60 kcal/mol  (average: -11.17 kcal/mol)

MK80 (pQLICE).                                                                     |-----------------| no net gain/loss
A  ACTACTAGGTCTTCTCTTAGCCTCAGCAGGAAAATCAGCCCAATTTGGACTTCATCCGTGACTTCCATCAGCTAGTGAAGGCCCTACCCCAGTATCAGCACTCCTTCATTCCAGTACAATAGTTATAGCTGGAGTATTTACCCTCATCCGCTTTTATCCATTAATAGAAAACAACCTCAC
R  ACTACTAGGTCTTCTCTTAGCCTCAGCAGGAAAATCAGCCCAATTTGGACTTCATCCGTGACTTCCAACACCCCGCCAGCCCCCGCCCCCAGTATCAGCACTCCTTCATTCCAGTACAATAGTTATAGCTGGAGTATTTACCCTCATCCGCTTTTATCCATTAATAGAAAACAACCTCAC
pQ CCCCACAAGGGCGCTGATAACCGCGCCTAGTGGATTATTCTTAGATAATCATGGATGGATTTTTCCAACACCCCGCCAGCCCCCGCCCCTGCTGGGTTTGCAGGTTTGGGGGCGTGACAGTTATTGCAGGGGTTCGTGACAGTTATTGCAGGGGGGCGTGACAGTTATTGCAGGGGTTCG
Pos                                                                    <-4325---------------4299->
R translated                                                     ...T  S  n  t  p  p  a  p  a  P...
ΔG0min = -14.01 kcal/mol
```

**MK132 (pQLICE).**

```
                                       |-- net 54 bp loss -------------------------------------------------------------------|
A  ACTACTAGGTCTTCTCTTAGCCTCAGCAGGAAAATCAGCCCAATTTGGACTTCATCCGTGACTTCCATCAGCTAGTGAAGGCCCTACCCCAGTATCAGCACTCCTTCATTCCAGTACAATAGTTATA-GCTGGAGTATTTACCCTCATCCGCTTTTATCCATTAATAGAAAACAACCTCAC
R  ACTACTAGGTCTTCTCTTAGCCTCAGCAGGAAAATCAGCCCGCTAAACCCCACACCAA-------------------------------------------------------ACCCCGCAGAAATACGCTGGAGTATTTACCCTCATCCGCTTTTATCCATTAATAGAAAACAACCTCAC
pQ GCCCCACCGCTGCGCGGCAGGGGGAAAGGCGGGCAAAGCCCGCTAAACCCCACACCAA-------------------------------------------------------ACCCCGCAGAAATACGCTGGAGCGCTTTTAGCCGCTTTAGCGGCCTTTCCCCCTACCCGAAGGGTGGG
Pos                                   <-4498---------------------------------------------------------------------------------------4541->
R translated                       ...S  P  l  n  p  t  p  n                                                     p  a  e  i  r  W  S...
```

$\Delta G^0_{min}$ = -5.90 kcal/mol + -10.00 kcal/mol (average: -7.95 kcal/mol)

**MK47 (pQLICE).**

```
                                                                         |----| no net loss/gain
A  ACTACTAGGTCTTCTCTTAGCCTCAGCAGGAAAATCAGCCCAATTTGGACTTCATCCGTGACTTCCATCAGCTAGTGAAGGCCCTACCCCAGTATCAGCACTCCTTCATTCCAGTACAATAGTTATAGCTGGAGTATTTACCCTCATCCGCTTTTATCCATTAATAGAAAACAACCTCAC
R  ACTACTAGGTCTTCTCTTAGCCTCAGCAGGAAAATCAGCCCAATTTGGACTTCATCCGTGACTTCCATCAGCTGGTGGGGGCCCTACCCCAGTATCAGCACTCCTTCATTCCAGTACAATAGTTATAGCTGGAGTATTTACCCTCATCCGCTTTTATCCATTAATAGAAAACAACCTCAC
pQ ATGCCGAAATTCAGCGGGTGCGGGCAAGGGAACAGCAGCAAGAGCGCAAGAACGAAACAAGGCGCAAGGTGCTGGTGGGGGCCATGATTTTGGCCAAGGTGAACAGCAGCGAGTGGCCGGAGGATCGGCTCATGGCGGCAATGGATGCGTACCTTGAACGCGACCACGACCGCGCCTTGT
Pos                                                                4776<-------------->4762
R translated                                                       ...S  w  w  g  P...
```

$\Delta G^0_{min}$ = -12.37 kcal/mol

**MK83 (pQLICE).**

```
                                                                     | net 3 bp loss |
A  ACTACTAGGTCTTCTCTTAGCCTCAGCAGGAAAATCAGCCCAATTTGGACTTCATCCGTGACTTCCATCAGCTAGTGAAGGCCCTACCCCAGTATCAGCACTCCTTCATTCCAGTACAATAGTTATAGCTGGAGTATTTACCCTCATCCGCTTTTATCCATTAATAGAAAACAACCTCAC
R  ACTACTAGGTCTTCTCTTAGCCTCAGCAGGAAAATCAGCCCAATTTGGACTTCATCCGTGACTTCCATC-TCCA--CCACGTTCGGCCCCAGTATCAGCACTCCTTCATTCCAGTACAATAGTTATAGCTGGAGTATTTACCCTCATCCGCTTTTATCCATTAATAGAAAACAACCTCAC
pQ TGGGCGTTGGCGGTGTCGATGTTCAGGGCCACGTCTGCCCGGTCGGTGCGGATGCCCCGGCCTTCCATC-TCCA--CCACGTTCGGCCCCAGGTGAACACCGGGCAGGCGCTCGATGCCCTGCGCCTCAAGTGTTCTGTGGTCAATGCGGGCGTCGTGGCCAGCCCGCTCTAATGCCCGG
Pos                                                          <-5766-------------------5733->
R translated                                           ...S  V  T  S  I  s  t   t  f  g  P  S...
```

$\Delta G^0_{min}$ = -13.21 kcal/mol + -7.64 kcal/mol (average: -10.43 kcal/mol)

**MK53 (pQLICE).**

```
                                                                    |---------| no net gain/loss
A  ACTACTAGGTCTTCTCTTAGCCTCAGCAGGAAAATCAGCCCAATTTGGACTTCATCCGTGACTTCCATCAGCTAGTGAAGGCCCTACCCCAGTATCAGCACTCCTTCATTCCAGTACAATAGTTATAGCTGGAGTATTTACCCTCATCCGCTTTTATCCATTAATAGAAAACAACCTCAC
R  ACTACTAGGTCTTCTCTTAGCCTCAGCAGGAAAATCAGCCCAATTTGGACTTCATCCGTGACTTCCAGCAGATCGAGCAGGCCCTACCCCAGTATCAGCACTCCTTCATTCCAGTACAATAGTTATAGCTGGAGTATTTACCCTCATCCGCTTTTATCCATTAATAGAAAACAACCTCAC
pQ TGCTGCTGCGTGAATCCAAGGGCAAGACCGCCACCGCTGGCCCGGCGCTGGTGCAGCAGGCTGGCCAGCAGATCGAGCAGGCCCAGCGGCAGCAGGAGAAGGCCCGCAGGCTGGCCAGCCTCGAACTGCCCGAGCGGCAGCTTAGCCGCCACCGGCGCACGGCGCTGGACGAGTACCGCA
Pos                                                               <-6830--------6849->
R translated                                                   ...S  s  r  s  s  R  P...
```

$\Delta G^0_{min}$ = -17.77 kcal/mol

**MK56 (pQLICE).**

```
                                                                         |--| no net gain/loss
A  ACTACTAGGTCTTCTCTTAGCCTCAGCAGGAAAATCAGCCCAATTTGGACTTCATCCGTGACTTCCATCAGCTAGTGAAGGCCCTACCCCAGTATCAGCACTCCTTCATTCCAGTACAATAGTTATAGCTGGAGTATTTACCCTCATCCGCTTTTATCCATTAATAGAAAACAACCTCAC
R  ACTACTAGGTCTTCTCTTAGCCTCAGCAGGAAAATCAGCCCAATTTGGACTTCATCCGTGACTTCCATCAGCAGGAGAAGGCCCTACCCCAGTATCAGCACTCCTTCATTCCAGTACAATAGTTATAGCTGGAGTATTTACCCTCATCCGCTTTTATCCATTAATAGAAAACAACCTCAC
pQ GCAAGACCGCCACCGCTGGCCCGGCGCTGGTGCAGCAGGCTGGCCAGCAGATCGAGCAGGCCCAGCGGCAGCAGGAGAAGGCCCGCAGGCTGGCCAGCCTCGAACTGCCCGAGCGGCAGCTTAGCCGCCACCGGCGCACGGCGCTGGACGAGTACCGCAGCGAGATGGCCGGGCTGGTCA
Pos                                                                       <-6855----6870->
R translated                                                         ...I  S  r  r  R  P...
```

$\Delta G^0_{min}$ = -15.50 kcal/mol

**MK45 (pQLICE).**

```
                                                                      |--------| no net gain/loss
A  ACTACTAGGTCTTCTCTTAGCCTCAGCAGGAAAATCAGCCCAATTTGGACTTCATCCGTGACTTCCATCAGCTAGTGAAGGCCCTACCCCAGTATCAGCACTCCTTCATTCCAGTACAATAGTTATAGCTGGAGTATTTACCCTCATCCGCTTTTATCCATTAATAGAAAACAACCTCAC
R  ACTACTAGGTCTTCTCTTAGCCTCAGCAGGAAAATCAGCCCAATTTGGACTTCATCCGTGACTTCCATCACCGCCTGCACGCCCTACCCCAGTATCAGCACTCCTTCATTCCAGTACAATAGTTATAGCTGGAGTATTTACCCTCATCCGCTTTTATCCATTAATAGAAAACAACCTCAC
pQ AGGTGGGCGAACTGCCCACCGGCCCGGTGATCTACCTGCCCGCCGAAGACCCGCCCACCGCCATTCATCACCGCCTGCACGCCCTTGGGGCGCACCTCAGCGCCGAGGAACGGCAAGCCGTGGCTGACGGCCTGCTGATCCAGCCGCTGATCGGCAGCCTGCCCAACATCATGGCCCCGG
Pos                                                                  <-8001--------8020->
R translated                                                    ...S  I  t  a  c  t  P  S...
```

$\Delta G^0_{min}$ = -15.31 kcal/mol

**MK55 (pQLICE).**

```
                                                                        |-----------|  net 3 bp loss
A  ACTACTAGGTCTTCTCTTAGCCTCAGCAGGAAAATCAGCCCAATTTGGACTTCCATCAGCTAGTGAAGGCCCTACCCCAGTATCAGCACTCCTTCATTCCAGTACAATAGTTATAGCTGGAGTATTTACCCTCATCCGCTTTTATCCATTAATAGAAAACAACCTCAC
   |||||||||||||||||||||||||||||||||||||||||||||||||||||||||||||||||||||  || | | ||  ||||||||||||||||||||||||||||||||||||||||||||||||||||||||||||||||||||||||||||||||||||||||||||||||||
R  ACTACTAGGTCTTCTCTTAGCCTCAGCAGGAAAATCAGCCCAATTTGGACTTCCATCCGCT-G-GCAGT-CCTACCCCAGTATCAGCACTCCTTCATTCCAGTACAATAGTTATAGCTGGAGTATTTACCCTCATCCGCTTTTATCCATTAATAGAAAACAACCTCAC
    |  | | |          ||  |     | |   |||||      |  |   |   ||   |  |||||||||| | ||||| ||||||     | ||   |              |  |  |   |               |  |      |  |     |       |   |  |  ||
pQ GCGGCCATGATGGGCGCAGGCGACCAGCAGCAGGCCAGCCGGGGCAGCTCGGTACTGGTCGATAACATCCGCT-G-GCAGT-CCTACCTGTCGAGCATGACCAGCGCCGAGGCCGAGGAATGGGGTGTGGACGACGACCAGCGCCGGTTCTTCGTCCGCTTCGGTGTGAGCAAGGCCAAC
Pos                                                                 <-8366-----------8385->
R translated                                                   ...S  I   r   g   s   P  T...
```

**$\Delta G^0_{min}$ = -13.06 kcal/mol**
