## Supplemental Tables and Figures for "Plasmids modulate microindel mutations in *Acinetobacter baylyi* ADP1"

Liljegren et al.

### Supplemental Material

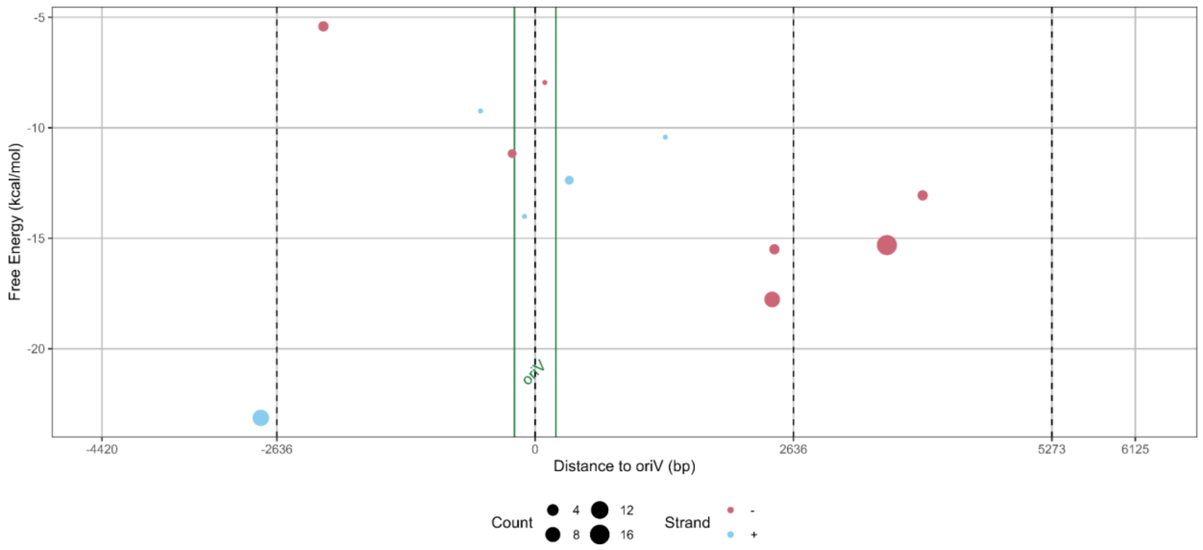


**Suppl. Fig. S1.** Characteristics of SPDIR events generated with DNA from pQLICE-derived vectors. The horizontal axis represents the location of templating DNA, with vertical green lines delimiting the origin of replication and vertical dashed black lines dividing the plasmid sequence in quartiles. The vertical axis represents the Free Energy of Hybridization. Circle color and size represent the donor strand (same as in Fig. 5B) and the number of observations, respectively.

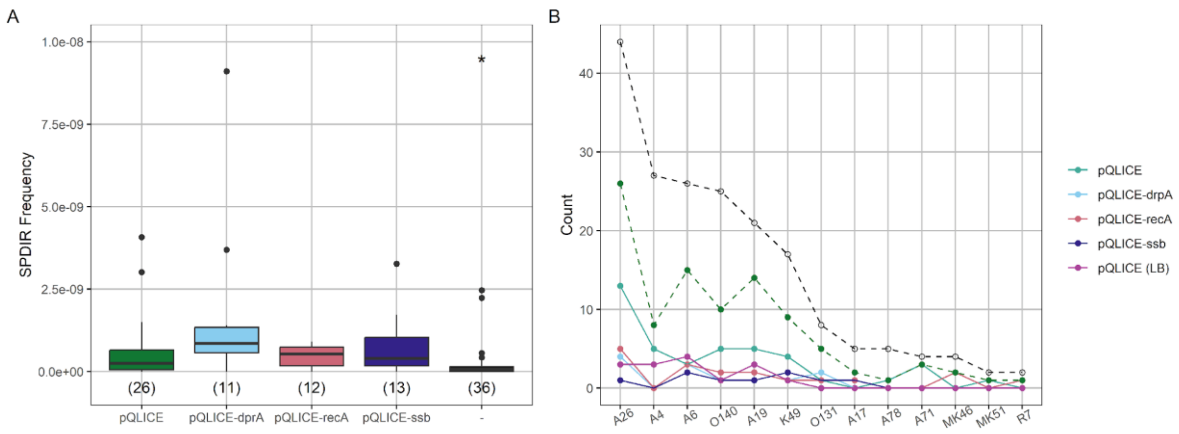


**Suppl. Fig. S2.** Effect of pQLICE-derived vectors on SPDIR in *A. baylyi* Δ*recJ* Δ*exoX*. A: SPDIR frequency determined for experiments where $His^+$ mutants were detected. Number of experiments is indicated in parenthesis for each plasmid; - indicates the plasmid-free strain and pQLICE the empty vector. The significant result of the Kruskal-Wallis rank sum test followed by Dunn many-to-one post hoc test (with pQLICE as reference group) are indicated as * for p = 0.026. B: Number of SPDIR events detected independently of plasmid carriage among experiments with distinct pQLICE-derivatives. The dashed black line indicates the total number of events, and the dashed green line the total number of events with pQLICE-derivatives.

**Supplemental Table S1:** Plasmid stability in *A. baylyi* Δ*recJ* Δ*exoX*.

| Plasmid | Antibiotic selection [mg/L] | Mean plasmid loss without selection | n: |
|---|---|---|---|
| pQLICE | streptomycin 40 | 43.% | 4 |
| pBBR1MCS-3 | tetracycline 10 | 0% | 6 |
| pRK415 | tetracycline 10 | 0.07% | 5 |
| RN3 | tetracycline 10 | 88% | 8 |
| R16a | kanamycin 25 | 0% | 5 |
| R388 | trimethoprim 100 | 0.02% | 8 |
| pK71-77-1-NDM | ampicillin 100 | 0% | 4 |
